# Cryo-EM structures of brain-derived G protein-coupled receptors

**DOI:** 10.64898/2026.04.01.715822

**Authors:** Nicholas J. Wright, Yi-Ting Chiu, Kensuke Sakamoto, D. Dewran Kocak, Pierre Llorach, Blake A. Fordyce, Kunjie Hua, Karen Lu Huang, Grégory Scherrer, Scott P. Lyons, Steven C. Bremmer, Lee Graves, Jessica J. Walsh, Bryan L. Roth

**Affiliations:** Department of Pharmacology, University of North Carolina at Chapel Hill, Chapel Hill, North Carolina, USA; Department of Cell Biology and Physiology, University of North Carolina at Chapel Hill, Chapel Hill, NC USA; UNC Neuroscience Center, University of North Carolina at Chapel Hill, Chapel Hill, NC USA; UNC Metabolomics and Proteomics Core Facility, Department of Pharmacology, The University of North Carolina at Chapel Hill, Chapel Hill, NC, USA

## Abstract

Glutamate is the main excitatory neurotransmitter in the brain and mediates its actions by both ionotropic (e.g. NMDA and AMPA) and metabotropic glutamate receptors (mGluRs). The Groups II and III mGluRs, which pre-synaptically inhibit glutamate release, are important for synaptic plasticity, modulating neuronal excitation, learning and memory. Our current understanding of the structural organization and dynamics of these and other mGluRs, as well as most other GPCRs, relies mainly on studies using recombinant and highly engineered systems *in vitro*. Here, we combine CRISPR-mediated protein tagging, proteomics and a rapid immunoaffinity purification method to isolate endogenous mGluR2-containing assemblies from mouse brain and visualize them via cryo-EM. Analysis of the particle sets reveals the molecular structures of at least 11 distinct endogenous receptor assemblies that span active and inactive states, homomeric and heteromeric dimers, and G protein-coupled and uncoupled species. We find that mGluR2 homodimers and mGluR2/3 heterodimers are the major endogenous mGluR2-containing species present in the brain, with the mGluR2/3 heterodimers detected only in active state complexes, potentially reflecting basal activation of mGluR3 containing dimers by chloride. Reconstructing a comprehensive conformational equilibrium for the brain-isolated receptors in detergent reveals endogenous ternary complexes comprising mGluR2 homodimers and mGluR2/3 heterodimers with a single Gα_oA_ heterotrimer which exhibit significant differences from prior studies with recombinant systems. Our work illuminates the endogenous conformational, proteomic and compositional landscape of the heterogenous mGluR2 complexes in the brain, thereby providing a structural framework for the pathophysiology of psychiatric disorders. This information has the potential to be leveraged for therapeutic targeting of endogenous glutamatergic signaling complexes.

## Introduction

Excitatory synaptic transmission in the brain is mediated principally by the amino acid neurotransmitter L-glutamate. Glutamate excites neurons by activating ionotropic receptors in the msec time scale^1,2^ and additionally modulates neuronal excitability and neurotransmission by activating metabotropic receptors in the sec to min time scale (Figure 1A). In total, there are 8 distinct metabotropic glutamate receptor subtypes (mGluR1-mGluR8) in humans, each with its own unique functional and pharmacological properties^3,4^. Of these, the Group II (mGluR2 and mGluR3) and group III (mGluR4, mGluR6, mGluR7 and mGluR8) mGluRs are mainly localized presynaptically where they function as auto- and heteroreceptors to inhibit neuronal firing and neurotransmitter release. Group II/III mGluRs induce neuronal inhibition by activating heterotrimeric G proteins^4^ whereby the Gα_i/o_ subunits inhibit cAMP formation and the liberated β/g subunits can activate K^+^ channels to hyperpolarize neurons^5,6^, inhibit voltage-gated Ca^2+^ channels^7–9^, and inhibit soluble NSF attachment protein (SNARE) complexes^10^ to impair neurotransmitter vesicle fusion (Figure 1A). Conversely, the Group I (mGluR1 and mGluR5) receptors are typically localized postsynaptically and are excitatory via G_q/11_ activation of phospholipase C (Figure 1A)^4,11,12^.

**Figure 1.**
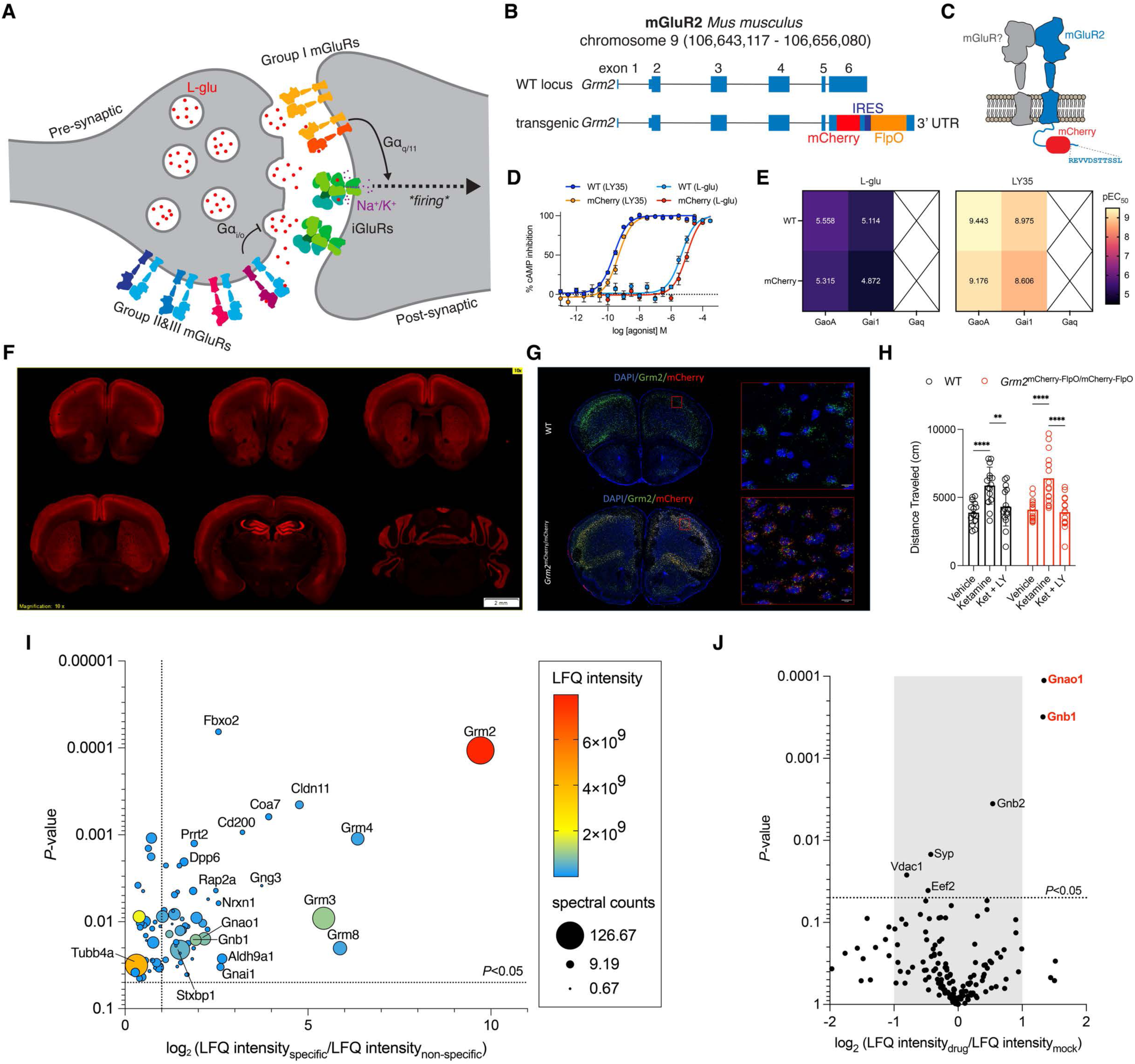
CRISPR/Cas9 engineered transgenic mouse line design and validation. **A**, Cartoon overview of a glutamatergic synapse. Canonical subcellular localization of glutamate receptors shown. **B,** Transgenic mouse line design. Thin boxes represent non-coding exons, thick boxes represent coding exons, and lines between boxes represent introns. Coordinates were drawn from GRCm38/mm10. **C,** Schematic of the mGluR2 gene product in the transgenic line. **D,** mGluR2-mediated glutamate response in HEK293T cells as measured by the GloSensor assay for unmodified and mCherry tagged receptor (n=5 independent experiments, with mean and s.e.m. shown for individual datapoints). **E,** Summary of G-protein selectivity as assessed by TRUPATH. Values shown are pEC50 fits of data shown in Figure S1A. **F,** Distribution of mGluR2-mCherry fusion protein in the brain. Brain sections from the transgenic mouse line were stained with anti-RFP antibody (1:1000) to amplify the mCherry signal and then were acquired with an Olympus slide scanner under a 10X objective. The experiments in this figure were conducted with 3 mice with similar results. **G,** *mCherry* and *Grm2* mRNA expression in mouse brain sections. In situ hybridization experiments were performed to probe *mCherry* and *Grm2* mRNA in wildtype (C57BL/6J) and knockin mice. *Grm2* (green) RNA was detected in brain sections from C57BL/6J; *Grm2* (green) and *mCherry* (red) RNAs were co-localized (yellow) in brain sections from knockin mice (Images were taken using an Olympus VS200 slide scanner with a 10X objective or a Leica confocal microscope under a 40X objective). **H,** Quantification of total distance traveled in the open field test in WT littermate control (*black, n=15*) and mGluR2-mCherry (*red, n=16*) mice following administration of vehicle, ketamine (10mg/kg), or LY354740 (10 mg/kg) pre-treatment 20 minutes prior to ketamine administration (*F2,58*=2.241, *p=*0.1155, *n=*15-16; mean ± s.e.m. **p < 0.01, ****p < 0.001; two-way ANOVA with Tukey’s multiple comparison post hoc test). **I,** LC-MS/MS analysis of RAPID pulldowns from GDN detergent solubilized mouse brain tissue crude homogenate in the absence of drug. LaM6 nanobody construct was used for the specific condition, GFP nanobody construct was used for the non-specific condition (*n*=3 biological replicates). **J**, LC-MS/MS analysis of RAPID pulldowns from GDN detergent solubilized mouse brain tissue crude homogenate, in the presence or absence of 10 μM LY354740 + 10μM JNJ-46281222 (*n*=3 biological replicates).

mGluRs are dimeric class-C G-protein coupled receptors (GPCRs) consisting of three structural domains: the extracellular “Venus-flytrap” domain where glutamate binds (VFT); a seven transmembrane domain (7TM) for transducer coupling; and a cysteine-rich domain (CRD) that connects the VFT to the 7TM. mGluR activation involves coupling of the glutamate-induced VFT closure to transducer binding at the intracellular surface of the 7TM domain – a signal relayed across a distance of approximately 140 Å. Previous structural^13–22^ and biophysical^18,23–26^ studies performed in recombinant systems have delineated the molecular features responsible for mGluR activation and transducer coupling.

As genetic studies have implicated dysregulation of the glutamatergic system in many common neuropsychiatric disorders, including schizophrenia, autism and bipolar disorder^27–31^, mGluRs are increasingly targeted for CNS drug discovery^29,32–35^. Indeed, a recent study demonstrated that an mGluR2 state-selective nanobody can rescue the NMDA receptor hypofunction associated with schizophrenia^36^. Despite this promise, all previously assessed mGluR modulators have failed to show consistent efficacy in clinical trials^34,35,37–41^. A major complication for developing mGluR-preferring medications is the reported presence of multi-subtype heteromeric species^42–45^ and it is well established that recombinant mGluR heterodimers exhibit unique functional and pharmacological properties^16,17,23,46–48^. Our understanding of endogenous, neuronal mGluR assemblies is limited to qualitative imaging and biochemical approaches^42–45^, confirms the presence of homo- and heterodimers but provides limited insight into their molecular organization and assembly in their endogenous environment^46,49,50^. Information regarding the brain distribution of distinct endogenous mGluR assemblies, their molecular structures, and their conformational ensembles— beyond information gained from studies in recombinant expression systems^13,15,45,49,51^—would be a gold standard for structure-based drug design, potentially facilitating therapeutic targeting. Such structural studies have been performed on brain-isolated ionotropic receptors in recent years^52–58^.

To address these fundamental questions, we first devised a method for detergent purifying endogenous mGluR2-containing assemblies from mouse brain tissue, facilitated by CRISPR/Cas9 epitope-tagging of endogenous mGluRs and rapid receptor purification. We chose to start with mGluR2 owning to the fact it is the main autoreceptor for glutamate in the brain^4^, a strong candidate target for neuropsychiatric disorders^59^, exhibits high levels of neuronal expression, and as is mainly localized to presynaptic termini^4^. This contrasts the group II glutamate autoreceptor mGluR3, which exhibits broader cell-type and subcellular expression patterns, including expression in non-neuronal cell populations^4^. Physiological receptor assemblies detergent-purified from whole brains of the reporter line were then quantified by mass spectrometry with 11 distinct assemblies visualized and reconstructed by single-particle cryogenic electron microscopy (cryo-EM) and exhaustive particle analysis. Combining the proteomics and structural information reveals molecular features not previously investigated with recombinant systems including, most notably, endogenous mGluR2 coupling with the endogenous G_oA_ heterotrimer. The various conformational states provide a plausible activation path and subsequent G-protein activation for the endogenous brain receptors. Finally, we find direct structural evidence for differential chloride modulation of receptor homo- and heterodimers, which has potential connections to disease pathophysiology in neuropsychiatric disorders.

## Results

### Generation and characterization of a CRISPR/Cas9 mGluR2-mCherry reporter mouse line

To isolate endogenous mGluR complexes in the brain, we used CRISPR/Cas9 editing to create a mouse line with two modifications at the endogenous *grm2* gene locus: 1) an mCherry tag inserted within the flexible C-terminal tail of the mGluR2 and 2) IRES-FlpO inserted proceeding the 3’ UTR of mGluR2 (Figures 1B-1C). The mCherry tag simultaneously enables visualization of endogenous neuronal mGluR2 complexes by immunohistochemistry and purification of mGluR2 complexes for cryo-EM and proteomics analysis. The FlpO recombinase enables the selective expression of transgenes into mGluR2-expressing cells. Importantly, the translated features of these modifications are installed within the flexible C-terminus of mGluR2 so as not to influence the overall conformational state of the receptor, as previous studies successfully tagged mGluRs in this region^46,48,60^. Notably, the tag was installed in a manner that placed the final 11 amino acids of the receptor C-terminus after the mCherry, to ensure unperturbed subcellular localization of mGluR2 (Figure 1C)^61^. We verified the added mCherry tag did not appreciably affect mGluR2 pharmacology, as reflected by minimal perturbation of agonist (glutamate or LY354740)-mediated inhibition of cAMP compared to an unmodified mGluR2 control (Figure 1D). Furthermore, this modified construct did not display appreciable differences in transducer (Gαi1 or GαoA) coupling preference compared to WT (Figure 1E, Figure S1A). Overall, this design provides an orthogonal approach to the use of conformational-selective mGluR nanobodies that bind to the receptor VFT _domains13,49,51._

To characterize the *Grm2*^mCherry-Flpo^ knockin mice, we first examined whether the distribution pattern of mCherry expression matched that of the endogenous receptor. We first immunolabelled brain sections from *Grm2*^mCherry-Flpo^ using an anti-RFP antibody (Figure 1F). The distribution was consistent with the expression pattern of *Grm2* as assessed by *in situ* hybridization experiments (Figure 1G), and prior studies^62^. Further, *in situ* hybridization revealed that *mCherry* mRNA was detected in the same cells as *Grm2* mRNA in brain sections from knockin mice, but not in brain sections from wildtype mice (C57BL/6J) (Figure 1G). Higher-magnification images of immunolabeled *Grm2*^mCherry-Flpo^ brain sections exhibited a clear subcellular expression pattern for mGluR2, featuring membrane expression^63^. This finding confirms that the C-terminal modification does not hinder proper membrane localization of the receptor.

We then subjected these mice to a behavioral assay to test mGluR2 activity *in vivo. Grm2*^mCherry-Flpo^ and WT littermate control mice were administered 10 mg/kg ketamine after pre-treatment with either saline injection or 10 mg/kg group II mGluR selective agonist LY354740 (LY35)^64^. Agonist pre-treatment ablated the hyper-locomotor activity of ketamine in both WT and *Grm2*^mCherry-Flpo^ mice (Figure 1H), suggesting the CRISPR modification to mGluR2 does not perturb it’s *in vivo* function. To validate FlpO recombinase activity, we crossed *Grm2*^mCherry-FlpO^ mice with reporter mice in which eGFP is expressed in a FlpO-dependent manner (RCE [R26R CAG-boosted EGFP]:FRT reporter mice) (Figure S1B). The distribution of the FlpO-activity-induced eGFP signal was consistent with that of mGluR2-mCherry distributions. Collectively, these data indicate that the insertion of mCherry-FlpO at the *Grm2* locus preserves normal pharmacology, receptor distribution and behavioral function. Further, the FlpO element provides genetic access to mGluR2-expressing neurons, enabling future chemo- and optogenetic studies of mGluR2 function at the circuit level.

### Endogenous neuronal mGluR2 complexes revealed

To enable facile receptor isolation from brain and subsequent downstream analysis, we modified a recently reported nanobody-based purification construct^65^ and refer to this modified protocol within our group as RAPID (**<u>R</u>**eceptor **<u>A</u>**ffinity **<u>P</u>**urification **I**n a **<u>D</u>**ay; Figure S2). We next utilized a high-affinity mCherry nanobody^66^ in the RAPID system to isolate endogenous mGluR2 complexes from glyco-diosgenin (GDN) detergent-solubilized whole-brain homogenates at a small scale (Figure 1I). GDN detergent contrasts with conventional maltoside detergents as it more seectively solubilizes cholesterol-rich plasma membranes, enabling analysis of neuronal receptors present at the presynaptic membrane. MS analysis of the purified fractions revealed that mGluR2 was most highly enriched over the non-specific control and was in high abundance as assessed by label-free quantification (LFQ) and peptide spectrum matches (PSM). Additional significant proteins identified include other endogenous mGluRs – especially mGluR3, and in relatively lower abundance, the group III members mGluR4 and mGluR8 (Figure 1I). Notable mGluR2-associated proteins significantly enriched include the endogenous G proteins Gγ3, Gα_oA_, Gβ1, and Gα_i1_. Intriguingly, other entities co-purify including Nrxn1 and Stxbp1 (Figure 1I). It is important to note a soluble domain of neuroligin, a known interactor of Nrxn1^67^, has been shown previously to modulate mGluR2 activity^68^. These findings suggested that a small fraction of the mGluRs isolated from brain homogenates form ternary complexes in the absence of ligands, and to some degree interact with other proteins implicated in presynaptic function.

To further stabilize these physiological ternary complexes, RAPID pulldowns were then performed in the presence of the group II mGluR selective agonist LY35 and the mGluR2-selective positive allosteric modulator (PAM) JNJ-46281222^69^ (JNJ462).

Notably, we found JNJ462 to be a potent ago-PAM for murine mGluR2, with a high degree of selectivity for mGluR2 over mGluR3 (Figure S3A). Including the drugs in combination during homogenization and in detergent purification buffers led to significant increases in relative abundance for endogenous Gα_oA_ and Gβ1 compared to mock controls (Figure 1J). A medium scale RAPID pulldown in the presence of LY35 and JNJ462 followed by further purification with size-exclusion chromatography confirmed the biochemical stability of these endogenous ternary complexes (Figures S3B-S3C). Taken together, these data demonstrate the existence of endogenous mGluR heterodimers in the brain, while also showing LY35 and JNJ462 further stabilize mGluR-G_oA_ ternary complexes for purification with the RAPID approach without the use of state-selective antibodies or apyrase (which is ubiquitously used to eliminate bound guanine nucleotides in GPCR-G protein complexes).

### Conformational and compositional landscape of endogenous mGluR2 complexes

To elucidate the molecular structures of the endogenous mGluR2 assemblies, a scale-up RAPID purification from whole brains was performed in the presence of LY35 and JNJ462 for cryo-EM sample preparation (Figure S3D, Methods). Preliminary two-dimensional (2D) classification analysis of a large dataset indicated considerable conformational heterogeneity within the sample, with the endogenous receptor dimers adopting either the inactive/relaxed or active/compact conformations (Figure S3E, Table S1). Notably, a number of 2D classes of the active state showed a strong extra density at the intracellular side with features consistent with a single G-protein heterotrimer per receptor dimer (Figure S3E). Three-dimensional (3D) classifications based on adopted receptor conformations were first performed, yielding reconstructions of five distinct major states within our dataset: (1) a relaxed dimer with both VFT domains open (ROO); (2) a relaxed dimer with one VFT open and one VFT partially closed (RC_i_O); (3) a relaxed dimer with one VFT open and one VFT completely closed (RCO); (4) an active/compact dimer with both VFT domains closed (ACC); and (5) an ACC dimer bound to a single endogenous G_oA_ containing heterotrimer (Figures S4-S5; see Methods).

Owing to the consistently higher local resolution within the VFTs, we next performed 3D classifications with this domain masked within each distinct conformational state, to investigate the compositional heterogeneity suggested by our proteomics data (Figure S5, see Methods). Notably, the mGluR2-selective PAM JNJ462 and the endogenous G_oA_ heterotrimer proved to be robust subtype-selective markers in active receptor states, thereby enabling the isolation of compositional heterogeneity to a single subunit position within the receptor dimer (Figure S4E). For the inactive states, we took advantage of the pseudo C2 symmetry in the ROO state to perform symmetry expansion and subsequent VFT-focused classification (Figure S5E). For the RCO states, the individual open and closed VFT domains were masked for focused classifications (Figure S5F).

Unambiguous assignments of receptor subtypes in the reconstructions were enabled by distinguishing glycosylation sites, distinct loop conformers in divergent regions, and, ultimately, side-chain densities at non-conserved positions (Figures S5-S6). We subsequently identified particle subsets corresponding to every possible assembly in which at least one subunit of the dimer is unambiguously assigned to an endogenous mGluR subtype and obtained their 3D reconstructions (Figure 2A, Figures S5-S6, Tables S1-4, Methods). Approximately 97% of the distinct particles used to first obtain consensus reconstructions of the 5 major conformational states remained in these 11 obtained reconstructions (Figure 2A, Figures S5-S8, Table S5). We then quantified the number of distinct subunit particle projection counts in our cryo-EM dataset and calculated a mol fraction estimate per subtype, which was cross-referenced with subtype fraction estimates from our untargeted proteomics data (Figure 2B, Table S5, see Methods). We obtained a good agreement between the two orthogonal approaches, revealing mGluR2 (∼84% subunit fraction from LC-MS; ∼85% from cryo-EM) and mGluR3 (∼12% subunit fraction from LC-MS; ∼15% from cryo-EM) are the major subtypes of mGluR2-containing dimers present in the brain. These subtype fraction estimates match well to recent studies on mGluR2 assemblies immunoprecipitated from mouse brain tissue in *n*-dodecyl-β-maltoside (DDM) detergent^49,70^. Notably, we obtained a reconstruction of a heteropentameric mGluR2-mGluR3-G protein ternary complex (Figure 2C). While group III receptors were consistently identified in our proteomics data (e.g. mGluR4/7/8 with an estimated 3.8% combined fraction via LC-MS/MS; Figure 2B), we could not unambiguously identify these subtypes in our cryo-EM dataset despite exhaustive further classification of the “RX” subsets. It is reasonable to speculate these species may be present in these “RX” classes, but their relatively low abundance, additional compositional heterogeneity, and potential flexibility prevents their efficient separation from junk particles. Thus, the group III receptors are below the limits of detection for this current structural approach.

**Figure 2.**
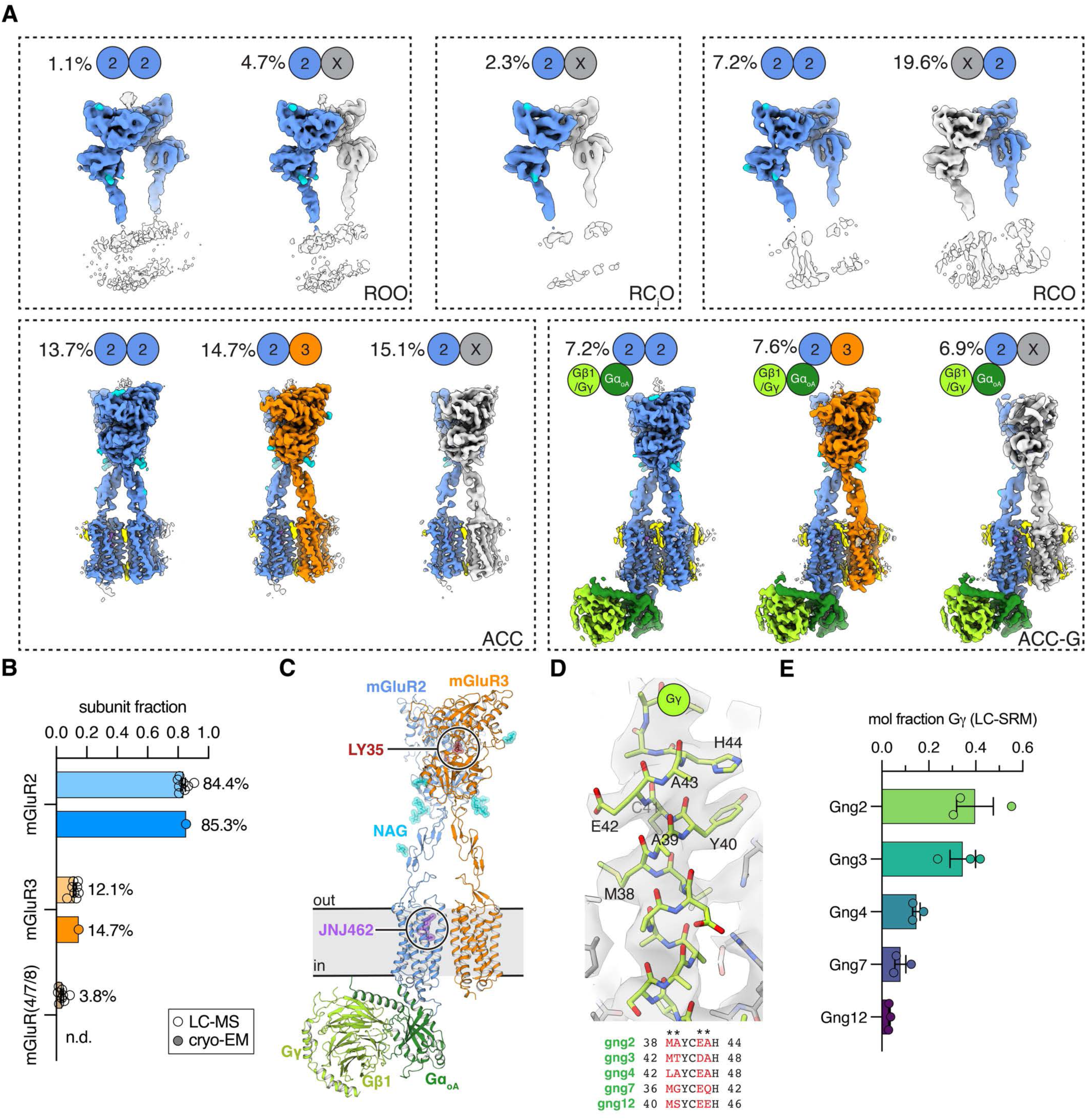
Conformational and compositional landscape of endogenous mGluR2 containing assemblies from mouse brain. **A**, Cryo-EM reconstructions (unsharpened maps) of distinct endogenous mGluR2 containing assemblies. mGluR2 assigned subunits depicted in blue, mGluR3 in orange, Gα_oA_ in dark green, Gβγ in light green, lipids/detergent in yellow, LY35 in dark red, JNJ462 in purple, N-linked carbohydrates (NAG;N-acetylglucosamine) in cyan, and unassigned/ambiguous subunits in grey. Assigned conformational state denoted in bottom right of boxes (“R” – relaxed, “A” – active/compact, “O” – VFT open, “C” VFT closed). **B,** Subunit fractions determined via LC-MS/MS (mass fraction estimates via label-free quantification; *n*=7 independent purifications shown as individual data points with mean and s.e.m.; summary of all proteomics datasets reported for this study) or cryo-EM (subunit mole fractions derived from final particle counts, see Methods). **C**, Structure of the heteropentameric mGluR2/3 endogenous ternary complex (same coloring scheme as panel a). **D,** Local cryo-EM map and model of the endogenous Gγ subunit helix 2 (H2) in the high-resolution G-protein subset. Multiple sequence alignment for the proteomics identified Gγ subunits for this region shown at bottom. **E,** Mol fraction for co-purified Gγ subunit subtypes, determined by LC-SRM (*n*=3 independent experiments shown as individual data points with mean and s.e.m.).

Our proteomics data strongly support Gα_oA_ and Gβ1 as major components of the heterotrimer (Figure 1I). Specifically, Gα_oA_ was the only Gα subtype significantly enriched upon drug incorporation during RAPID pulldown in the controlled experiment (Figure 1J). Additionally, Gβ1 was consistently detected at an abundance of ∼10-fold higher than subtype Gβ2 (Figure 1J, Figure S3D).

To confirm the findings from untargeted LC-MS/MS of Gα_oA_/Gβ1 in the isolated ternary structures, we inspected amino acid positions divergent amongst the subtypes in a higher-resolution reconstruction of the endogenous heterotrimer obtained from focused classification (G-protein-focused subset, Figure S4C). We found the side-chain densities at divergent positions to be most consistent with Gα_oA_/Gβ1 (Figure S9A). The assignment of Gα_oA_ was further substantiated by LC-SRM (liquid chromatography selected reaction monitoring), using proteolytic peptide standards identified from untargeted LC-MS/MS datasets (Figure S9B-C, Table S6). For Gβ, the high conservation amongst the subtypes currently prohibits the design of robust subtype-selective proteolytic peptide standards for LC-SRM for confident subtype quantification (Figure S9B, Table S6).

Owing to the lower molecular weight of Gγ, the label-free abundance values and number of peptide spectrum matches per experiment were consistently low, with different subtypes identified across different datasets (Figure 1I, Figure S3D). Further, within the particle subset that led to a cleaner G-protein signal (G-protein-focused subset, Figure S4C), the best resolved structural region (H2 α-helix) of the Gγ subunit does not enable confident map-based subtype assignment (Figure 2D). Using LC-SRM, we find that Gγ2 and Gγ3 are the predominant subtypes present in the isolated ternary complexes, albeit at mol fractions of ∼0.3-0.4 (Figure 2E, Figure S9B, Table S6). Gγ4 is the next most abundant subtype, followed by Gγ7 and Gγ12 (Figure 2E, Figure S9B, Table S6). Based on these findings, we have tentatively modelled this subunit as Gγ2 in our structures, but emphasize we could not separate the subtypes with further 3D classification.

### Plausible activation trajectory of endogenous metabotropic glutamate receptors

Our current understanding of mGluR activation involves the VFT closure of both subunits, which drives CRD rearrangements, ultimately leading to dimer compaction and the canonical TM6-TM6 packing within the 7TM domain in the active state^13^. These structural features of mGluR activation are enabled by the availability of numerous high-quality structures of various states from recombinant systems^13–20^, including multiple ternary complex structures with recombinant Gα_i_^14–17^. However, there remain minor inconsistencies in many aspects of the reported mGluR activation mechanisms. Notably, there is reported disagreement amongst prior work regarding the symmetry of transitions and intermediate states^18^, the diversity of 7TM arrangements in the inactive states^16,20^, and G-protein interactions^14,15^. Here, we investigated endogenous receptors without the use of artificial expression systems, state-stabilizing antibodies^13,14,51^, or engineered G-proteins^15–17^, which were all unavoidable technical limitations in prior work.

To rigorously interrogate potential differences between endogenous mGluR2 assemblies and mGluR2 isolated recombinantly, we purified a murine mGluR2-mCherry receptor recombinantly expressed with lentivirus in HEK293T cells and solved structural ensembles by cryo-EM under conditions similar to those of our endogenous dataset (Methods, Figure S10, Table S1, Table S4). In addition to the expected absence of co-purified G-proteins and non-mGluR2 subtypes, the ROO and RC_i_O states were not detected in the recombinant mGluR2 dataset. Although the absence of ROO and RC_i_O under these conditions could reflect intrinsic detection limits from two distinct datasets, it does suggest an equilibrium shift in recombinantly expressed mGluR2 receptors compared to the heterogenous endogenous receptors (Figure S10, Table S5).

Given these potential differences, we took the opportunity to revisit the mechanism of mGluR activation using our endogenous mGluR2 cryo-EM dataset as five distinct states were reconstructed from the same sample. To adequately compare the transitions between each conformational state, we used refined coordinates for the mGluR2 homodimer structures for ROO, RCO, ACC, and modeled a poly-alanine chain for the ambiguous C_i_ subunit in the RC_i_O state. Consensus reconstructions for each state, which comprise larger particle projection numbers and higher nominal resolutions, are used for visual comparison of the maps (Figure S5F). We acknowledge the following caveats: 1) we propose a plausible order for transitions between the major conformational states, but note the assigned order is arbitrary in the absence of kinetic data (i.e. smFRET^26^); 2) while the 7TMs are well resolved in our active state reconstructions, they are poorly resolved in the inactive states - therefore, the following analysis is limited to VFT and CRD arrangements in the first 3 transitions; 3) the receptors are solubilized in GDN detergent and thus not in their endogenous lipid environment; 4) while the added orthosteric agonist LY35 is a glutamate analog, JNJ462 is more exogenous as it is a 7TM binding ago-PAM. Incorporation of this R2-selective modulator proved useful in selectively marking mGluR2 subunits in the active state reconstructions to enable facile subtype assignments and subsequent classifications during image processing. Despite these current technical limitations, our data provides the most physiologically relevant structural representation of G-protein coupled receptor activation and transducer coupling to date.

We first inspected a single open mGluR2-VFT, for which we obtained a 3.44 Å GS-FSC focused reconstruction in the symmetry expanded particle stack for ROO (Methods, Figure S5E, Figure S7K) and compared it to a single closed mGluR2-VFT from the 2.82 Å GS-FSC focused R2RX ACC VFT consensus reconstruction (Methods, Figure S5A, Figure S7H). Transitions between these two end states appear to be mediated by a set of state-specific interactions between the upper and lower VFT lobes (Figure 3A). Notably, R271 is ∼13 Å away from D146 in the open state, a distance that decreases to ∼4 Å upon domain closure. These interactions are further bolstered by the presence of LY35, for which we observed strong signal in focused reconstruction maps for all subtype-assigned closed and open states in the dataset, including the resolved mGluR3 VFTs in the active states (Methods, Figure 3B, Figure S8). Orthosteric agonist binding to the VFT in both conformations implies that the glutamate analog shifts an existing equilibrium, owing to increased interactions in the closed state (Figure 3B). The action of orthosteric agonist binding to the VFT in promoting domain closure through bridging the gap between the upper and lower lobes bears resemblance to molecular glues^71–73^ (Figure 3B). Consistent with these structural observations, glutamate was previously shown to bind to the VFT of recombinant mGluR1 in both open and closed states^74^, and is modeled in open VFT subunits in numerous recombinant mGluR cryo-EM structures^16^.

**Figure 3.**
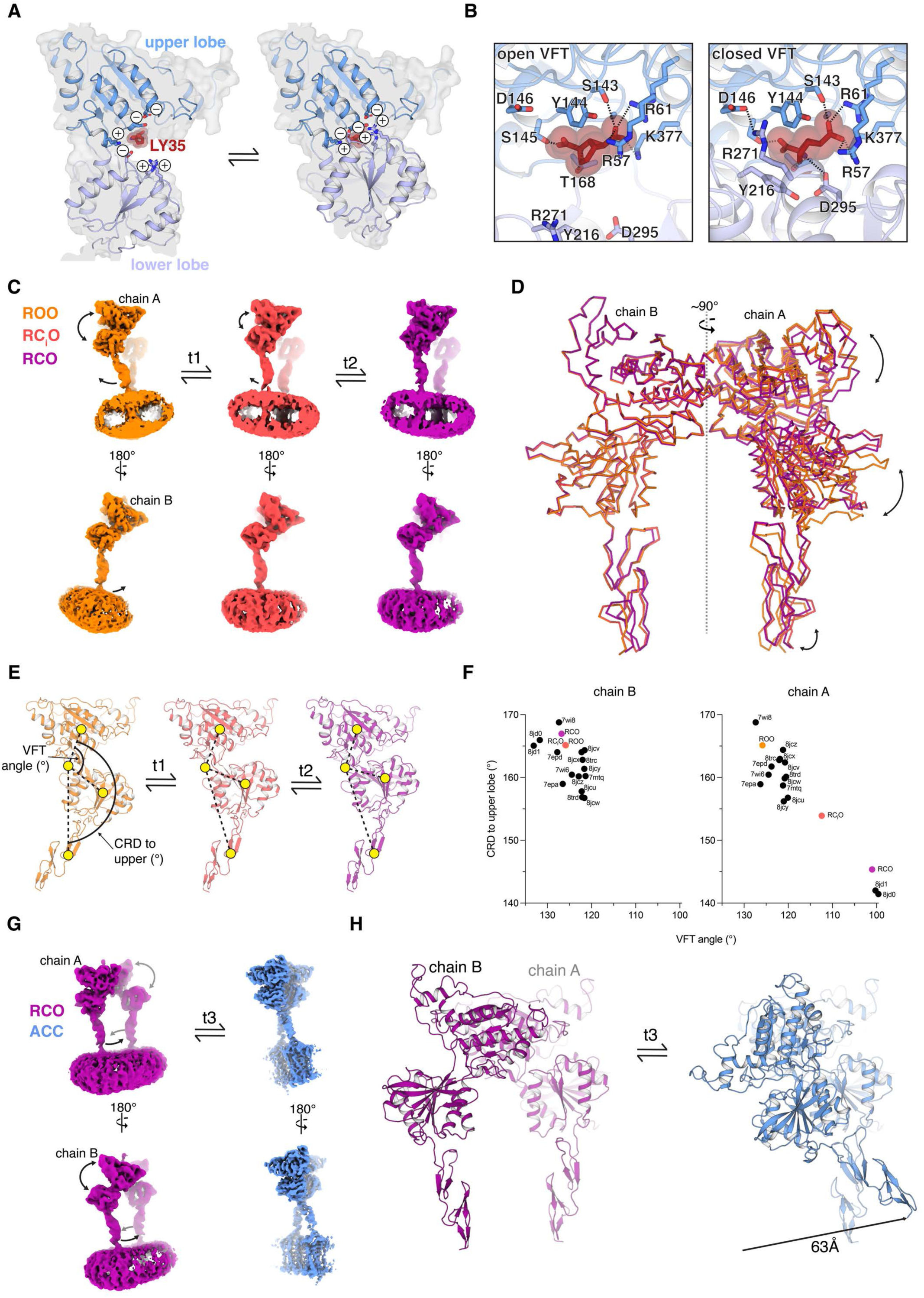
Venus-flytrap domain closure drives CRD rearrangements that lead to dimer compaction. **A**, State-dependent interactions at the interface between the upper and lower VFT lobes, in the open and closed end states. **B,** State-dependent ligand interactions in the orthosteric binding pocket, in the open and closed end states. **C,** Reconstructions of ROO, RC_i_O, and RCO from consensus particle stacks. Volumes manually aligned based on the open VFT protomer. **D,** Structural superposition of the ROO, RC_i_O and RCO states, with alignment focused on the open VFT protomer (chain B; open protomer subunit at left). **E**, First VFT closure, from ROO to RC_i_O and RCO. Angles between domains (VFT upper to lower lobes; VFT upper lobe to CRD) are shown to highlight VFT closure and CRD displacement during the transitions. **F,** Angular measures shown in panel E, depicted per chain for ROO, RCiO, ROO and 15 representative published inactive state mGluR2/3 structures. **G,** Reconstructions of RCO and ACC from consensus particle stacks, highlighting the conformational changes between the two states. **H,** Structural superposition of the RCO and ACC states, with alignment focused on the closed VFT protomer.

Having established the general features of VFT closure in isolation, we then examined the first steps required to prime the inactive ROO endogenous mGluR2 homodimer for compaction into the active state via the first VFT closure event. Structural superpositions of the ROO, RC_i_O, RCO states were performed, in which alignments were targeted to the open VFT subunit to isolate the conformational heterogeneity onto the opposite protomer (chain B; Figure 3C-D). The most obvious conformational change is the sequential closure of the VFT – using center-of-mass points for three distinct subdomains(VFT upper lobe, VFT lower lobe, CRD), with a hinge-point at the upper/lower lobe interface, we measured angles for each state (Figure 3E-F). There is an apparent decrease in the upper/lower VFT lobe angle of ∼13° from ROO to C_i_ (transition 1; t1), and an additional ∼11° from C_i_ to C (transition 2; t2). This closure results in the outward displacement of the CRD, as indicated by the sequential decrease in the VFT upper lobe to CRD angles (Figures 3E-F). When comparing the inter-domain angles of ROO, RC_i_O, and RCO to a representative set of previously published inactive state mGluR2/3 structures, there is clustering amongst the fully open and fully closed protomers in chain A (reference PDBs aligned based on open protomers to chain B of ROO). The RC_i_O state is positioned between these two clusters (Figure 3F).

The transition from RCO to ACC exhibits the largest conformational rearrangement, when manually aligning the consensus reconstructions (Figure 3G). To better visualize this transition, we then aligned the RCO and ACC states, with superposition targeted to the closed VFT of either state (chain A).. Closure of this second VFT (chain B) drives CRD displacement by a direct distance of approximately 60 Å, as measured by the C-terminus of the CRD (Figure 3G-H). This ultimately leads to dimer compaction and the canonical TM6-TM6 packing within the 7TMs in the ACC state^13,15^, which we observe in our active state structures (Figure 2A). In total, the first VFT closure during t1 and t2 displaces the CRD of chain A slightly outward (Figure 3C-F). This appears to prime the dimer for compaction upon the second VFT closure event, which involves the most extensive rearrangement – the chain B CRD rotates in a counterclockwise manner (t3) to the compact, ACC state (Figure 3G-H).

### Structural features of endogenous GαoA coupling to mGluR2 complexes

Signaling complexes of endogenous receptor dimers with strong signal for the endogenous G_oA_ heterotrimer represented ∼30% of the ACC dimers in our cryo-EM dataset (Figure 2A, Table S5). It is important to note every prior reported mGluR-G protein ternary complex structure consists of a recombinantly expressed receptor and recombinant G_i_ (rG_i1_ or rG_i3_)^14–17^. Our proteomics findings and endogenous structures unequivocally show that mGluR2-G_oA_ assemblies are the major endogenous ternary complexes for the glutamate autoreceptors (Figures 1I-1J, Figures S3C-S3D, Figure S9). Therefore, these endogenous complex structures uncover a more physiologically relevant molecular-level understanding of group II mGluR-mediated signal transduction in the brain.

We first evaluated the structural features of JNJ462 recognition in the ACC states, as these structures represent the first experimental structure of JNJ462 in complex with mGluR2^69^. Endogenous G_oA_ heterotrimer engagement occurs on the mGluR2 subunit bound to JNJ462, consistent with reports in recombinant systems that many mGluR2 PAMs signaling is *in cis*^14–17,48^ (Figure 2A, Figure 4A, Figure S4E). Notably, the subtype selectivity exhibited by JNJ462 appears to be driven by sequence divergence in TM5 rather than residues directly contacting the ligand (Figures 4A-4C). Owing to the high degree of selectivity of JNJ462 for mGluR2 over mGluR3 (Figure S3A), we only observed endogenous G_oA_ engagement to mGluR2 receptor subunits in our dataset.

**Figure 4.**
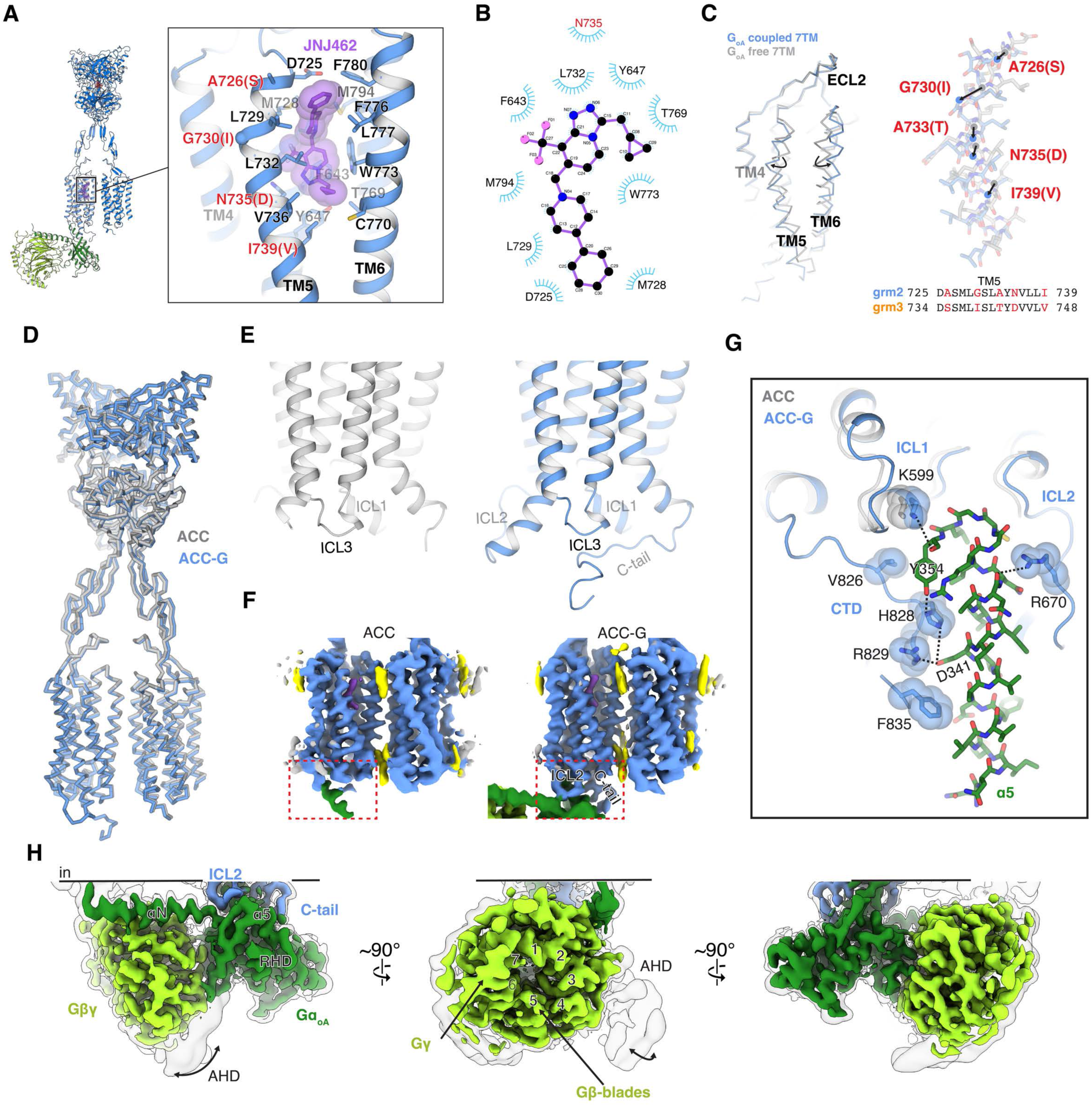
Structures of active-state endogenous mGluRs and physiological ternary complexes. **A**, The 7TM allosteric site in which JNJ462 binds. Side-chains for all residues within 5 Å ligand are shown as sticks. Residues labeled in black are conserved between subtypes R2 and R3, residues labeled in red differ between the subtypes (R3 amino acid substitution shown in parenthesis). **B**, Ligplot representation of the JNJ462 binding site within the 7TM. JNJ462 contacting residues identified that are conserved between R2 and R3 are labeled in black, variable positions are labeled in red. **C**, Structural superposition of the 7TM in either the JNJ462 or the ligand-free/G_oA_-free subunit at left, with focus on TM5 rearrangement at right. Multiple sequence alignment of R2 and R3 at TM5 is shown at bottom right. **D,** Structural superposition of the R2R2-ACC and R2R2-ACC-G structures. **E**, Structural changes within the cytoplasmic side of the JNJ462 bound 7TM in the ACC and ACC-G states. **F**, Unsharpened composite maps for the R2R2-ACC and R2R2-ACC-G structures, highlighting signal increase in ICL2 and the receptor C-tail upon nG_oA_ binding. **G,** Zoom-in view of the brain-isolated complex mGluR-G_oA_ interface, with interacting residues shown as sticks. The JNJ462-bound subunit from the transducer free ACC state is superposed and shown in grey for reference. Significant portions of the ICL2 and CTD were unsolved in the ACC structures and hence left unmodeled. **H,** Focused nG_oA_ subset cryo-EM reconstruction (reconstruction locally filtered in cryoSPARC, threshold=0.45, with transparent rendering overlayed at threshold=0.2 for reference). Gα_oA_ subunit shown in dark green, Gβγ subunits in light green, mGluR2 in blue.

We next investigated the features of the ACC to ACC-G transition (transition 4; t4). While there are no substantial global conformational changes within the receptor dimer during t4 (Figure 4D), we note that ICL2 and the C-terminal tail were poorly resolved in the ACC reconstructions (Figures 4E-4F). This indicates these structural regions potentially become more ordered upon G-protein binding (Figures 4E-4F), which is consistent with what was previously reported in recombinant mGluR2-G_i_ coupling studies^14^. G-protein interaction mainly occurs at the α5 helix of endogenous Gα_oA_ via the endogenous mGluR2 ICL1, ICL2 and C-terminal tail, with some contributions from Ras-like helical domain (RHD) (Figure 4G). In addition to extensive packing interactions, K599 of ICL1 forms ionic interactions with the C-terminal carboxylate of the endogenous Gα_oA_ subunit (Y354), while R670 from ICL2 coordinates a backbone carbonyl of the α5 helix (Figure 4G). Additionally, contributions to the receptor-G protein interaction interface occur at the endogenous mGluR2 C-terminal tail, including ionic interactions between H828 and R829 of receptor with acidic residues on the endogenous Gα_oA_ α5 helix and packing interactions mediated by V826 and F835 (Figure 4G). Considering the orientation of the endogenous Gβ1 subunit in relation the membrane and the absence of guanine nucleotide in the nucleotide binding site (Figure 4H), we assign the functional state of the endogenous G_oA_ as nucleotide free. Indeed, there is weak signal in this higher-resolution focused reconstruction of the endogenous G-protein heterotrimer for the all-helical domain (AHD), consistent with a canonical flexible “open-AHD” nucleotide-free pre-activation state (Figure 4H).

Compared to previous structures of recombinant mGluR2 complexes with recombinant G_i_^14–17^, all of which were solved in complex with scFv16, there are several notable distinctions in G-protein activation (Figure S11A). When targeting structural alignments to the transducer bound 7TM, which aligns well between the structures, rigid body rotations between the endogenous G_oA_ and recombinant G_i_, relative to the membrane inner leaflet, are apparent (Figure S11B). Additionally, there are several differential interactions between receptor and recombinant G_i_ or endogenous G_oA_. The interaction between R2 H828 and the phenolic hydroxyl group of endogenous Gα_oA_ Y354 is not present in recombinant Gα_i_ – a phenylalanine is the final C-terminal residue in the recombinant-G protein. The increased interaction between receptor and transducer within the endogenous complex appears to result in a more ordered receptor C-terminal tail, which contrasts with reported recombinant mGluR2-G_i_ structures (Figure S11C). Whether these differences arise from intrinsic features of G_oA_ vs. G_i_ coupling, or due to recombinant vs. endogenous complex formation, requires further studies.

### Structural evidence for a physiological role of chloride ions in mGluR activation in the brain

At the dimer level, mGluR2/3 heterodimers constitute ∼23% of the particles in the obtained endogenous receptor ensemble, and appear to be exclusively present in the activate states (Figure 2A) – despite exhaustive classification efforts, we did not detect heterodimers in the inactive state dimers (Figure S5). This structural observation suggests mGluR2/3 heterodimers are enriched in the active states. This led us to pose the question: why is the distribution of subtype mGluR3 within the conformational landscape potentially skewed? One plausible explanation is the role of chloride, which has been reported to modulate mGluR activity *in vitro*^75,76^ and present at physiological concentrations in our purification buffers. Indeed, we found increasing expression levels of mGluR3 relative to mGluR2 led to significant decreases in cAMP levels in HEK293T cells under basal conditions in the absence of L-glu (Fig. 5A), consistent with heightened chloride modulation at mGluR3 compared to mGluR2.

**Figure 5.**
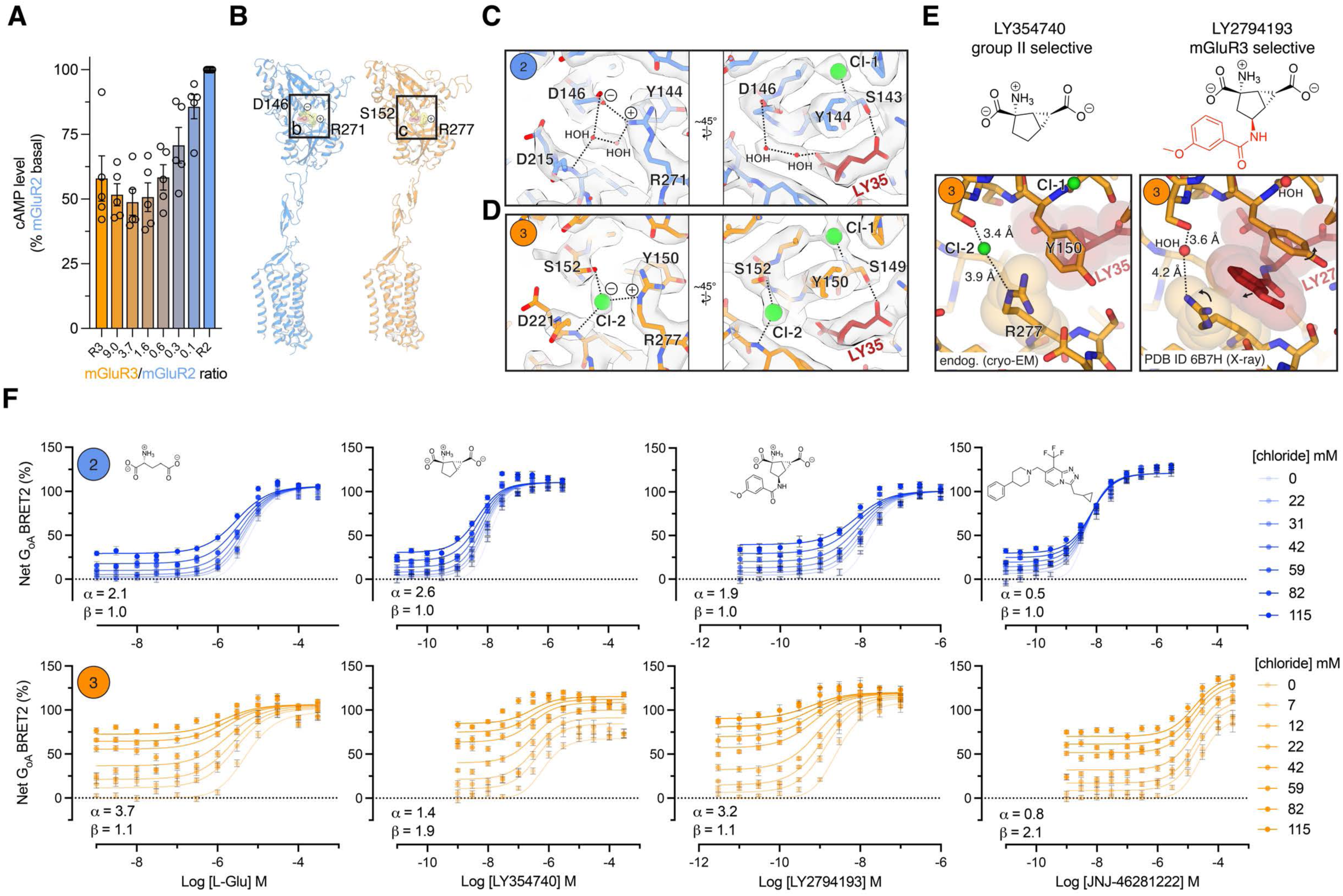
A modulatory role for chloride in physiological mGluR signaling. **A**, Glosensor assay of basal activity for varying ratios of co-transfected mGluR2 and mGluR3 in Locke’s buffer (154 mM NaCl) in the absence of of L-glu (n=5 independent experiments, mean and s.e.m. shown). **B,** Side-view of a single endogenous mGluR protomer for mGluR2 (left) or mGluR3 (right). Location of the chloride modulatory sites boxed, with critical residues and their charges shown. **C,** Chloride modulatory site within the endogenous mGluR2 Venus-flytrap domain (VFT). Map and model from the highest resolution VFT-focused reconstruction for mGluR2 (2.82 Å GS-FSC resolution R2RX VFT focused consensus, sharpened map at threshold=0.5). **D,** Chloride modulatory sites within the endogenous mGluR3 VFT. Map and model from the highest resolution VFT-focused reconstruction for mGluR3 (2.97 Å GS-FSC resolution R2R3 heterodimer VFT focused consensus, sharpened map at threshold=0.5). **E,** Local structural rearrangements at chloride site 2 within mGluR3, induced by mGluR3 selective agonist LY2794193 (LY27), relative to the parent scaffold LY35. **F,** BRET2 (G_oA_) concentration-response assays for L-glutamate, LY35, LY27 and JNJ462, in the presence of various chloride concentrations (murine mGluR2 and mGluR3). Data was fit with the Black-Leff-Ehlert allosteric operational model to describe the allosteric parameters for chloride (response normalized to E_max_ for L-glutamate control at the lowest chloride concentration; n=3-4 independent experiments for each dataset, with mean and s.e.m. shown, see Methods).

We then closely examined previously postulated chloride binding sites^75,76^ within the highest resolution obtained VFT focused reconstructions for mGluR2 and mGluR3 (Figures 5A-5C; 2.82 Å GS-FSC resolution R2RX VFT focused consensus and the 2.97 Å GS-FSC resolution R2R3 heterodimer VFT focused consensus, respectively). These sites are adjacent to the orthosteric agonist binding site, for which we discern unambiguous LY35 density in every obtained mGluR2 and mGluR3 reconstruction^64^ (Figures 5B-5D). In both subtypes, we observed a spherical density consistent with ion in proximity to the mGluR2 S91 sidechain (T98 in R3), as well as the mGluR2 S143 backbone amide (S149 in R3). We refer to this position as chloride site 1 for consistencies sake with previous work^75^ (Figures 5C-5D). This Cl^-^ binding site resides exclusively within the upper lobe of the VFT and does not directly interact with the orthosteric agonist LY35. Notably, mGluR2 S143/mGluR3 S149 sidechains form direct interactions with LY35, bridging the gap between the site 1 chloride and the orthosteric agonist (Figures 5C-5D).

In close proximity to site 1, mGluR2 contains charged side-chain interactions and a water network connecting LY35 to the upper and lower VFT lobes through D146 (upper lobe) and R271 (lower lobe) at this site, which is well resolved in the higher resolution maps (Figures 5B-5D). The cryo-EM map signal at the analogous position in mGluR3 is distinct compared to mGluR2. Specifically, mGluR3 features a serine (S152) at the upper lobe position corresponding to mGluR2 D146, a substitution previously reported to be an important molecular determinant of the higher chloride sensitivity observed in mGluR3^75,76^. Approximately 3 Å from the mGluR3 S152 is a spherical density consistent with ion in the reconstruction, rather than the peanut shaped density consistent with two water molecules in mGluR2 (Figures 5C-5D). This difference in solvation at site 2 between the subtypes is also reflected by the rearrangement of the R277 sidechain (R271 in mGluR2) (Figures 5C-5D). Based on these clear map features, and strong precedent from the prior mutagenesis studies^75,76^, we have tentatively modelled chloride at this site in mGluR3 (“site 2”; Figure 5D). It is important to note these assignments are also strongly supported by prior crystallographic studies that show clear halide spherical OMIT F_o_-F_c_ signals at site 1 (in mGluR2 and mGluR3) and site 2 (mGluR3 only; Figure S12)^77–79^.

Our high-resolution reconstructions of the endogenous mGluR2/3 VFTs support the previously proposed model for chloride mediated mGluR modulation^75,76^, where subtype mGluR2 contains one main chloride site proximal to the orthosteric site while subtype mGluR3 contains two. To further confirm this model and verify our structural findings on the endogenous receptors isolated from mouse brain in the most relevant manner, we tested the allosteric effects of chloride on activation of the murine orthologs using TRUPATH^80^ BRET2 PAM assays. We focused on G_oA_, rather than G_i_ BRET2 or GloSensor (cAMP inhibition relying on endogenous G_i_ activity in HEK293T cells), based on our finding that G_oA_ is the primary transducer for mGluR2 in the brain (Figures 1I-1J, Figure S9).

In PAM assays with L-glutamate, we found general features of allostery in mGluR2 and mGluR3 are consistent with previous reports^75,76^ – chloride coupling with the orthosteric agonist in mGluR2 is affinity-driven, as evident by an affinity cooperativity α=2.1 (data fit with Black-Leff-Ehlert allosteric operational model^81^, see Methods; Figure 5F). Chloride effects on L-glutamate concentration-response in mGluR3 features stronger affinity driven allostery and some weak efficacy driven allostery, with α=3.7 and efficacy cooperativity β=1.1. In agonist-free conditions, the signal is ∼80% of the L-Glu-evoked E_max_ at 115mM chloride. As a control, the 7TM domain binding ago-PAM JNJ462 was also assessed. For both mGluR2 and mGluR3, chloride exhibited subtle negative allostery with JNJ462 (α-values <1), consistent with chloride binding sites distal to the 7TM domain in which JNJ462 binds (Figure 5D). For mGluR3, chloride also had a large efficacy-driven allosteric effect on JNJ462, with a β=2.1 (Figure 5F).

We then utilized LY2794193 (LY27) as a chemical biology tool to pharmacologically probe the second proposed chloride site in mGluR3. This mGluR3-selective orthosteric agonist is a derivative of LY35 (Figure 5E), and appears to selectively target the second chloride site in mGluR3^79^. Specifically, the additional methoxybenzyl substituent group of LY27 displaces R277 and Y150 in the reported recombinant mGluR3 VFT crystal structure, compared to our structure of endogenous mGluR3 in complex with the parent scaffold LY35 (Figure 5E). The resulting rotamer flip of R277 appears to rearrange chloride site 2 (Figure 5E, Figure S12).

As a control, we first quantified the chloride-dependent PAM activity of LY35 (parent scaffold) and LY27 at mGluR2. The fitted α values differed modestly (LY35, 2.6; LY27, 1.9; F-test, p = 0.02; see Source Data). In contrast, the effects at mGluR3 were more pronounced (Figure 5F). In the absence of chloride, LY27 behaves as a full agonist at mGluR3, whereas LY35 exhibits weak partial agonism under the same conditions. In the presence of chloride, LY27 shows a significantly lower β than LY35 (1.1 vs 1.9; F-test, p < 0.0001) but a higher α (3.2 vs 1.4; F-test, p < 0.0001; see Source Data; Figure 5F), indicating that at mGluR3 LY35 exhibits the greater chloride-dependent enhancement of efficacy (β): without chloride it is a partial agonist, leaving headroom for chloride to increase efficacy, whereas LY27’s methoxybenzyl substituent supports full agonism in the chloride-free condition, so chloride exerts only a marginal effect on efficacy, which is apparently compensated by an increase in α.

Taking our structural and pharmacological data in concert with reports from other groups^75,76^, we propose mGluR3’s heightened sensitivity to chloride is largely due to the presence of this second ion binding site. This site appears to form interactions with both the upper and lower VFT lobes, which would ultimately assist in domain closure. This is consistent with the relatively stronger intrinsic agonist activity and efficacy-drive allostery observed for chloride at mGluR3, with the exception of LY27. Our data supports prior structural and pharmacological studies^79^, confirming LY27 perturbs this site (Figures 5E-5F). Direct visualization of these ion-binding sites within the endogenous receptors, and our structural finding that mGluR3 is only detected in the active states, suggests a unique physiological role for chloride in group II mGluR function in the brain.

## Discussion

The low abundance, dynamic nature, and compositional heterogeneity of mGluRs, and GPCRs generally, in the brain have thus limited their structural elucidation to recombinant systems with highly engineered components^13–19^. These prior studies have revealed a wealth of information, but fundamentally limit our understanding of endogenous GPCR physiological signaling mechanisms. In order to gain more biologically relevant insights into endogenous GPCR architecture, composition, and activation trajectory in the brain, we developed a CRISPR/Cas9 edited mouse-line for efficient and rapid affinity purification of class-C GPCR mGluR2 containing complexes directly from whole brain tissue. The purification platform was built off of a recently reported method^65^, which allowed us to proceed from tissue collection to cryo-EM sample vitrification in under 8 hours.

We found that mGluR2 homodimers and mGluR2/3 heterodimers represent the major mGluR2 containing complexes in the mouse brain, as assessed by both proteomics approaches and cryo-EM analysis (Fig. 2). Although our proteomics data also revealed the physiological presence of mGluR2/4, mGluR2/7, and mGluR2/8 heterodimers (Figure 1I, Figures 2A-2B, Figures S3C-S3D), they are minor species in the context of the whole brain and thus below the limits of detection for our current cryo-EM approach. We obtained numerous structures of homo- and heterodimers across 5 main conformational states from the same cryo-EM dataset, uncovering a comprehensive conformational equilibrium, including endogenous ternary complexes.

Our findings suggest that Gα_oA_/Gβ1 represent the major population of Gα/Gβ in isolated ternary complexes from mouse whole brain under these conditions (Fig. 1I-J, Figure S9). Gα_oA_ is well known to have distinct kinetic and signaling properties compared with Gα_i_ ^82–84^, and the compositional make-up Gβγ dimers differentially modulates ion channel activity^5–8^ and SNARE complex formation^9,10^. Intriguingly, we find multiple Gγ subtypes co-purified in the ternary complexes (Figure 2D-E), a finding which implies complex downstream signaling outputs driven by Gγ diversity^85,86^. We also find SNAP25, Syntaxin 1a/b, Syntaxin binding protein 1, and synaptobrevin (VAMP2)--all components of SNARE complexes--co-purify with the brain-isolated ternary complexes in the large-scale preparation used to prepare cryo-EM grids (Figure S3D). The role of Gγ -diversity in these endogenous complexes for both ligand and location-dependent signaling, and their potential interaction with downstream effectors, awaits further exploration.

Using our high-resolution endogenous structures, we revisited the activation mechanism for mGluRs. Our reported scheme (Figure 6) is largely consistent with previous studies conducted using recombinant systems. Notably, our structural data is in agreement with a recent study that proposed a “rolling TMD” model for receptor activation, in which VFT closure repositions the CRD in a manner to enable proper 7TM rotation and domain packing into the active states^20^. There are a few notable distinctions in our structural data on the endogenous receptors compared to previous studies: 1) all states reported in this study were observed from the same biochemical condition without the assistance of varying combinations of agonists, antagonists, allosteric modulators, or state-specific nanobodies to deliberately stabilize various conformations for comparison across many separate datasets, 2) we were able to capture multiple inactive state intermediates that yield insight into how closure of one VFT primes the dimer for compaction in a step-wise fashion, consistent with prior biophysical studies^25,26^, 3) we find potential differences in the conformational equilibria between endogenous and recombinant receptors, 4) we also find differences in endogenous G_oA_ recognition by mGluR2 compared with previously published recombinant mGluR2-G_i_ ternary complexes. Finally, under the conditions used in this study mGluR2/3 heterodimers were only detected in the active states. We propose this may be in part due to a unique chloride activity at subtype mGluR3 (Figure 6). The role of chloride as an allosteric modulator of mGluR activity is consistently reported *in vitro*^75–78^ – our visualization of the ion binding sites in the endogenous receptors provides direct structural evidence that such an activity may play an important physiological role. This subtype-specific functional property appears to shape the conformational landscape of the heterogeneous endogenous receptor assemblies under the conditions used in this study, and in concert with our pharmacology data suggests mGluR3-containing receptor dimers may be hyperactive under physiological chloride concentrations. Regulation of mGluR3 expression levels relative to mGluR2 could potentially be a mechanism to tune the basal threshold for transmitter release from glutamatergic presynaptic. Taken together with a recent study showing extracellular chloride concentrations can exhibit high degrees of local heterogeneity in the brain^87^, mGluR3 activation by this anion adds an additional layer of complexity to glutamatergic signaling. Our finding that mGluR3-containing dimers occupy the active state under physiological chloride raises the possibility that altered mGluR3 levels or local chloride concentrations could shift this balance in disease — a hypothesis our endogenous structural framework is now positioned to test.

**Figure 6.**
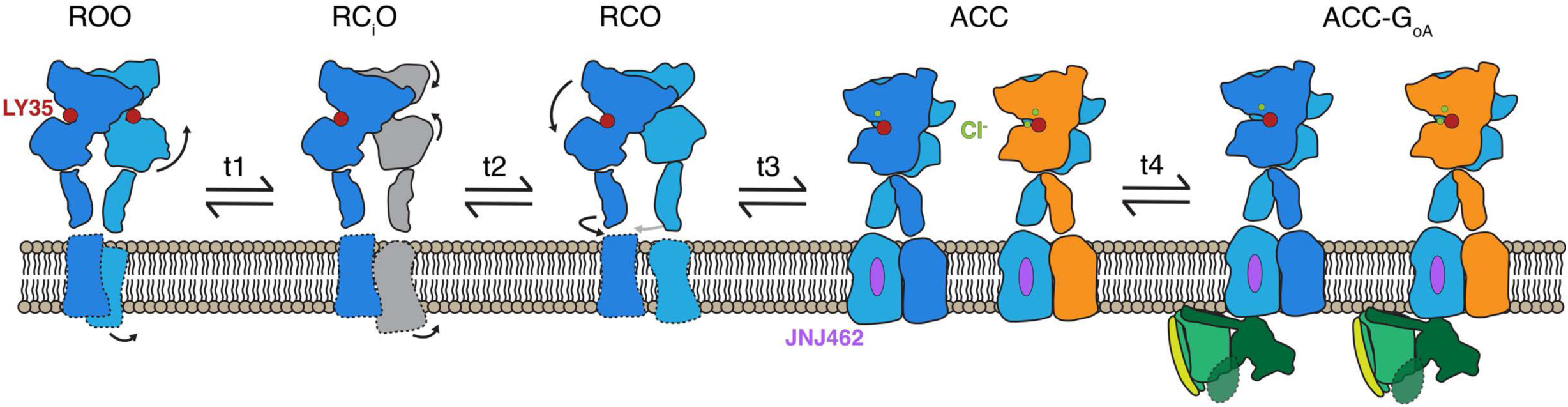
Proposed activation mechanism for endogenous mGluR assemblies in the brain. Cartoon representation of the proposed conformational rearrangements in the endogenous receptors from the inactive ROO state to the active ternary complex. Activation involves sequential rearrangements induced by the first VFT closure (t1 and t2). Closure of the second VFT leads (t3) to dimer compaction and subsequent G_oA_ coupling (t4). In addition to the added chemical modulators, LY35 and JNJ462, the ion chloride plays an important role.

It is important to note this proposed model has potential connections to the pathophysiology of schizophrenia and depression. Indeed, there are disease-associated mutations in *GRM3* identified from previous GWAS studies^31,88^. Furthermore, mGluR3 expression was found to be lower in the dorsolateral prefrontal cortex post-mortem tissue in schizophrenia patients^89^, and mGluR2/3 heterodimers were shown to be present at higher levels in post mortem tissue in patients with severe depression^70^. Our structural findings on the conformational landscape of these endogenous receptor assemblies, in concert with the previous human studies on the pathology of neuropsychiatric disorders, necessitates further exploration as well.

Given these aforementioned findings, it is important to conduct studies on endogenous systems to obtain more physiologically relevant insights into biomolecular structure and function. Here, we provide a generic approach for such efforts, particularly for endogenous neuronal GPCR signaling complexes. This study paves the way for such approaches by focusing on receptor assemblies isolated in detergent from whole brain tissue – thus the complexes represent a bulk average of species present across all mGluR2 expressing cell types present in the mouse brain. Forthcoming work in this area will include single particle cryo-EM analysis of neuronal GPCRs in their endogenous membrane environments, isolated from specific brain regions and ultimately defined cell-types. In combination with frontier methods to visualize the organization of neuronal GPCRs *in situ*, we anticipate more relevant structural mechanisms of signal transduction at chemical synapses will continue to emerge in the near future.

### Limitations of the study

In our view there are two generic approaches to isolating G-protein coupled receptors from brain tissue for structural studies: 1) CRISPR modifications to incorporate affinity tags at the genome level for complex isolation from tissue with established antibody systems, 2) use of antibodies/nanobodies that bind to an epitope on WT receptor with tissue from WT animals. Pursuing the latter subjects one to artifacts related to antibody-induced stabilization of a subset of conformational states and, potentially, blocking interacting partner binding. Genetic modification is not without its limitations – changes in receptor expression, localization, and function are possible with the introduction of any modification. We addressed this by inserting the tag in the flexible C-terminus of mGluR2, and have performed extensive validation of the generated mouse line and found that the introduced tag results in minimal perturbation of receptor function (*in vitro* and *in vivo*), tissue distribution, and resulted in the expected co-purified pre-synaptic entities (Figure 1).

Despite the high sequence conservation of mGluR subtypes between mouse and human, our findings may not be fully relevant to mGluR composition in human brain tissue. A recent report highlights differential populations of mGluR2 homodimers and mGluR2/3 heterodimers in mouse versus human brain regions^70^. Future structural and proteomic studies are required to further interrogate these potential differences.

Our cryo-EM data has uncovered a comprehensive conformational and compositional landscape of brain-derived mGluR2 complexes from this mouse line. Albeit wide-ranging, the detection limits for cryo-EM are inherently limited by various factors (dataset size, microscope specifications, target conformational flexibility, image processing approach, etc). Presumably, with larger datasets, one could detect and obtain 3D reconstructions of additional lowly populated states (i.e. mGluR2/4, 2/7, 2/8 heterodimers; additional intermediate states). These avenues with be important areas of focus in our future work.

## Resource Availability

Coordinates have been deposited in the PDB for the following structures: R2R2-ROO (9PWV), R2RX RCiO (9PWW), R2R2 RCO (9PWX), R2R2 ACC (9PWT), R2R3 ACC (9PWU), R2R2 ACC-G (9PWS), R2R3 ACC-G (9PWR), R2RX ACC VFT consensus (9PWZ), R2R2 ACC VFT consensus (9PX0), R2R3 ACC VFT consensus (9PX1), GoA higher resolution subset (9PWY), R2 ROO VFT C2-expanded (9PX2), recombinant mGluR2 control - ACC state (9PX3), recombinant mGluR2 control - RCO state (9PX4). All cryo-EM reconstructions have been deposited in the Electron Microscopy Data Bank with IDs: R2R2-ROO (EMD-71943), R2RX RC_i_O (EMD-71944), R2R2 RCO (EMD-71945), R2R2 ACC (EMD-71941 – composite; EMD-71931 – ECD focused; EMD-71932 7TM focused; EMD-72048 – global reconstruction), R2R3 ACC (EMD-71942 – composite; EMD-71933 – ECD focused; EMD-71934 – 7TM focused; EMD-72049 – global reconstruction), R2R2 ACC-G (EMD-71940 – composite; EMD-71925 – ECD focused; EMD-71926 – 7TM focused; EMD-71927 – GoA focused; EMD-72046 – global reconstruction), R2R3 ACC-G (EMD-71939 – composite; EMD-71928 – ECD focused; EMD-71929 – 7TM focused; EMD-71930 – GoA focused; EMD-72047 – global reconstruction), R2RX ACC VFT consensus (EMD-71947), R2R2 ACC VFT consensus (EMD-71948), R2R3 ACC VFT consensus (EMD-71949), GoA higher resolution subset (EMD-71946), R2 ROO VFT C2-expanded (EMD-71950), RXRX ROO starting consensus (EMD-72039), RXRX RCiO starting consensus (EMD-72040), R2RX RCO starting consensus (EMD-72041), R2RX ACC starting consensus (EMD-72042), R2RX ACC-G starting consensus (EMD-72051 – composite; EMD-72043 – ECD focused; EMD-72044 7TM focused; EMD-72045 – GoA focused; EMD-72050 – global reconstruction), recombinant mGluR2 control - ACC state (EMD-71951 – composite; EMD-71935 – ECD focused; EMD-71936 – 7TM focused; EMD-72037 – global reconstruction), recombinant mGluR2 control - RCO state (EMD-71952). Raw cryo-EM movies for the brain-isolated mGluR2 dataset will be released on MyEMSL upon publication (PNCC project ID 160598). Motion corrected micrographs for the recombinant mGluR2 control dataset will be uploaded to EMPAIR upon publication. All raw proteomics data will be deposited to the PRIDE repository upon publication. All plasmids constructed for this study will be uploaded to Addgene. The *grm2*^mCherry-FlpO^ mouse line will be deposited to MMRC upon publication. All other source data are provided with this paper. Any additional information is available upon reasonable request.

## Acknowledgments

This research was supported by the National Institutes of Health grants R37DA045657 R01MH112205 (B.L.R), and the Brain and Behavior Research Foundation Young Investigator grant (N.J.W). Cryo-EM samples were screened and collected at the UNC-Chapel Hill CryoEM Core Facility and at the Pacific Northwest Center for Cryo-EM (PNCC) at Oregon Health & Science University (OHSU). A portion of the cryo-EM image processing was performed on the PNCC high performance computing cluster at EMSL Boreal. We acknowledge Clara Lenger and Joshua Strauss at UNC-Chapel Hill for their technical assistance and microscope operation for this project. We thank Nancy Meyers and Trevor Moser at PNCC for assistance with data collection and computing. PNCC at OHSU is supported National Institutes of Health grant R24GM154185. UNC-Chapel Hill NeuroTools Vector Core is supported by BRAIN Initiative Grant U24NS124025.

Microscopy was performed at the UNC Hooker Imaging Core Facility, supported in part by P30 CA016086 Cancer Center Core Support Grant to the UNC Lineberger Comprehensive Cancer Center and Leica STELLARIS 8 FALCON STED microscope supported by NIH grant 1S10OD030300 instrumentation grant to Dr. Stephanie Gupton. CRISPR knock-in mice were generated at the UNC Animal Model Core Facility, which is supported in part by P30CA016086 Cancer Center Core Support Grant to the UNC Lineberger Comprehensive Cancer Center. This research is based in part upon work conducted using the UNC Metabolomics and Proteomics Core Facility, which is also supported in part by NCI Center Core Support Grant (2P30CA016086-45) to the UNC Lineberger Comprehensive Cancer Center. We also thank Zhi Cheng for assistance with computation, and Jon Fay for helpful discussions.

## Author Contributions

N.J.W. and B.L.R. conceptualized the study. N.J.W. designed the purification approach, conducted receptor isolation from tissue for proteomics analysis and cryo-EM, cryo-EM sample freezing, single-particle cryo-EM image processing, model building and refinement, and structural analysis. Y.T.C. performed all IHC experiments, contributed to tissue harvesting, and performed mouse line validation. K. S. performed all of the pharmacology assays and their subsequent data analysis. D.D.K. designed the mouse line and produced lentivirus. P. L. performed mouse behavioral assays under the supervision of J.W. B.A.F. purified nanobody and assisted with brain tissue harvesting. K. H. performed mouse breeding. S. B. performed LC-SRM data analysis. L.G. provided guidance for LC-SRM experimental design and data analysis. S.P.L. performed untargeted LC-MS/MS data analysis and proteomics database depositions. K.L.H. contributed to RNAscope experiments under the supervision of G.S. N.J.W. and B.L.R. wrote the paper with input from all authors.

## Declaration of interests

B.L.R. is currently SAB/Founder of Epidoyne, SkapeBio and ImprintBio.

## Supplemental figure titles and legends

**Figure S1 | Validation of mGluR2-mCherry and *Grm2*^mCherry-FlpO^ transgenic line**

**A,** Dose-response curves from TRUPATH for the WT unmodified and mCherry tagged mGluR2 (n=3 independent experiments; data shown as mean and s.e.m.) **B,** *Grm2* ^+/mCherry-FlpO^ x RCE (R26R CAG-boosted EGFP):FRT mice were used to validate FlpO functionality. Immunofluorescence was performed on brain sections an anti-RFP antibody (1:1000) to enhance the mGluR2–mCherry signal. GFP expression, resulting from FlpO–FRT–mediated recombination was detected and imaged using an Olympus slide scanner with a 10× objective. The experiments in this Figure were conducted with 3 mice with similar results.

**Figure S2 | Nanobody-based receptor isolation optimization and validation**

**A,** Overview of the RAPID construct (top). Typical detection limits and dynamic range with fluorescence HPLC shown at bottom left. Timescale of SENP_EuB_ protease mediated liberation of nanobody from streptavidin magnetic beads at bottom right. **B,** Cartoon schematic of RAPID mediated target isolation from cell lysates. **C,** Fluorescence HPLC analysis pipeline of RAPID isolation of recombinantly expressed (lentivirus) mGluR2-mCherry from HEK293T cells. Bead loading is evident by target peak decrease in the flowthrough (mCherry signal; top left); target monodispersity in bead elution (mCherry signal; bottom left) and SEC profile (A280 signal; top right) indicates biochemically stable receptor. SDS-PAGE analysis confirms target purity and identity (top right). Target identity was also confirmed by fluorescence HPLC of the SEC fractions, utilizing both mCherry and mTFP1 signal (bottom right).

**Figure S3 | RAPID isolation of mGluR2 containing endogenous complexes from brain tissue**

**A,** Concentration-response of L-glutamate, LY35, or JNJ at the murine receptors as assessed by G_oA_ BRET2. Assays were performed in chloride-containing buffer for mGluR2 and in chloride-free buffer for mGluR3 (see Methods; data shown as mean and s.e.m. from *n*=3 experiments). **B,** Tissue work-up pipeline chart for isolation of mGluR2 complexes from mouse whole brains. **C,** Medium-scale mGluR2 complex RAPID isolation and subsequent size-exclusion chromatography (SEC) from whole brain tissue (10 brains). The dashed lines in the top left panel denote range of SEC fractions taken to FSEC analysis (F22-30), while the dark shaded box denotes SEC pooled fractions for SDS-PAGE and subsequent proteomics analysis (F23-26). Top 10 hits as ranked by peptide spectrum matches (PSMs) are shown in the top right panel, with possible homo-and heterodimeric assemblies shown in the bottom right panel. **D**, Size-exclusion chromatography profile of GDN-purified endogenous mGluR2 assemblies (top left), SDS-PAGE analysis of pooled peak fractions used for cryo-EM vitrification (top right), and proteomics analysis of selected band slices from the gel (bottom). **E**, Representative micrograph from the endogenous mGluR2 cryo-EM dataset (top) and representative 2D classes of the receptor particle projections (bottom).

**Figure S4 | Initial image processing and 3D classification based on conformational state**

**A,** Motion correction, CTF correction, particle picking, 2D classification and initial 3D classifications. **B**, Representative pick locations, 2D classes, and 3D classes from one of the replicate picking/classification runs. **C,** Processing pipeline for the active state conformational states, based on G-protein signal strength. **D,** Processing pipeline for the inactive states, based on overall receptor conformation. **E,** Composite cryo-EM map (unsharpened, threshold=0.25) for the higher resolution consensus reconstruction of the ternary complex. The R2 assigned subunit is colored in blue, ambiguous subunit colored in grey, agonist colored red, ago-PAM colored purple, glycosylation (N-acetylglucosamine; NAG) in cyan, G-proteins colored green and detergents/brain lipids colored in yellow. A lower threshold (0.15) map rendering is overlayed in transparent grey to show the GDN micelle in relation to the protein elements of the complex.

**Figure S5 | VFT focused classifications based on subtypes**

**A,** Focused refinements and classifications of the VFT dimer within the active state consensus stack, to separate receptor subtypes. **B**, Schematic for finding particle intersects for distinct assemblies in distinct receptor states. **C**, Final reconstructions for distinct assemblies in the ACC-G state (unsharpened composite maps from ECD, 7TM and G-protein local refinements). GS-FSC resolutions from the local maps are shown next to the domain. **D**, Final reconstructions for distinct assemblies in the ACC state (unsharpened composite maps from ECD and 7TM local refinements). GS-FSC resolutions from the local maps are shown next to the domain. **E,** VFT focused classification (symmetry expansion) within the ROO state to identify R2 subtype subunit particles. **F**, VFT focused classification of the open or closed protomers within RCO to identify R2 subtype subunit particles. A partially closed state (C_i_) was identified from further classification of the closed protomer. **G**, Reversion of symmetry expansion to identify particle subsets and obtain reconstructions for R2R2 ROO homodimer and R2RX ROO dimer. **H**, Particle intersection to identify particle subsets and obtain reconstructions for R2RX RC_i_O dimer, R2R2 RCO homodimer, and R2RX RCO dimer.

**Figure S6 | mGluR2 and mGluR3 subtype distinguishing features in the ACC states**

**A,** Overlay of the higher resolution R2R2 ACC VFT consensus and R2R3 ACC VFT consensus cryo-EM reconstructions, highlighting differences in divergent loop regions and subtype-specific glycosties (unsharpened maps shown). **B,** Subtype distinguishing positions within the VFT at chain B (non-G_oA_ coupled subunit), highlighting differences in side-chain densities and subtype-specific glycosties (auto sharpened maps shown or the R2R2 ACC VFT and R2R3 ACC VFT consensus reconstructions). Multiple sequence alignment of subtypes R2 and R3 shown above each highlighted position. **C,** Subtype distinguishing positions within the 7TM domain of the final ACC-G reconstructions, highlighting differences in side-chain densities (chain B of the final R2R2 ACC-G and R2R3 ACC-G reconstructions; sharpened composite maps used). Signal from this domain was not utilized in 3D classifications to sort homodimers and heterodimers.

**Figure S7 | GS-FSC curves, particle assigned angular distributions, and local resolution for the endogenous receptor active state reconstructions**

Conical GS-FSC, masked GS-FSC, particle assigned orientation distributions and local resolution estimates for the ECD focused (left), 7TM focused (center), and G-protein focused (right) reconstructions for the distinct ACC-G states: **A,** R2R2-ACC-G; **B,** R2R3-ACC-G. Conical GS-FSC, masked GS-FSC, particle assigned orientation distributions and local resolution estimates for the ECD focused (left), 7TM focused (right), and G-protein focused (right) reconstructions for the distinct ACC states: **C,** R2R2-ACC; **D,** R2R3-ACC. Conical GS-FSC, masked GS-FSC, particle assigned orientation distributions and local resolution estimates for inactive states: **E**, R2R2-ROO; **F**, R2RX-RC_i_O; **G**, R2R2-RCO. Conical GS-FSC, masked GS-FSC, particle assigned orientation distributions and local resolution estimates for the local reconstructions: **H,** R2RX ACC VFT consensus; **I,** R2R2 ACC VFT consensus; **J,** R2R3 ACC VFT consensus; **K,** R2 ROO VFT C2-expanded; **L,** G_oA_ higher resolution subset. Volume regions outside of the refinement masks that have no local resolution value assigned are shown in grey.

**Figure S8 | Ligand densities for the endogenous receptor reconstructions**

Cryo-EM map signal ligands in the endogenous receptor reconstructions: **A**, R2R2 ACC VFT consensus; **B**, R2R3 ACC VFT consensus; **C**, R2RX ACC VFT consensus; **D**, R2R2 ACC; **E**, R2R3 ACC; **F**, R2R2 ACC-G; **G**, R2R3 ACC-G; **H**, R2 ROO VFT (C2-expanded); **I**, R2R2 ROO; **J**, R2R2 RCO. LY35 colored in dark red, JNJ in purple, R2 protein in blue and R3 protein in orange. The title of the reconstruction and chain ID is shown per panel, and the map threshold used is shown at the bottom right of each panel.

**Figure S9 | Map-based subtype assignment for Gα and Gβ subunits**

**A**, Cryo-EM map and model for the high-resolution G-protein local subset, with multiple sequence alignment of relevant subtypes shown above each structural Region. Gα model depicted in dark green, Gβ model depicted in light green. Divergent positions of interest highlighted in red. **B**, Representative standard curves for LC-SRM for each proteolytic peptide standard, with unknown sample peak integrations plotted from a representative mGluR2 purification (drugs added). **C**, Summary of mol fractions of Gnao1, Gnai1 and Gnaq co-purified with mGluR2 from brain homogenates (drugs added) determined from LC-SRM (n=3 independent purifications).

**Figure S10 | Image processing and validation of the recombinant control mGluR2 dataset**

**A,** Processing pipeline for the recombinant mGluR2 dataset. Conical GS-FSC, masked GS-FSC, particle assigned orientation distributions and local resolution estimates for: **B**, the ACC-ECD focused reconstruction; **C,** the ACC-7TM focused reconstruction; **D,** RCO-ECD focused reconstruction. **E,** Consensus reconstructions from the endogenous dataset compared with the consensus reconstructions from the recombinant mGluR2 control dataset (**F**). **G,** Particle population distributions across the conformational equilibria for either dataset. Only distinct particles were used for the distribution calculation (see Methods).

**Figure S11 | Endogenous mGluR2-GoA protein interactions compared with recombinant mGluR2-Gi1**

**A**, Comparison of the endogenous R2R2-ACC-G structure with representative published recombinant mGluR2-G_i_ complex structures (PDB IDs 7E9G and 7MTS). **B,** Structural superposition of the endogenous R2R2-ACC-G structure with 7E9G and 7MTS. Alignment is targeted on the G-protein coupled 7TM**. C,** Divergent residues between Gnao1 and Gnai1 shown in yellow on the endogenous R2R2-ACC-G complex and on a representative recombinant mGluR2-G_i1_ structure (7MTS). **D,** Comparison of the G-protein interaction interface in the endogenous or recombinant (7MTS) complex structures. The interaction interface is mainly comprised of receptor elements ICL1, ICL2 and C-terminus, in addition to the alpha-5 helix of the Gα subunit.

**Figure S12 | VFT ion modulatory sites within previous published X-ray structures**

OMIT mFo-DFc difference maps at the proposed chloride sites in previously published crystal structures of recombinantly expressed mGluR2 or mGluR3 VFTs (see Methods). Reported resolution and PDB ID shown in top left of each panel.

## Supplemental tables

**Table S1.**
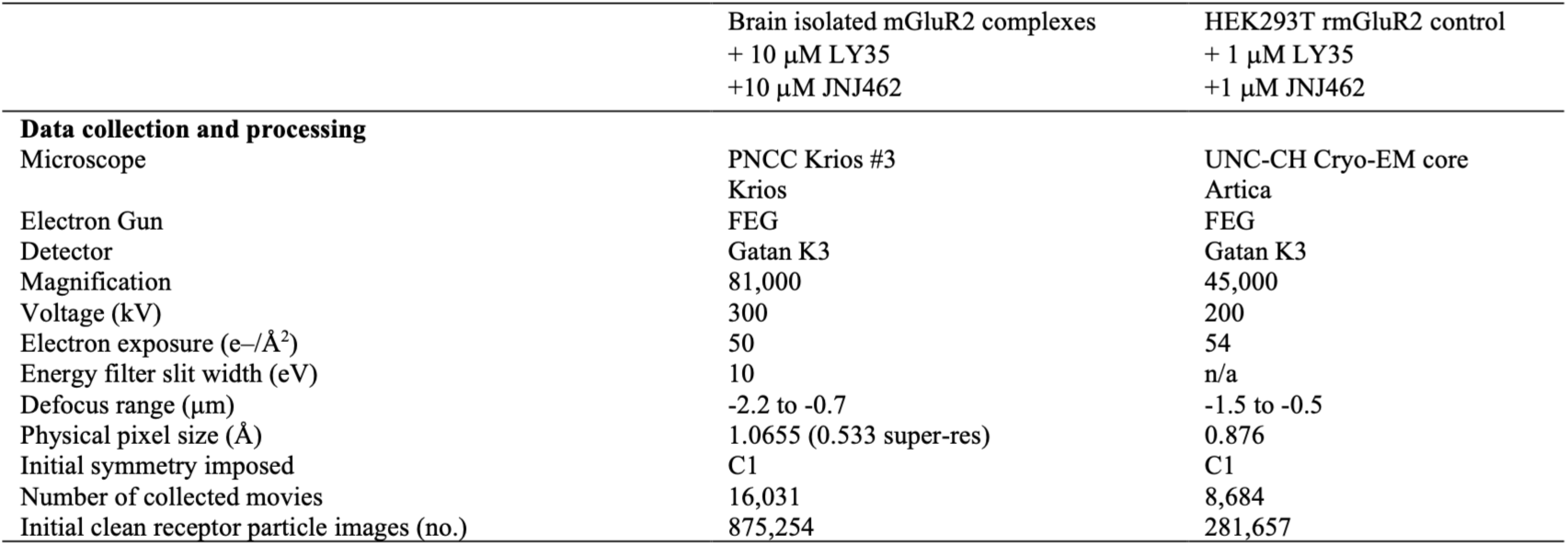
Cryo-EM data collection statistics.

**Table S2.**
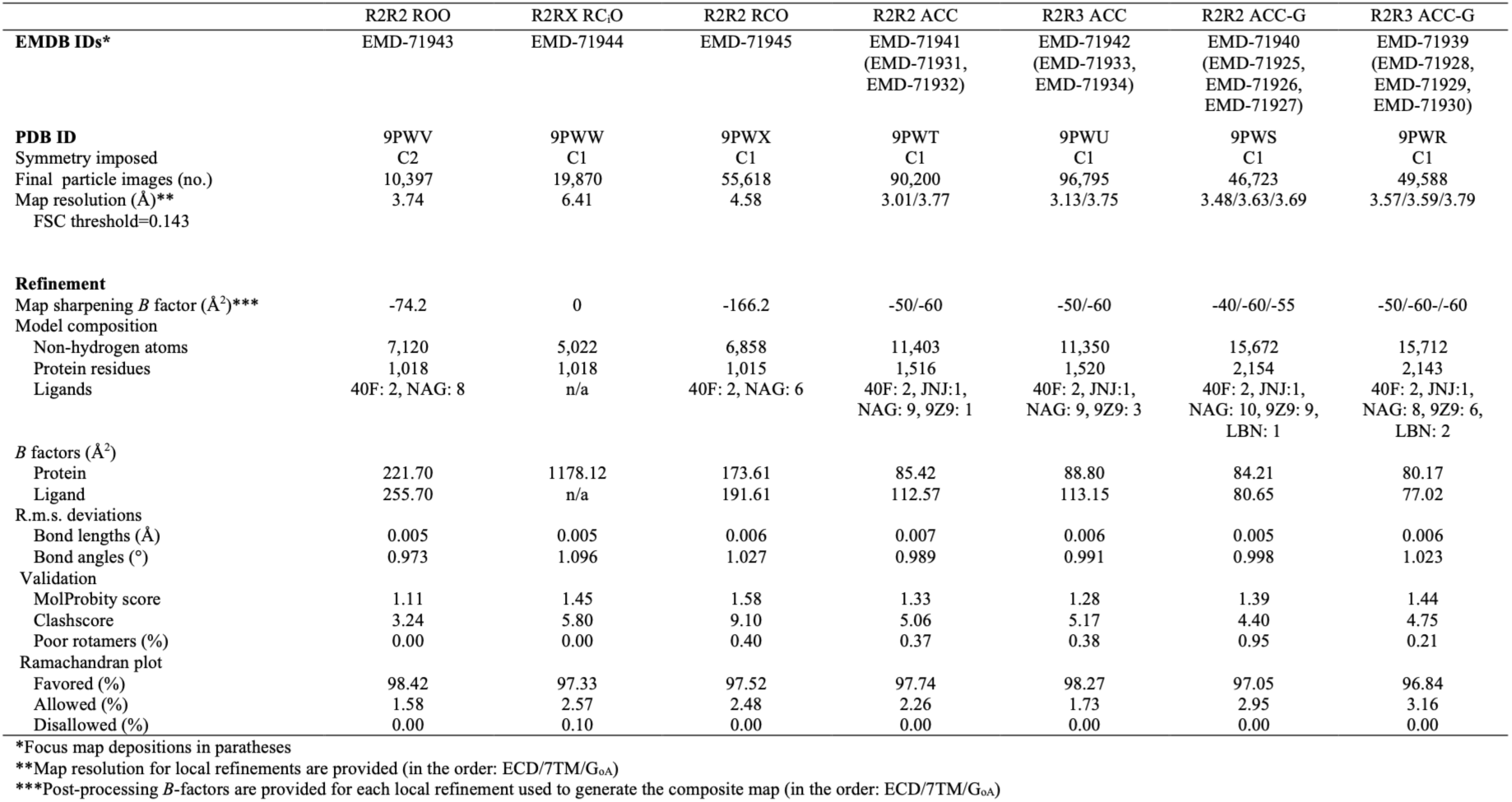
Refinement and validation statistics for brain-isolated mGluR2 assemblies (global and composite reconstructions)

**Table S3.**
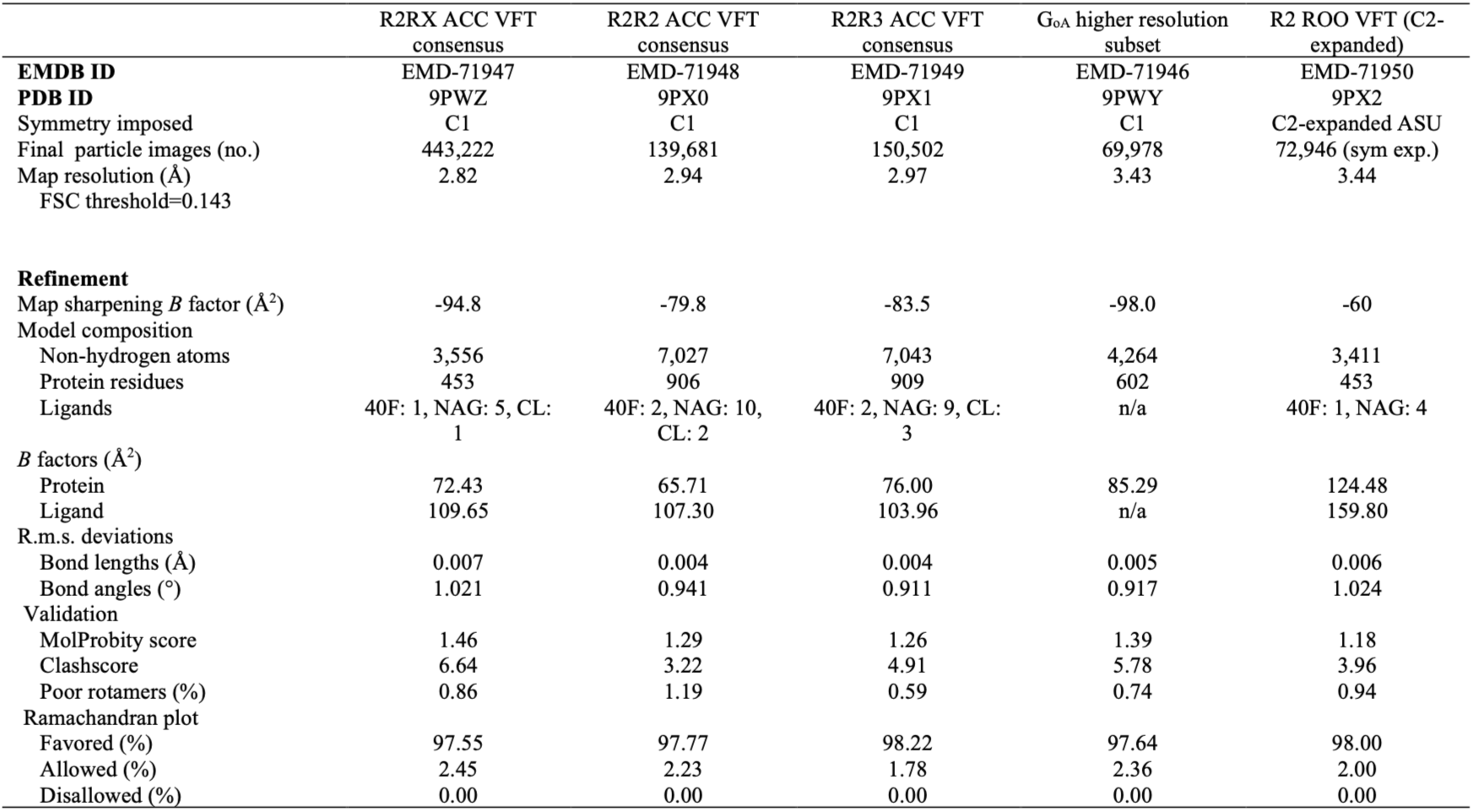
Refinement and validation statistics for brain-isolated mGluR2 assemblies (focused reconstructions)

**Table S4.**
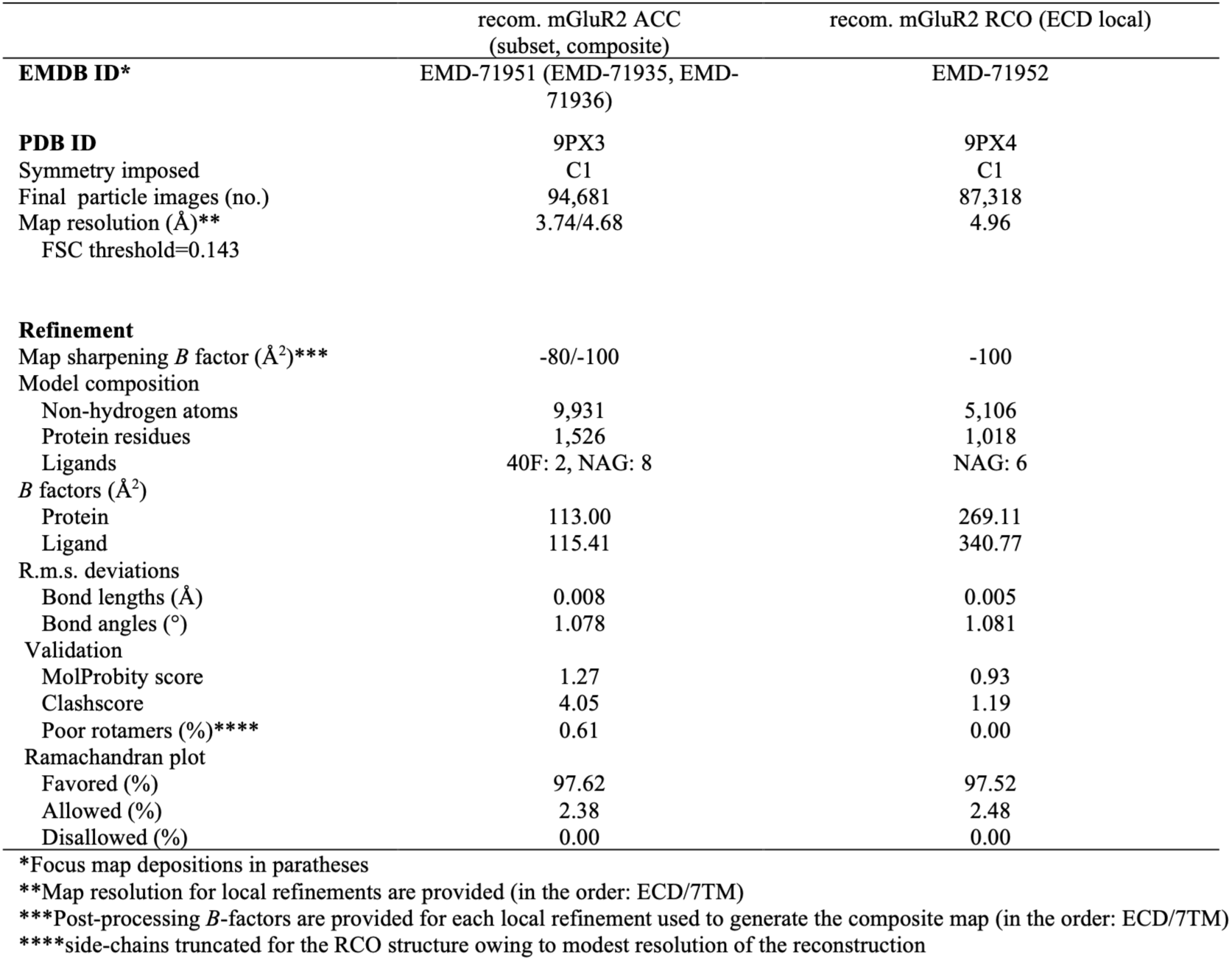
Refinement and validation statistics for recombinant mGluR2.

**Table S5.**
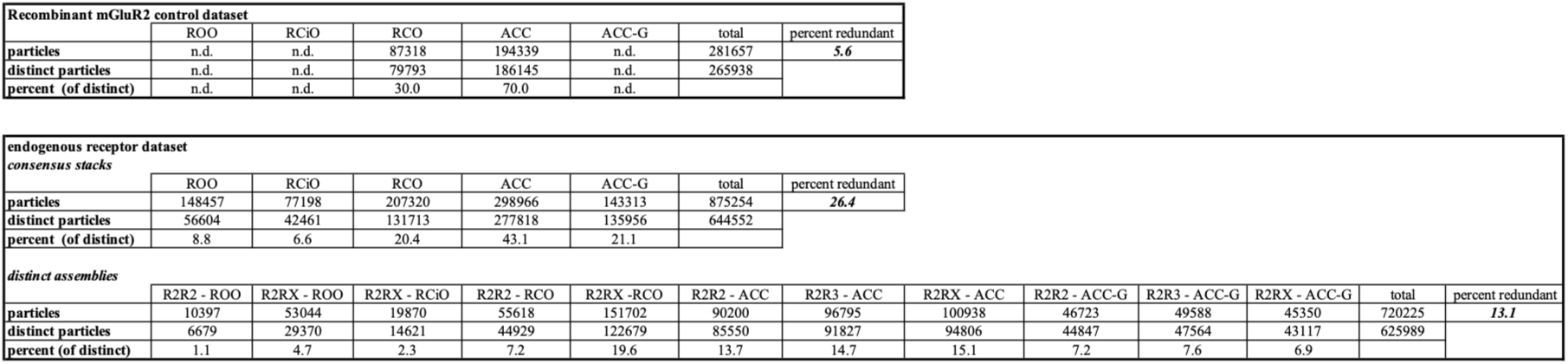
Final distinct particle counts used for population analyses.

**Table S6.**
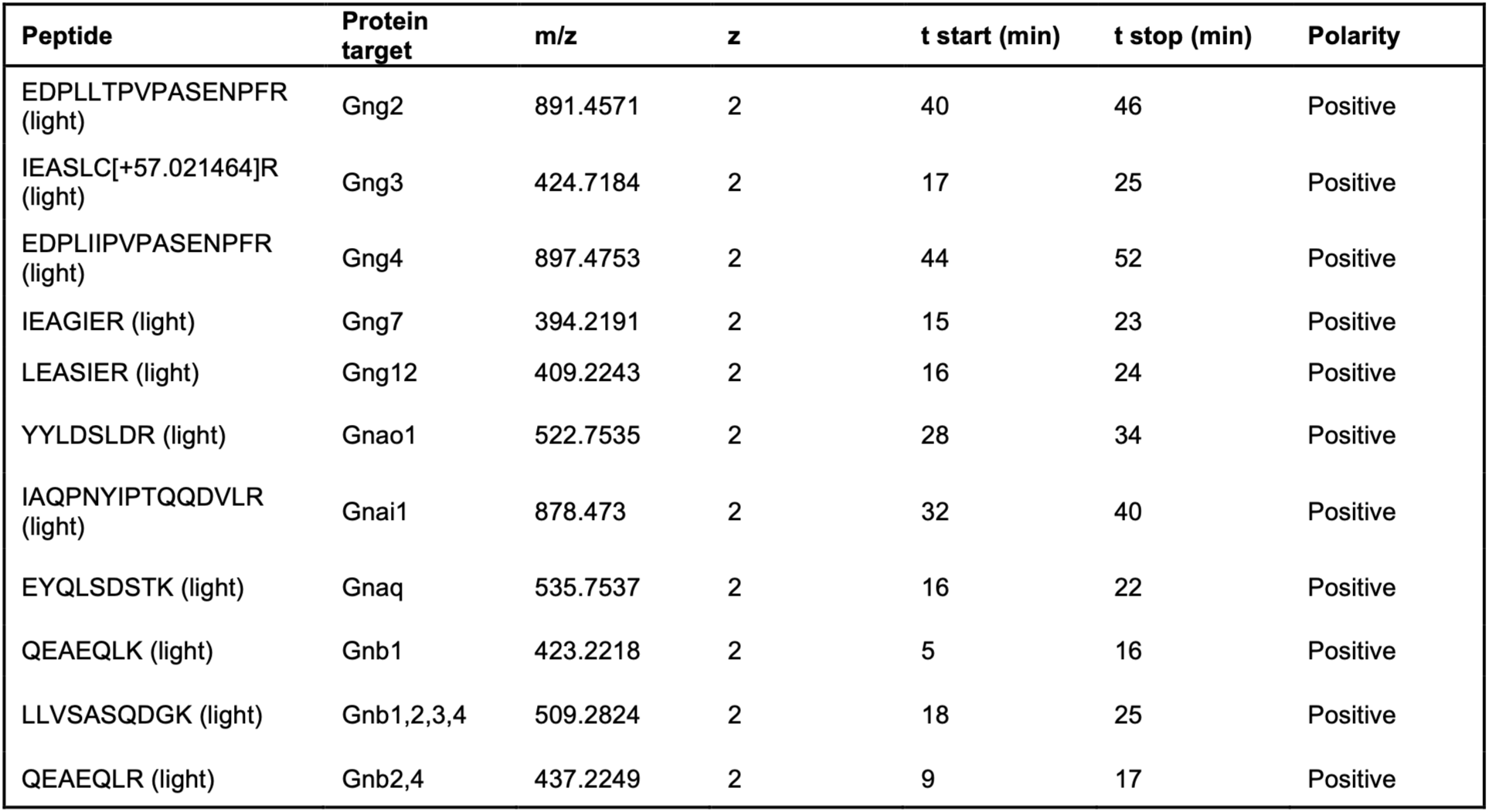
Proteolytic peptide standards for LC-SRM.

**Table S7.**
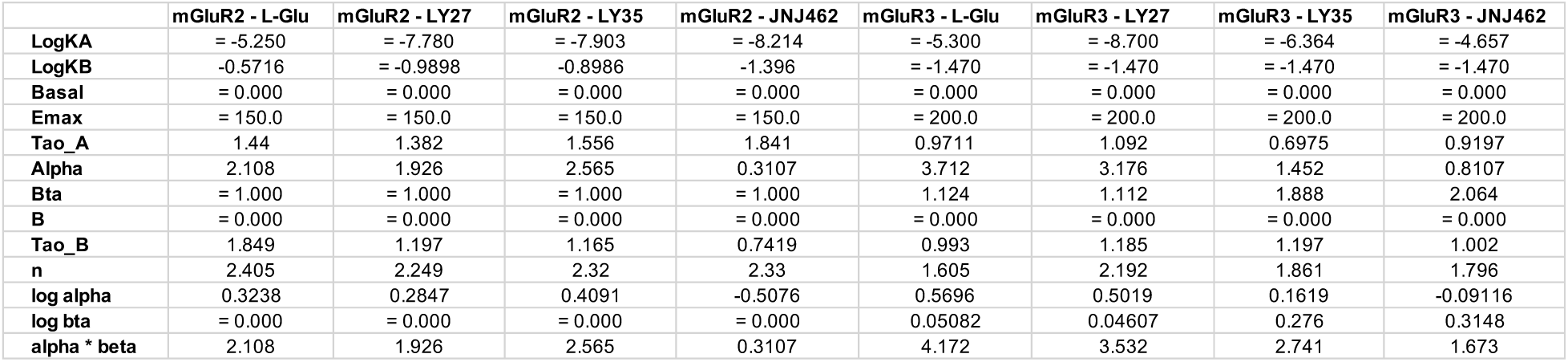
Fitting parameters for chloride PAM assays.

## Methods

### Animals

C57BL/6J and RCE (R26R CAG-boosted EGFP):FRT mice (#000664 and#010812, Jackson Laboratories, Bar Harbor, ME) and in-house mGluR2 knock-in mice [mGluR2-mCherry-FlpO (*Grm2*^mCherry-FlpO^ mice)] were used in these studies. All mice were on a C57BL/6J genetic background. All animal handling and experiments were carried out in accordance with the National Institutes of Health Guide for Care and Use of Laboratory Animals and as approved by the Division of Comparative Medicine (DCM) of the University of North Carolina at Chapel Hill. All animals were housed 1-5 per cage in temperature- and relative humidity-controlled room with a 12:12-hr light/dark cycle with food and water provided *ad libitum*.

### *Grm2*^mCherry-FlpO^ mouse line generation

#### Production of Cas9 protein, guide RNAs, and donor vectors

For the production of recombinant Cas9 protein, a human codon-optimized FLAG-Cas9 cDNA (Addgene 42230) was modified by C-terminal insertion of an additional nuclear localization signal and 6His tag and cloned into the pET-28a(+) vector (Novagen/Sigma-Aldrich, St. Louis, MO, USA). Cas9 protein expression and purification were performed by the UNC Protein Expression and Purification Core Facility. The Cas9 protein was purified using a HisTrap Ni–NTA column followed by SP cation exchange column^90^, and size exclusion column. The final protein was stored in 20 mM HEPES pH 7.5, 150 mM KCl, 1 mM DTT, 50% glycerol. Cas9 guide RNAs in target regions of the mouse Grm2 (mGluR2) locus were identified using Benchling software (www.benchling.com). Selected guide RNAs were cloned into a T7 promoter-guide RNA vector (UNC Animal Models Core) followed by T7 in vitro transcription (HiScribe T7 High Yield RNA Synthesis Kit, New England BioLabs) and RNeasy spin column purification (Qiagen), with elution in microinjection buffer (5 mM Tris-HCl pH 7.5, 0.1 mM EDTA). Functional testing was performed by co-electroporating a mouse embryonic fibroblast cell line with guide RNA and Cas9 protein. The guide RNA target site was amplified from transfected cells and PCR products were analyzed by Sanger sequencing followed by ICE analysis (Synthego) to detect Cas9 cleavage and indel formation. The guide RNAs selected for production of knock-in mice were: mGluR2-5g70T (5’-gACTGTTTGCAATGGCCGTG-3’) and mGluR2-3g66T (5’-gCAGCGTAGACCCTCTTCCA-3’). A double-stranded supercoiled DNA donor plasmid was used to generate the insertion event. The donor plasmid included 1) a 1000 bp 5’ homology arm; 2) linker-mCherry-linker cassette; 3) C-terminal 36 bp mGluR2 coding sequence and stop codon; 4) spacer and EMCV IRES; 5) codon-optimized FlpO recombinase coding sequence; 6) 1004 bp 3’ homology arm with point mutations inserted to disrupt the PAM site of the 3’ guide RNA and a second 3’ guide RNA that was not used. The donor plasmid was prepared by HiSpeed Plasmid Maxi Kit (Qiagen). Eluted DNA was spot dialyzed in microinjection buffer.

#### Transgenic mouse production

C57BL/6J zygotes were microinjected with 400 nM Cas9 protein, 25 ng/ul each guide RNA and 20 ng/ul donor vector. To prepare the microinjection mix, guide RNAs were diluted in microinjection buffer, heated at 95°C for 3 min, and placed on ice prior to addition of Cas9 protein. The mixture was then incubated at 37°C for 5 min and placed on ice, after which the donor vector was added, and the mixture was held on ice prior to pronuclear microinjection. Microinjected embryos were implanted in recipient pseudopregnant B6D2F1 females (#100006, Jackson Labs). Resulting pups were screened by PCR and sequencing for the presence of the correct insertion allele. Two founders with the correct allele were mated to wild-type C57BL/6J animals to transmit the modified allele through the germline. Offspring from a single founder line were selected for additional breeding to maintain and characterize the line.

### In situ hybridization

For in situ hybridization experiments, we used the RNAscope Multiplex Fluorescent Reagent Kit v2 assay (Advanced Cell Diagnostics Bio, Newark, CA, USA), with slight modifications. Fresh brains from C57BL/6J (*Grm2*^+/+^) and *Grm2*^mCherry-FlpO/mCherry-FlpO^ mice (8-12 weeks) were dissected and immediately embedded into O.C.T specimen matrix (Tissue-Tek® O.C.T Compound, Sakura ®Finetek). Brain sections at 18 µm thickness were cut using a cryostat and directly mounted onto slides (Superfrost^TM^ Plus microscope slides, Fisherbrand^TM^) followed by 1 hr post-fixation with 4% paraformaldehyde (PFA) in PBS at 4°C. Brain sections were passed through a serial gradient ethanol dehydration step, followed by H_2_O_2_ treatment to quench endogenous peroxidase activity. The sections were next subjected to protease 3 digestion for 20 min. Mouse *Grm2* (# 317831 -C1, Advanced Cell Diagnostics Bio) and *mCherry* (#1569311-C3 customization from ACD Bio) probes were used for hybridization over 2 hrs at 40°C in a humidity-controlled oven (HybEZII; Advanced Cell Diagnostics Bio) and then the signal was amplified using Opal Dye570 for the *Grm2* probe and Opal Dye690 for the *mCherry* probe. Slides were counterstained with DAPI and mounted. Images were collected under an Olympus VS200 virtual slide microscope (Olympus, Tokyo, Japan) and Leica STELLARIS8 FALCON STED Microscope (Leica, Wetzlar, German).

### Immunohistochemistry

*Grm2*^mCherry-FlpO/mCherry-Flpo^ mice *or Grm2*^+/mCherry-FlpO^ *x RCE:FRT mice (8-12 weeks)* were euthanized and were intracardially perfused with heparin (10 unit/ml, Sigma) in PBS and then 4% PFA in PBS. Brains were harvested, post-fixed in 4% PFA/PBS overnight, and dehydrated in 30% sucrose/PBS until sinking. Brains were sectioned by cryostat at 40 µm. The free-floating brain sections were washed 3 times with 0.1% TX-100/PBS prior to a 1 hr incubation in blocking buffer, 5% normal donkey serum (Sigma) in 0.4%TX-100/PBS and then incubated overnight at 4°C with anti-rabbit RFP (1:1000, #600-401-379; Rockland, Pottsdown, PA, USA). On the next day, brain sections were washed 3 times with 0.1% TX-100/PBS followed by two hours incubation of anti-rabbit-Alexa594 secondary antibody (Jackson Immunoresearch, West Grove, PA, USA). Tissue sections were imaged on an Olympus VS200 virtual slide microscope (Olympus, Tokyo, Japan).

### Open field test

Test mice were first habituated to an open field arena (40 × 40 cm) and allowed to freely explore for 20 minutes. Subsequently, mice were tested for 30 minutes on three occasions separated by 72-hour intervals. On the first test day, all mice received a vehicle injection prior to the open field test. On the second test day, mice were randomly assigned to receive either vehicle or LY354740 (10 mg/kg) 20 minutes prior to ketamine administration (10 mg/kg). On the third test day, treatment groups were crossed over. The open field test began immediately following the final injection on each test day and lasted 30 minutes. Total distance traveled was measured automatically using BIOBSERVE, and the first 20-minutes of the assay were quantified.

### Lentiviral production and purification

Murine mGluR2 (UniProtKB ID: Q14BI2) was cloned into pFUGW vector, with an added C-terminal mCherry tag. Specifically, the mCherry tag was inserted after amino acid positions 1-861 of Q14BI2, and before Q14BI2 positions 862-872, with GSGGGS linkers flanking the mCherry. The open-reading frame of this construct exactly matches the amino acid sequence of mGluR2-mCherry fusion receptor expressed in the *Grm2*^mCherry-FlpO^ mouse line. Lentivirus was produced by UNC-Chapel Hill NeuroTools Vector Core. HEK293T cells were transfected using a DNA-PEI protocol and grown in 10% FBS/1% L-glutamine/clear DMEM media. Forty-eight hours post transfection, supernatant is filtered followed by centrifugation at 28,000 x rpm for 90 minutes and then concentrated to 1 mL using another round of ultra centrifugation. Concentrated virus underwent purification using the anion exchange column^91^. Column was washed 5 times prior to use with 1X PBS at a flow rate of 5 mL/min, then washed with elution buffer at 5 mL/min for 5 min and equilibrated with starting buffer at 5 mL/min for 5 min. Concentrated virus vector was diluted in the starting buffer to a total volume of 10 mL and then loaded into a 10 mL syringe. Virus was eluted at a flow rate of 2.5 mL/min through the column. Column was then first washed with 0.2M NaCl, then 0.4M NaCl. Recovered virus from the 0.4M NaCl wash was concentrated by centrifugation at 28,000 x rpm for 90 minutes over a 20% sucrose cushion. The pellet was resuspended in 110 mL of 1X PBS and aliquoted 10 mL per tube. Virus titers (IU/mL) were determined by qPCR.

### Nanobody construct expression, purification and biotinylation

We systematically screened published anti-RFP nanobodies^66^ for their binding to mCherry with fluorescence size exclusion chromatography (FSEC) and found LaM6 to exhibit the best apparent antigen binding and performance in receptor purifications. LaM6 was inserted into the recently reported nanobody based purification platform^65^ (Addgene #149336). We added additional components to this system: a 3C site, photostable fluorescent protein mTFP1^92^, and the AlfaTag epitope^93^. These components were placed between the SUMO_Eu_ site and nanobody with generous GS linker permutations flanking each element. The mTFP1 tag enables low-detection limit fluorescence quantification of target after purification with fluorescence detection size exclusion chromatography (FSEC), whereas the AlfaTag provides an option for target immobilization after isolation. This construct was transformed in SHuffle® T7 Express Competent *E. coli* cells (NEB C3029J). A saturated overnight of the clone was used to inoculate 2.0L LB, which was grown at 37° C in LB media to an optical density at 600nm (OD_600_) of ∼1.0. Isopropyl β-D-1-thiogalactopyranoside was then added to the media at a final concentration of 1.0 mM and the temperature was dropped to 18° C for overnight expression. Cells were harvested via centrifugation at 4,000*g* for 10 minutes. Cell pellets were resuspended in TBS (20 mM Tris-HCl pH 8.0, 150 mM NaCl) supplemented with 1 mM MgCl_2_, 500 μM AEBSF, 1 μM E-64, 1 μM Leupeptin, 150 nM aprotinin, ∼1mg/mL DNase powder, and ∼1mg/mL egg white lysozyme powder. A total of 3 cycles of freeze thaw using liquid nitrogen was the performed. After freeze thaw and lysozyme mediated lysis, the cell lysate was clarified for 30 minutes at 20,000*g*. A final concentration of 20 mM imidazole was added to the clarified extract, which was then applied to 4.0 mL Ni-NTA resin (Takara His60 Ni Superflow resin #635660) pre-equilibrated with TBS+20 mM imidazole, for 1 hr at 4° C with gentle agitation. The resin was then collected in a gravity column and washed with 2x10 column volumes (CV) TBS+20 mM imidazole at room temperature. Nanobody was eluted from the resin by addition of 5x1 CV elution buffer (TBS+300mM imidazole). Fractions with the strongest color were then combined for nanobody biotinylation. This reaction was assembled on ice, with 1% v/v recombinantly expressed and purified BirA enzyme (Addgene #20857), 1 mM biotin, 10 mM adenosine triphosphate added. The reaction was carried out at 4° C with gentle agitation overnight, and terminated by sample desalting with a PD-10 column equilibrated with TBS. Final nanobody concentration was estimated with BCA and OD_280_, complete biotinylation was confirmed with a streptavidin gel-shift assay, and aliquots were snap frozen in liquid nitrogen for long term storage at - 80° C.

#### Detergent-purification of mGluR2 recombinantly expressed in HEK293T cells

mGluR2-mCherry lentivirus was added at a final MOI of ∼60 to HEK293T (1 million cells in a single well of a 6-well dish), and the infected cell pool was gradually expanded in Dulbecco’s Modified Eagle Medium media (4.5 g/L glucose, L-glutamine, sodium pyruvate, 10% v/v fetal bovine serum, 100 units/mL penicillin, and 100 μg/mL streptomycin) at 37° C, 5% CO_2_. Stable recombinant expression of mGluR2-mCherry was confirmed by visual inspection of membrane localized mCherry fluorescent signal through the duration of cell expansion. Once the cells reached >80% confluency at a scale of 50x15cm^2^ dishes, media was aspirated, ice-cold TBS supplemented with protease inhibitors (500 μM AEBSF, 1 μM E-64, 1 μM Leupeptin, 150 nM aprotinin) was added to the cells, and cells were detached from plates via scraping. The resuspended cells were collected into centrifuge bottles on ice, followed by gentle centrifugation. The cell pellets were then snap frozen for storage at -80° C.

On the day of purification, cell pellets were thawed on ice and resuspended in 20 mL lysis buffer (TBS supplemented with protease inhibitors, 0.5 mM EDTA, 1 μM LY354740, 1 μM JNJ-46281222). All of the following steps were carried out at 4° C. The thawed, resuspended cells were disrupted with 30 strokes in a traditional Dounce homogenizer. Membranes were collected with centrifugation at 20,000*g* for 30 minutes, after which were resuspended and homogenized in 10 mL lysis buffer. A final concentration of 2% w/v glyco-diosgenin (GDN; Anatrace) detergent was added to the homogenized membranes, and detergent extraction was carried out for 1 hr with gentle rotation. The solubilized membranes were then clarified via centrifugation at 20,000*g* for 30 minutes. During clarification, purification beads were prepared. In brief, biotinylated anti-mCherry nanobody construct was added to TBS equilibrated High Capacity Magne^®^ Streptavidin Beads (Promega V782A). A total of 2 mg biotinylated nanobody was loaded onto 1 mL bead slurry for 15 minutes at room temperature with gentle agitation (0.2 mL bead bed). The absence of visual color (mTFP1) in the supernatant confirmed efficient loading of the fluorescently tagged nanobody construct to the beads. The beads were then washed with 3x1.0 mL wash buffer (TBS, 0.02% w/v GDN, 1 μM LY354740, 1 μM JNJ-46281222) and added to the clarified, detergent solubilized cell lysate for 1.5 hr with gentle rotation. Washing was performed rapidly with 3x100 CV wash buffer (TBS, 0.02% GDN, 1 μM LY354740, 1 μM JNJ-46281222) using a magnetic rack. Target was eluted off of the beads with 0.5 mL elution buffer (wash buffer supplemented with 2% v/v purified SENP_EuB_ protease, Addgene construct 149333). Digestion completion was confirmed to be complete in ∼60s by the appearance of color (mTFP1) in the supernatant, indicating liberation of nanobody from beads through protease cleavage. The bead elution was then subjected to centrifugation at 20,000g for 15 minutes, followed by injection over a Superose 6 Increase 10/300 size exclusion chromatography column equilibrated with wash buffer. The peak fractions (∼2.5 mL) were pooled, concentrated to OD_280_=3.6 with a 100kDa MWCO spin column concentrator (final volume ∼10μL), and immediately used for cryo-EM sample vitrification.

### Detergent-purification of mGluR2 complexes from mouse whole brains

Homozygous *Grm2*^mCherry-FlpO/mCherry-Flpo^ mice (8-12 weeks old, mixed sex) were used for mGluR2 purifications. Mice were euthanized via cervical dislocation and decapitated. Whole brains were then quickly removed from the skull, placed in plastic 1.7mL Eppendorf tubes, and immediately snap frozen in liquid nitrogen. Whole brain tissue was stored at - 80° C until the day of purification.

One the day of purification, brains were removed from storage tubes and batched (50 per prep) over dry ice. The frozen brains were then quickly added to 70 mL ice cold homogenization buffer (TBS supplemented with 1.5 μM AEBSF, 3 μM E-64, 3 μM Leupeptin, 450 nM aprotinin, 0.5 mM EDTA, 10 μM LY354740, 10 μM JNJ-46281222; final volume ∼100 mL). The tissue was then immediately homogenized on ice with a Dounce homogenizer (pestle attached to a JoanLab overhead stirrer set to 1,500 rpm). After tissue was completely thawed on ice by repeated pressing of the rotating pestle onto the tissue in the Douncer, a total of 10 full strokes were performed. All of the following steps were performed at 4° C. A final concentration of 2% w/v GDN detergent powder was added to the crude homogenate in a large beaker, and detergent extraction was carried out with gentle magnetic bar stirring for 1 hr. The detergent solubilized crude homogenate was then clarified by centrifugation at 35,000*g* for 30 minutes. Clarified sample was collected in clean 50 mL conical tubes, with care taken to avoid a loose runny pellet of insoluble material. A total of 2 mL loaded (4 mg nanobody) magnetic streptavidin bead slurry was added to the sample, and bead binding was carried out with gentle rotation for 1 hr. The beads were rapidly washed on a magnetic rack with 10x50CV wash buffer (TBS, 0.02% GDN, 10 μM LY354740, 10 μM JNJ-46281222). Target was eluted with 2x2.5 mL elution buffer (wash buffer supplemented with 2% v/v SENP_EuB_ protease). The elution was concentrated to 0.5 mL with a 100kDa MWCO spin column concentrator, centrifuged for 15 minutes at 20,000*g*, and injected over SEC equilibrated with wash buffer. Peak fractions (∼2 mL) were pooled, concentrated to OD_280_ = 3.7 with a 100kDa MWCO spin column concentrator (final volume ∼10μL), and immediately used for cryo-EM sample vitrification.

### Mass spectrometry

#### Sample Preparation for all projects

Immunoprecipitated samples were subjected to SDS-PAGE and stained with Coomassie. Lanes (1cm) for each sample were excised, and the proteins were reduced with 5mM DTT for 30 min at 55 °C, alkylated with 15mM IAA for 45 min in the dark at room temperature, and in-gel digested with trypsin overnight at 37°C. Peptides were extracted, desalted with C18 spin columns (Pierce), and dried via vacuum centrifugation. Peptide samples were stored at -80°C until further analysis.

#### LC-MS/MS analysis for untargeted projects

The peptide samples were analyzed by liquid chromatography-tandem mass spectrometry (LC-MS/MS) in technical replicates using an Easy nLC 1200 coupled to a QExactive HF mass spectrometer (Thermo Scientific).

Samples were injected onto an Easy Spray PepMap C18 column (75 μm id × 25 cm, 2 μm particle size; Thermo Scientific) or an Aurora Ultimate TS column (75 μm id × 25 cm, 1.7 μm particle size; IonOpticks) and separated over a 45 min. method. The gradient for separation consisted of 5–38% mobile phase B at a 250 nl/min flow rate, where mobile phase A was 0.1% formic acid in water and mobile phase B consisted of 0.1% formic acid in 80% ACN. The QExactive HF was operated in data-dependent mode, where the 15 most intense precursors were selected for subsequent fragmentation. Resolution for the precursor scan (m/z 350–1750) was set to 60,000, with a maximum injection time of 100ms, and AGC set to 1e5. Following the full MS scan, a product ion scan was collected with a resolution set to 15,000, AGC set to 5e3, and dynamic exclusion set to 30s. The normalized collision energy was set to 27% for HCD. Peptide match was set to preferred, and precursors with unknown charge or a charge state of 1 and ≥ 8 were excluded.

#### LC-MS/MS analysis for PC1440 and PC1454

The peptide samples were analyzed by liquid chromatography-tandem mass spectrometry (LC-MS/MS) using an Ultimate3000 coupled to an Exploris480 mass spectrometer (Thermo Scientific). Samples were injected onto an IonOpticks Aurora series 2 C18 column (75 μm id × 15 cm, 1.6 μm particle size; IonOpticks) and separated over a 90-minute method. The gradient for separation consisted of 2–40% mobile phase B at a 250 nl/min flow rate, where mobile phase A was 0.1% formic acid in water and mobile phase B consisted of 0.1% formic acid in ACN. The Exploris480 was operated in data-dependent mode with a cycle time of 2s. Resolution for the precursor scan (m/z 375–1500) was set to 120,000, with AGC set to 300%. Following the full MS scan, a product ion scan was collected with a resolution set to 15,000, and normalized AGC set to 200%. The normalized collision energy was set to 30% for HCD.

Peptide match was set to preferred, and precursors with unknown charge or a charge state of 1 and ≥ 7 were excluded

#### Data analysis for PC1227, PC1233, PC1288, PC1352, PC1374, PC1454

Raw data files were searched against the Uniprot reviewed mouse database (containing 47,932 entries, downloaded January 2024), appended with a contaminants database, using the Sequest HT search engine node within Proteome Discoverer (v3.1, Thermo Fisher). Enzyme specificity was set to trypsin, up to two missed cleavage sites were allowed, methionine oxidation and N-terminus acetylation were set as variable modifications and cysteine carbamidomethylation was set as a static modification. The Minora node was used to extract label-free quantification (LFQ) intensities. A 1% peptide-level false discovery rate (FDR) and a 5% protein-level FDR was used to filter all data. Match between runs was enabled, and a minimum of two peptides was required for label-free quantitation using the LFQ intensities.

Data filtering, imputation, and statistical analysis were performed in Perseus software (version 1.6.14.0)^94^. Proteins with log2 fold change ≥ 1 and a p-value < 0.05 are considered significant.

#### Data Analysis PC1440

Raw data files were searched using the default LFQ-MBR workflow in Fragpipe (v22)^95^ against the Uniprot reviewed mouse database downloaded April 2025 (containing 17,230 sequences), appended with a contaminants database. Enzyme specificity was set to strict trypsin, up to two missed cleavage sites were allowed, methionine oxidation and N-terminus acetylation were set as variable modifications and cysteine carbamidomethylation was set as a static modification. IonQuant^96^ was used to extract label-free quantification (LFQ) intensities. Match between runs was enabled. Data filtering, imputation, and statistical analysis were performed in Perseus software (version 1.6.14.0)^94^. Proteins with log2 fold change ≥ 1 and a p-value < 0.05 are considered significant.

#### Targeted LC-MS PRM assay

Peptide separation was performed using a Thermo Scientific UltiMate 3000 RSLCnano. Samples were separated on an IonOpticks Aurora series 2 C18 column (75 μm id × 15 cm, 1.6 μm particle size; IonOpticks), heated to 50 °C, utilizing a 60-minute method. The gradient for separation consisted of 2–40% mobile phase B at a 250 nl/min flow rate, where mobile phase A was 0.1% formic acid in water and mobile phase B consisted of 0.1% formic acid in 80% ACN.

Data acquisition was performed on a Thermo Scientific Orbitrap Exploris 480 mass spectrometer equipped with a Nanospray Flex ion source. The mass spectrometer was operated in positive ion mode with a static spray voltage of 2100 V and an ion transfer tube temperature of 275 °C. Targeted MS2 scans were acquired for a predefined mass list over specific scheduled retention time windows. Precursor ions were isolated using a 1 m/z isolation window and fragmented using HCD with a normalized collision energy of 30%. The MS2 scans were acquired in profile mode at a resolution of 60,000. The AGC target was set to Standard with a maximum injection time of 120 ms utilizing 1 microscan. The RF lens amplitude was set to 40%. The complete targeted inclusion list containing all precursor m/z values and scheduling is provided in Table S1. Data was processed and peptide quantities estimated by comparing intensities to standard curves developed with synthetic peptide standards in Skyline (64-bit) 26.1^97^.

#### Standard Curve generation

The peptide synthesis was performed in the UNC Peptide Synthesis Core facility (RRID:SCR_017837), work was supported by the National Cancer Institute of the National Institutes of Health under award number P30CA016086. The synthetic peptides (Table S6) were accurately weighed and diluted to working concentrations (100 µM) based on expected formula weights of the TFA salts with 5% acetonitrile in water. The working stocks were further diluted by 4-fold serial dilution to obtain standard curve samples adequate to deliver between 0-7.8 fmol on column of each peptide. The serial-diluted peptides were used to confirm peptide identification, fragmentation, retention time and to generate standard curves used for estimating sample protein abundances in Skyline (64-bit) 26.1^97^.

### Cryo-EM grid preparation

All cryo-EM samples were prepared using a ThermoFisher Scientific Vitrobot Mark IV. Quantifoil gold R1.2/1.3 300 mesh grids were glow discharged (2 rounds of 25s, 15 mA; Pelco easiGlow) immediately prior to vitrification. A total of 3 μL purified receptor was applied to the glow discharged grid, blotting was carried out for 3s with a force of -10 at 4° C and 100% humidity, followed by plunge-freezing in liquid ethane.

### Cryo-EM data collection

#### recombinant mGluR2 1 μM LY354740 + 1 μM JNJ-46281222

A total of 8,684 movies from two distinct collection sessions (dataset A – 4,023; dataset B – 4,661), using two replicate grids, were collected on a Talos Artica transmission electron microscope (Thermo Fisher) operated at 200 kV, equipped with a K3 (Gatan) detector in counting mode (UNC Chapel Hill CryoEM Core facility). SerialEM^98^ was utilized with an in-house script^99^ to increase automated acquisition speed. A magnification of 45,000x with physical pixel size of 0.876 Å was utilized, with 60 frames collected over a 3.0s acquisition with an accumulated dose of ∼54 e^-^ Å^-2^. The defocus target range was -0.5 to -1.5 μm.

#### Brain-derived mGluR2 complexes 10 μM LY354740 + 10 μM JNJ-46281222

A total of 16,031 movies from three distinct collection sessions (dataset A - 5,778; dataset B - 4,616; Dataset C - 5,637) using two replicate grids were collected on a Titan Krios transmission electron microscope (Thermo Fisher) operated at 300 kV, equipped with a K3 (Gatan) detector in counting mode and a BioContinuum energy filter set to a slit width of 10 eV (Pacific Northwest Cryo-EM Consortium facility). A magnification of 81,000x was used, with a pixel size of 0.533 Å in super resolution mode (physical pixel size of 1.0655 Å). An exposure time of 2.7s over 60 frames was used, with an accumulated dose of ∼50 e^-^ Å^-2^. The defocus target range was -0.7 to -2.2 μm.

### Cryo-EM image processing

#### Recombinant mGluR2 1 μM LY354740 + 1 μM JNJ-46281222

In total, 8,684 movies were motion corrected using MotionCorr2^100^ implemented in Relion^101^. Micrographs were then imported to cryoSPARC^102^ for CTF estimation with CTFFIND4^103^. Curation criteria of <1000 Å astigmatism and <6 Å estimated CTF resolution were used to throw away bad images. For all of the data processing presented in this study, we devised a particle picking and initial classification strategy to maximize the number of good particles obtained into consensus reconstructions, which improves the signal used for downstream 3D classification based on finer features and more completely samples the particles present on the grids for population distribution estimates. All of the following steps are performed in cryoSPARC^102^. For this dataset, blob picking is performed over the 7,661 good micrographs (elliptical blob, minimum diameter 100 Å, maximum diameter 250 Å). After particle curation, particles were extracted with 4x Fourier binning (120px box size, 3.504 Å/px) and subjected to 2D classification. Good particle projections were then selected and subjected to ab initio and heterogenous refinement (class number “k”=4). In parallel, good 2D classes were selected and used as templates for picking over the entire dataset; and a selection of these good particles from a random subset of 100 micrographs were also used to train two Topaz^104^ picking models (ResNet8 and Conv63). Particles obtained from template picking and the 2 parallel Topaz runs were subjected to 2D classification, ab initio (k=4) and heterogenous refinement (k=4), all at 4x Fourier binning (120px box size, 3.504 Å/px). Good volumes corresponding to distinct conformational states obtained from these 4 parallel picking/classifications (blob, template, 2xTopaz) were then merged, and duplicates removed using a 50 Å inter-distance cutoff. A resulting total of 237,983 particles were obtained for the ACC state, and 209,160 particles for the inactive state. These stacks were reextracted at 2x Fourier binning for the active state (240px box size, 1.752 Å/px) or 3x Fourier binning (160px box size, 2.628 Å/px) for the inactive state. These stacks were further cleaned up with n=3 parallel runs of ab-initio/heterogenous refinements (k=3). Good volumes from the replicates were merged with duplicates removed, resulting in 194,339 particles for the active state and 152,946 particles for the inactive state.

For the active state stack, particles were transferred to Relion and subjected to Refine3D. Blush regularization^105^ was critical in obtaining higher quality reconstructions, and was utilized in all Relion refinements and 3D classifications described for this dataset. The 194,339 stack was transferred back to cryoSPARC for a local refinement using a global mask, resulting in a GS-FSC of 4.13 Å. Owing to the clear conformational state assignment (ACC) this stack was utilized in the population distribution analysis (particle count and reconstruction shown in Figure S10A). To further improve the density in the 7TM region, which is broken in this reconstruction, 3D classification in Relion was performed with global search (7.5°), tau_fudge=24, and k=3, for 32 iterations. One class featured cleared 7TM density, and particles corresponding to this class were merged from the last 4 iterations with duplicates removed. The resulting 94,681 particles were subjected to gold-standard refinement in Relion with an overall mask, followed by transfer to cryoSPARC for ECD (VFT+CRD) and 7TM focused refinements, resulting in 3.74 Å and 4.68 Å GS-FSC resolution, respectively (Figure S10B).

The 152,946 particles corresponding to the inactive state were transferred to Relion and re-extracted (120px box size, 3.504 Å/px). 3D classification was performed with k=4, tau_fudge=24, and global angular search (7.5°), for 40 iterations. Particles from the last 4 iterations corresponding to the class with the clearest features were merged with duplicates removed, resulting in 87,318 particles. This particle stack was subjected to gold-standard refinement in Relion, followed by transfer to cryoSPARC and re-extraction (240px box size, 1.752 Å/px). A local refinement using an overall mask resulted in a reconstruction to GS-FSC 6.43 Å. To improve the resolution, a ECD focused refinement was performed, resulting in a reconstruction to GS-FSC 4.96 Å resolution. This 87,318 particle subset was assigned as the RCO state and was used in the population distribution analysis (Figure S10B).

### Brain-isolated mGluR2 complexes 10 μM LY354740 + 10 μM JNJ-46281222

#### Obtaining consensus reconstructions

In total, 16,031 movies from three distinct collection sessions (5,778/4,616/5,637) were motion corrected using MotionCorr2^100^ implemented in Relion^101^, with 2x binning applied to the resulting corrected micrographs (1.066 Å/px). Micrographs were then imported to cryoSPARC^102^ for CTF estimation with CTFFIND4^103^. Curation criteria of <4.5 Å estimated CTF resolution were used to throw away bad images. Micrographs from the first two collection runs were first processed together in batch: these 9,697 good micrographs were subjected to a total of 5 parallel picking/initial classification runs: blob picking (circular), blob picking round 2 (elliptical), template picking, Topaz round 1 (ResNet8), Topaz round 2 (Conv63). Particles were then extracted (100px box, 4.26 Å/px) for 2D classifications and subsequent ab-initio/heterogenous refinements (k=4 or 5). Good volumes corresponding to distinct conformations were merged, with duplicates removed, resulting in 419,339 particles for the active state and 418,994 particles for the inactive state. These particle sets were further cleaned up with n=4 parallel runs of ab-initio/heterogenous refinements (k=3). Particles corresponding to good volumes were merged with duplicates removed, resulting in 290,258 particles for the active state and 330,086 particles for the inactive state. These particle subsets are referred to as “Dataset A+B consensus stacks”. Data from the third collection run (5,637 movies) was then processed in a similar manner: 5,503 good micrographs were subjected to a total of 4 parallel picking/initial classification runs: blob picking (elliptical), template picking, Topaz round 1 (ResNet8), Topaz round 2 (Conv63). Particles were then extracted (100px box, 4.26 Å/px) for 2D classifications and subsequent ab-initio/heterogenous refinements (k=4). Good volumes corresponding to distinct conformations were merged, with duplicates removed, resulting in 231,269 particles for the active state and 212,248 particles for the inactive state. These particle sets were further cleaned up with n=4 parallel runs of ab-initio/heterogenous refinements (k=3). Particles corresponding to good volumes were merged with duplicates removed, resulting in 153,659 particles for the active state and 162,032 particles for the inactive state. These particle subsets were merged with the “Dataset A+B consensus stacks” to obtain what are referred to as the “Dataset A+B+C consensus stacks”.

At this point, particle stacks were transferred to Relion with csparc2star.py^106^, where “Dataset A+B” and “Dataset A+B+C” stacks were processed independently. For active state stacks, particles were re-extracted without binning (400px box, 1.066 Å/px) and subjected to gold-standard Fourier Shell Correlation (GS-FSC) auto refinement in Relion, with an initial angular sampling of 7.5° (local sampling of 1.8°). Blush regularization^105^ was utilized in all Relion refinements and 3D classifications described for this dataset. Active state particles were then subjected to 3D classification with alignment (tau_fudge=12). For the “Dataset A+B” stack, 25 iterations were run with k=4. For the “Dataset A+B+C” stack, 35 iterations were run with k=5. In both cases, a single class had stronger G-protein and TMD features in the maps. The final 4 iterations corresponding to the stronger G-protein signal classes were merged and duplicates removed. For “Dataset A+B”, a total of 76,053 particles were refined to a GS-FSC resolution of 4.58 Å; for “Dataset A+B+C”, a total of 88,720 particles were refined to a GS-FSC resolution of 4.31 Å. Both of these stacks were merged and duplicates removed, resulting in 143,313 distinct particles – this set was refined to a GS-FSC resolution of 4.10 Å. Round of Bayesian polishing were performed, which improved the GS-FSC resolution to 3.95 Å. We refer to this particle subset as “ACC-G consensus.” This stack was then transferred to cryoSPARC, and masked local refinements were performed focused on the VFT/CRD, TMD, and G-protein elements, leading to GS-FSC resolutions of 3.31 Å, 3.22 Å and 3.52 Å, respectively. To further improve the G-protein density, focused 3D classifications without alignment in cryoSPARC were performed in triplicate (k=3). Classes corresponding to stronger G-protein signal were merged and duplicates removed, and this subset (69,978 particles) was subjected to a local refinement resulting in a reconstruction of 3.43 Å resolution and clear side-chain densities for the Gα and Gβ subunits. This reconstruction is referred to as the “focused GoA subset”, which enabled straightforward model building for these complex components. The “ACC-G” consensus subset was then removed from the starting “Dataset A+B+C” active state subset, and these remaining particles (298,996 particles) were refined in Relion to an overall resolution of 3.32 Å. Owing to the lack of strong G-protein signal in these leftovers, this particle subset is referred to as the “ACC consensus”.

The inactive state particles were transferred to Relion and re-extracted with 2x binning (200px box, 2.13 Å/px) and subjected to 3D classification with alignment (tau_fudge=12). For the “Dataset A+B” inactive subset, 25 iterations were run with k=3; for “Dataset A+B+C” inactive subset 40 iterations were run with k=5. Upon careful inspection of the volumes, two distinct conformational states were apparent – one pseudo C2 symmetric reconstruction in which both VFTs are open (ROO) and one C1 reconstruction in which one VFT is open, while the other subunit’s VFT is apparently closed (RCO). Particles corresponding to classes for either ROO or RCO with the clearest features were combined from the last 4 iterations per classification run, with duplicates removed. These stacks were then combined across the two parallel classification jobs (“Dataset A+B” and “Dataset A+B+C”) for either ROO or RCO, with duplicates removed. This resulted in 148,457 distinct particles for ROO and 293,405 distinct particles for RCO. The particle stacks were re-extracted without binning (400px box, 1.066 Å/px). The ROO stack was subjected to refinement with C2 symmetry applied, yielding a reconstruction of 3.81 Å overall resolution. For RCO, the particles were subjected to a refinement with C1 applied, resulting in a reconstruction of 4.14 Å overall resolution. This subset is referred to as the “ROO consensus” – the RCO subset was further classified into two distinct substates, which is described in the next section.

#### Venus flytrap domain focused classifications in the ACC states

In parallel to the aforementioned classification approaches to separate particles based on their overall conformational state, 3D classifications focused on the VFTs were performed. First, a prominent non-protein density is present within the 7TM of one subunit of the mGluR dimer in both the ACC and ACC-G consensus reconstructions. In ACC-G, the ligand is present within the 7TM of G-protein coupled receptor subunit; the 7TM of the non-G protein coupled subunit is devoid of this signal. The size and shape of the volume is consistent with the ago-PAM JNJ462 that was added during the purification (Figure S4E). Owing to the high degree of subtype selectivity this ago-PAM exhibits (Figure S3A), and the fact that mGluR2-selective PAMs typically signal asymmetrically *in cis* from structural^14,16,17^ and functional^48^ standpoints, the subunits containing JNJ462 can be confidently assigned as subtype R2. The side chain densities, loop conformations, and a R2-specific N-linked glycosylation site support this assignment (Figure S4E). Upon closer inspection of the mGluR subunit of the dimer in which JNJ462 signal is absent within the 7TM, broken densities in divergent loop regions of the VFT, weak or non-existent R2 specific N-linked glycan, and spurious side chain densities at positions variable amongst the mGluR subtypes are apparent, all of which indicate compositional heterogeneity (Figure S4E). For these reasons, the following classification efforts were focused on breaking the ambiguity present in this subunit.

The “Dataset A+B+C” starting active particle stack (443,222 particles) was subjected to Bayesian polishing and refinement in Relion to improve the overall resolution of the reconstruction to 3.47 Å. The particle stack was then transferred to cryoSPARC and subjected to VFT masked local refinements and global CTF refinement (per optics group), resulting in a resolution for the VFT dimer to 2.82 Å. The relatively higher resolution of this reconstruction allowed us to perform 3D classification without alignment in cryoSPARC with a focused mask around the heterogenous subunit’s VFT, using signal to 3.5 Å during the classification runs. A total of 9 classification jobs were ultimately performed – n=3 replicates for k=4,5, or 6 job settings. Clean volumes corresponding to R2 or R3 subtypes were combined first within the triplicate runs, duplicates removed, and any particles that came up in both R2 or R3 classes across the replicates were removed with the Particle Sets tool (intersects removed). Removing intersects during the classification procedure led to cleaner separation of R2 and R3. This procedure was then repeated to combine R2 or R3 stacks from the k=4,5 and 6 classification runs (merge and remove duplicates, remove intersecting particles between R2 and R3 stacks). The resulting 139,681 particles corresponding to the R2R2 homodimer, 150,502 particles corresponding to the R2R3 heterodimer, and 153,039 particles with signal features at the heterogenous position that could not be clearly classified (R2/RX), were subjected to local refinements yielding final resolutions for 2.94 Å, 2.97 Å and 3.38 Å, respectively. Validation of R2 or R3 subunit assignment in these reconstructions by inspection of finer map details are summarized in the later section (Figure S6).

#### Obtaining ensembles of ACC and ACC-G mGluR assemblies

Having separated the active state particles (443,222 particles starting, from all data) based on presence of endogenous G-protein, in parallel with VFT focused classifications to separate based on subtype composition, intersecting particle pick locations from both approaches were identified to reconstruct distinct compositional species for each conformational state (remove duplicates tool in cryoSPARC, 50 Å inter-particle distance). Angular priors were retained from the ACC-G consensus stack for the ternary complexes, which greatly improved the map quality in the 7TM and G-protein regions in the resulting reconstructions^107^. For the ACC state, intersects were dropped at random which did not impact the overall reconstruction quality. Masked local refinements were then performed on these particle subsets, focused on the VFT/CRD, TMD, and G-protein elements (Figure S5A). A total of six particle subsets were obtained from these approaches: R2R2-ACC-G (46,526 ptcls), R2R3-ACC-G (49,250 ptlcs), R2RX-ACC-G (45,349 ptlcs), R2R2-ACC (90,200 ptlcs), R2R3-ACC (96,795 ptlcs), R2RX-ACC (100,938 ptlcs). Masks used for final local refinements, and GS-FSC resolutions are shown in Figure S5.

#### Venus flytrap domain focused classifications in the ROO state

The particle stack corresponding to the ROO consensus was transferred to cryoSPARC. Symmetry expansion (C2) was then performed, resulting in 296,914 expanded particles. A mask was generated over a single VFT subunit (the asymmetric unit in symmetry expanded real space) and used for local refinement. The resulting reconstruction (3.55 Å) featured smeary signal and broken loops, indicating conformational and/or compositional heterogeneity. Clean volumes with clear signal features of subtype R2 were obtained when using 5 or 6 classes for cryoSPARC focused 3D classification, using signal to 4.5 Å and the asymmetric VFT mask. A total of 6 classification jobs were ultimately performed: n=3 replicates for k=5 or 6 job settings. R2 corresponding classes from replicates were combined and intersected against classes with apparently mixed signal features. This procedure resulted in 72,946 expanded particles, which were subjected to an additional round of local refinement (3.44 Å). This focused reconstruction was of sufficient quality to enable straightforward model building of the VFT in the open configuration, and placement of the orthosteric agonist LY354740 into this conformational state. Further classification of the leftover particles, in the attempts to identify other subtypes in this conformer, proved to be recalcitrant, indicating additional unresolvable heterogeneity.

The remove duplicates tool was used to revert symmetry expansion. Parent particles that contained two copies of R2 assigned symmetry expanded particles were identified (10,397 particles) – corresponding to R2R2 homodimers. These particles were then subjected to local refinement with a full mask and C2 applied, which is referred to as the final “R2R2 ROO” reconstruction (3.74 Å). Similarly, instances in which parent particles contained one copy of an R2 subunit were identified (53,044 particles) – a local refinement using a full mask was performed, leading to the “R2RX ROO” reconstruction (4.02 Å).

#### Venus flytrap domain focused classifications in the RCO state

The particle stack corresponding to RCO was transferred to cryoSPARC. The closed VFT was masked and subjected to 3D classification with signal to 6 Å. Triplicate runs were performed with 3 classes. Two distinct substates were apparent in these classification runs – a fully closed VFT and partially closed VFT. The partially closed VFT is referred to as state C_i_ and the fully closed VFT as state C. Classes for either state were merged, duplicates removed, with intersects removed between C_i_ and C, resulting in 77,197 and 207,320 particles, respectively. These stacks were subjected to local refinements, yielding local reconstructions of 4.03 Å and 3.59 Å resolution, respectively. While the “C_i_” state does not contain signal features that allow clear subtype assignment, the signal in the “C” state reconstruction is consistent with subtype R2. Subsequent classification of the “C” state did not reveal any obvious signs of other mGluR subtypes; the “C_i_” is relatively lowly populated, which prohibited subsequent classification.

The open VFT in the RCO stack was also masked and subjected to 3D classification in parallel. Signal to 4.5 Å was utilized with k=6, giving the cleanest class corresponding to subtype R2. Triplicate classification jobs were performed, the clear R2 classes from replicates were merged, duplicates removed and intersected against classes exhibiting mixed features. A total of 77,803 particles were obtained with this approach, and a local refinement was performed resulting in a reconstruction of 3.98 Å, which was used to confirm assignment as subtype R2.

To obtain all possible reconstructions in which at least one subunit of the dimer was confidently assigned as subtype R2, intersecting particles between the open and closed VFT classification runs were identified. This resulted in three distinct reconstructions: “RXR2 RC_i_O” (19,870 particles; 6.41 Å resolution), “R2RX RCO” (151,702 particles; 4.36 Å resolution), and “R2R2 RCO” (55,168 particles; 4.58 Å resolution) (Figure S5E, S5H).

### Particle distribution analysis

A majority of the aforementioned processing approaches we employed for this study involve running replicates of particle picking and 3D classifications, to compensate for the intrinsic inefficiency of cryo-EM image processing. This results in some particle projections getting assigned into more than one distinct final particle subset. To account for this in our population analysis, we identified distinct particle projections for all final particle subsets using the remove duplicates tool in cryoSPARC. We also calculated redundancy values, which reflects the percentage of the particle projections that are present in two or more final particle subsets (Extended Data Table 5). For the recombinant mGluR2 dataset, we found a 5.6% redundancy in the final assigned particles in the dataset (Extended Data Table 5). For the brain-isolated mGluR2 dataset, we found a 26.4% redundancy across the consensus stacks and a 13.1% redundancy in the final distinct assemblies (Extended Data Table 5). Only distinct particle projections were utilized in calculating particle population distributions (Figure S10E-G, Extended Data Table 5), and subtype mole fraction estimates (Figure 2B, Extended Data Table 5).

### Model building, refinement and validation

High-resolution crystal structures (PDB IDs 4XAS and 6B7H) were used as starting models for the VFT. Owing to the lower local resolution of the CRD, an AlphaFold model was used as starting coordinates for this domain (AF-Q14BI2-F1). The starting model for the 7TM domain was obtained from a SwissModel^108^ template prediction of a previously solved mGluR2 cryo-EM structure (PDB ID 7MTS). All manual adjustments were performed in Coot^109^, hydrogens were added with MolProbity to aid in real-space refinements^110^, and real-space refinements were carried out in Phenix^111^.

### OMIT map calculation

Phenix^111^ was used to generate OMIT maps show in Figure S12. In brief, HETATM were deleted from PDBs 4XAQ, 5CNI (mGluR2 VFT structures), 5CNM, 4XAR, 5CNK, 6B7H (mGluR3 VFT structures). These protein-only coordinates, and deposited structure factors from the PDB, were used to calculate mFo-DFc difference maps. Figures of the mFo-DFc maps and deposited coordinates were generated in ChimeraX^112^.

### Pharmacology assays

#### BRET2 (TRUPATH G_oA_)

HEK293 cells were maintained in Dulbecco’s modified Eagle medium (DMEM) supplemented with 10% (v/v) fetal bovine serum (FBS), 10,000 U ml⁻¹ penicillin and 10 mg ml⁻¹ streptomycin at 37 °C in 5% CO₂. For each experiment, 2–3 × 10⁶ cells were plated per 6-cm dish. After 24 h, cells were transfected with TransIT-2020 (Mirus Bio; 3 µl µg⁻¹ DNA). For each condition, 250 ng of each plasmid—mouse Grm2 (mGluR2) *or* mouse Grm3 (mGluR3), human SLC1A1 (EAAT3), GαoA-RLuc8, Gβ₃ and Gγ₈-GFP2—was combined in Opti-MEM (Thermo Fisher Scientific) to form the transfection complex. Twenty-four hours later, cells were detached with 0.05% trypsin–EDTA, resuspended in Basal Medium Eagle (BME) without glutamine supplemented with 1% dialyzed FBS (dFBS), and seeded at 1 × 10⁴ cells per well into poly-L-lysine-coated white 384-well plates (Greiner Bio-One).

For BRET2 agonist assay, after a further 24 h, plate bottoms were sealed with white backing tape (Revvity), medium was removed and wells were washed once with 20 µl assay buffer (Locke’s buffer for mGluR2 or Locke-Cl⁻ buffer for mGluR3, each supplemented with 0.1% bovine serum albumin; adapted from^113^). Locke’s buffer contained 154 mM NaCl, 5.6 mM KCl, 1.3 mM CaCl₂, 1 mM MgCl₂, 3.6 mM NaHCO₃, 5.6 mM glucose, and 20 mM HEPES (pH 7.4), whereas Locke-Cl⁻ buffer contained equimolar sodium gluconate and potassium gluconate in place of NaCl and KCl. Serial 3-fold stock dilutions of L-glutamate, LY354740, LY2794193, or JNJ-46281222 were prepared in assay buffer to final in-well concentrations up to 300 µM. To each well, 20 µl buffer containing 5 µM coelenterazine-400a (Nanolight Technologies) and 10 µl ligand solution were added, followed by incubation for 15 min at room temperature before reading.

For BRET2 chloride-sensitivity assay, assay buffers spanning 2–162 mM chloride were generated by mixing Locke’s buffer (162 mM Cl⁻) with Locke-Cl⁻ buffer (2 mM Cl⁻) in defined ratios. For these experiments, wells were washed with 20 µl of the designated chloride buffer concentration and subsequently reloaded with 20 µl of the same buffer. After equilibration at 37 °C for 15 min, agonists diluted in the same buffer containing coelenterazine-400a (final 5 µM) were added and plates were incubated for a further 15 min at room temperature before reading.

Emission at 395 nm (donor) and 510 nm (acceptor) was measured on a PHERAstar FSX plate reader (BMG Labtech). The BRET2 ratio was calculated as GFP2 (acceptor) emission divided by RLuc8 (donor) emission.

#### BRET 2 (TRUPATH G_oA_) Data analysis

BRET2 signals reporting Gα_oA_ dissociation were expressed as donor/acceptor emission ratios, normalized to the glutamate control E_max_ in the absence of chloride, and analyzed in Prism v10 (GraphPad). To quantify chloride-dependent allosteric effects on agonist responses, concentration-response datasets collected at multiple chloride concentrations were fit to the allosteric operational model of agonism (Black-Leff-Ehlert^81,114,115^).

Within this framework, the affinity cooperativity factor (α) describes how the modulator (chloride) alters the apparent binding affinity of the orthosteric agonist, whereas the efficacy cooperativity factor (β) describes how the modulator alters agonist efficacy. Values of α > 1 or β > 1 indicate positive cooperativity in affinity or efficacy, respectively; values between 0 and 1 indicate negative cooperativity.

The orthosteric and allosteric affinity terms (K_A_ and K_B_) were constrained to empirically determined apparent potencies measured under reference conditions (no chloride for K_A_; no orthosteric agonist for K_B_). When chloride produced little or no change in maximal response, β was constrained at 1. The remaining parameters—α, K_B_ (if not fixed as above), orthosteric efficacy (τ_A_), and allosteric efficacy (τ_B_)—were shared globally across curves within a dataset to reflect common system properties assumed by the model.

E_max_ is a system parameter representing the maximal possible response. Because the largest observed effect does not necessarily imply full saturation in this assay, we allowed E_max_ to be constrained between 100% and 200%, which improved the fit quality as reflected by the R² value. All parameter fitting and statistical tests pertinent to these analyses are provided in the source data.

#### GloSensor cAMP assay (G_i_ pathway)

G_i_-dependent modulation of intracellular cAMP was quantified with the GloSensor-22F luciferase-based cAMP reporter (Promega). HEK293 cells were transiently transfected with the GloSensor plasmid together with mouse Grm2 (mGluR2) or an mGluR2–mCherry fusion construct, then seeded onto poly-L-lysine-coated white, clear-bottom 384-well plates at 1.0 × 10⁴ cells per well in 40 µl BME without glutamine containing 1% dFBS. After 24 h, the medium was replaced with 20 µl per well of assay buffer (Hank’s balanced salt solution (HBSS) supplemented with 20 mM HEPES, pH 7.4, and 0.1% BSA) containing serial dilutions of L-glutamate together with D-luciferin (GoldBio) at a fixed 3 mM, and plates were incubated for 15 min at room temperature. Isoproterenol was then added (10 µl per well; 100 nM final) to activate endogenous β₂-adrenergic receptors and elevate cAMP via Gs. Luminescence was recorded after a further 15 min using a SpectraMax L microplate luminometer (Molecular Devices).

### Cell line statement

Cell lines were originally obtained from ATCC. No commonly misidentified cell lines were used in this study.

## Notes

### Summary of Updates

This version has been updated to include additional validation data (mouse line validation, additional pharmacology, quantitative LC-MS/MS). Authors contributing to these additional data were added. Main text, main figures, and supplemental materials were updated to incorporate the new data and additional analyses.

## References

1. Twomey, E.C., and Sobolevsky, A.I. (2018). Structural Mechanisms of Gating in Ionotropic Glutamate Receptors. Biochemistry 57, 267–276. 10.1021/acs.biochem.7b00891.

2. Hansen, K.B., Wollmuth, L.P., Bowie, D., Furukawa, H., Menniti, F.S., Sobolevsky, A.I., Swanson, G.T., Swanger, S.A., Greger, I.H., Nakagawa, T., et al. (2021). Structure, Function, and Pharmacology of Glutamate Receptor Ion Channels. Pharmacol Rev 73, 1469–1658. 10.1124/pharmrev.120.000131.

3. Ferraguti, F., and Shigemoto, R. (2006). Metabotropic glutamate receptors. Cell Tissue Res 326, 483–504. 10.1007/s00441-006-0266-5.

4. Niswender, C.M., and Conn, P.J. (2010). Metabotropic Glutamate Receptors: Physiology, Pharmacology, and Disease. Annual Review of Pharmacology and Toxicology 50, 295–322. 10.1146/annurev.pharmtox.011008.145533.

5. Wickman, K.D., Iñiguez-Lluhi, J.A., Davenport, P.A., Taussig, R., Krapivinsky, G.B., Linder, M.E., Gilman, A.G., and Clapham, D.E. (1994). Recombinant G-protein βγ-subunits activate the muscarinic-gated atrial potassium channel. Nature 368, 255–257. 10.1038/368255a0.

6. Whorton, M.R., and MacKinnon, R. (2013). X-ray structure of the mammalian GIRK2–βγ G-protein complex. Nature 498, 190–197. 10.1038/nature12241.

7. Ikeda, S.R. (1996). Voltage-dependent modulation of N-type calcium channels by G-protein β γsubunits. Nature 380, 255–258. 10.1038/380255a0.

8. Herlitze, S., Garcia, D.E., Mackie, K., Hille, B., Scheuer, T., and Catterall, W.A. (1996). Modulation of Ca2+ channels βγ G-protein py subunits. Nature 380, 258–262. 10.1038/380258a0.

9. Jarvis, S.E., Magga, J.M., Beedle, A.M., Braun, J.E.A., and Zamponi, G.W. (2000). G Protein Modulation of N-type Calcium Channels Is Facilitated by Physical Interactions between Syntaxin 1A and Gβγ *. Journal of Biological Chemistry 275, 6388–6394. 10.1074/jbc.275.9.6388.

10. Eitel, A.R., Mueller, B.K., Kaya, A.I., Young, M., Cassada, J.B., Bell, E.W., Schnitkey, L., Zurawski, Z., Yim, Y.Y., Zhou, Q., et al. (2025). Molecular basis for Gβγ-SNARE-mediated inhibition of synaptic vesicle fusion. Journal of Biological Chemistry 301, 110377. 10.1016/j.jbc.2025.110377.

11. Sladeczek, F., Pin, J.-P., Récasens, M., Bockaert, J., and Weiss, S. (1985). Glutamate stimulates inositol phosphate formation in striatal neurones. Nature 317, 717–719. 10.1038/317717a0.

12. Nicoletti, F., Meek, J.L., Iadarola, M.J., Chuang, D.M., Roth, B.L., and Costa, E. (1986). Coupling of Inositol Phospholipid Metabolism with Excitatory Amino Acid Recognition Sites in Rat Hippocampus. Journal of Neurochemistry 46, 40–46. 10.1111/j.1471-4159.1986.tb12922.x.

13. Koehl, A., Hu, H., Feng, D., Sun, B., Zhang, Y., Robertson, M.J., Chu, M., Kobilka, T.S., Laeremans, T., Steyaert, J., et al. (2019). Structural insights into the activation of metabotropic glutamate receptors. Nature 566, 79–84. 10.1038/s41586-019-0881-4.

14. Seven, A.B., Barros-Álvarez, X., de Lapeyrière, M., Papasergi-Scott, M.M., Robertson, M.J., Zhang, C., Nwokonko, R.M., Gao, Y., Meyerowitz, J.G., Rocher, J.-P., et al. (2021). G-protein activation by a metabotropic glutamate receptor. Nature 595, 450–454. 10.1038/s41586-021-03680-3.

15. Lin, S., Han, S., Cai, X., Tan, Q., Zhou, K., Wang, D., Wang, X., Du, J., Yi, C., Chu, X., et al. (2021). Structures of Gi-bound metabotropic glutamate receptors mGlu2 and mGlu4. Nature 594, 583–588. 10.1038/s41586-021-03495-2.

16. Wang, X., Wang, M., Xu, T., Feng, Y., Shao, Q., Han, S., Chu, X., Xu, Y., Lin, S., Zhao, Q., et al. (2023). Structural insights into dimerization and activation of the mGlu2–mGlu3 and mGlu2–mGlu4 heterodimers. Cell Res 33, 762–774. 10.1038/s41422-023-00830-2.

17. Huang, W., Jin, N., Guo, J., Shen, C., Xu, C., Xi, K., Bonhomme, L., Quast, R.B., Shen, D.-D., Qin, J., et al. (2024). Structural basis of orientated asymmetry in a mGlu heterodimer. Nat Commun 15, 10345. 10.1038/s41467-024-54744-7.

18. Krishna Kumar, K., Wang, H., Habrian, C., Latorraca, N.R., Xu, J., O’Brien, E.S., Zhang, C., Montabana, E., Koehl, A., Marqusee, S., et al. (2024). Stepwise activation of a metabotropic glutamate receptor. Nature 629, 951–956. 10.1038/s41586-024-07327-x.

19. Wen, T., Du, M., Lu, Y., Jia, N., Lu, X., Liu, N., Chang, S., Zhang, X., Shen, Y., and Yang, X. (2025). Molecular basis of β-arrestin coupling to the metabotropic glutamate receptor mGlu3. Nat Chem Biol, 1–8. 10.1038/s41589-025-01858-8.

20. Strauss, A., Gonzalez-Hernandez, A.J., Lee, J., Abreu, N., Selvakumar, P., Salas-Estrada, L., Kristt, M., Arefin, A., Huynh, K., Marx, D.C., et al. (2024). Structural basis of positive allosteric modulation of metabotropic glutamate receptor activation and internalization. Nat Commun 15, 6498. 10.1038/s41467-024-50548-x.

21. Fang, W., Yang, F., Xu, C., Ling, S., Lin, L., Zhou, Y., Sun, W., Wang, X., Liu, P., Rondard, P., et al. (2022). Structural basis of the activation of metabotropic glutamate receptor 3. Cell Res 32, 695–698. 10.1038/s41422-022-00623-z.

22. Zhao, J., Deng, Y., Xu, Z., Xu, C., Zhao, C., Li, Z., Sun, H., Tian, X., Song, Y., Cimadevila, M., et al. (2025). Structural characterization of five functional states of metabotropic glutamate receptor 8. Molecular Cell 85, 3460–3473.e6. 10.1016/j.molcel.2025.08.019.

23. Habrian, C.H., Levitz, J., Vyklicky, V., Fu, Z., Hoagland, A., McCort-Tranchepain, I., Acher, F., and Isacoff, E.Y. (2019). Conformational pathway provides unique sensitivity to a synaptic mGluR. Nat Commun 10, 5572. 10.1038/s41467-019-13407-8.

24. Liauw, B.W.-H., Afsari, H.S., and Vafabakhsh, R. (2021). Conformational rearrangement during activation of a metabotropic glutamate receptor. Nat Chem Biol 17, 291–297. 10.1038/s41589-020-00702-5.

25. Latorraca, N.R., Sabaat, S., Habrian, C.H., Bleier, J., Stanley, C., Kinz-Thompson, C.D., Marqusee, S., and Isacoff, E.Y. (2025). Domain coupling in activation of a family C GPCR. Nat Chem Biol, 1–11. 10.1038/s41589-025-01895-3.

26. Vafabakhsh, R., Levitz, J., and Isacoff, E.Y. (2015). Conformational dynamics of a class C G-protein-coupled receptor. Nature 524, 497–501. 10.1038/nature14679.

27. Moghaddam, B., and Adams, B.W. (1998). Reversal of Phencyclidine Effects by a Group II Metabotropic Glutamate Receptor Agonist in Rats. Science 281, 1349–1352. 10.1126/science.281.5381.1349.

28. Krystal, J.H., Abi-Saab, W., Perry, E., D’Souza, D.C., Liu, N., Gueorguieva, R., McDougall, L., Hunsberger, T., Belger, A., Levine, L., et al. (2005). Preliminary evidence of attenuation of the disruptive effects of the NMDA glutamate receptor antagonist, ketamine, on working memory by pretreatment with the group II metabotropic glutamate receptor agonist, LY354740, in healthy human subjects. Psychopharmacology (Berl) 179, 303–309. 10.1007/s00213-004-1982-8.

29. Moghaddam, B. (2004). Targeting metabotropic glutamate receptors for treatment of the cognitive symptoms of schizophrenia. Psychopharmacology 174, 39–44. 10.1007/s00213-004-1792-z.

30. Moghaddam, B., and Javitt, D. (2012). From Revolution to Evolution: The Glutamate Hypothesis of Schizophrenia and its Implication for Treatment. Neuropsychopharmacol 37, 4–15. 10.1038/npp.2011.181.

31. Saini, S.M., Mancuso, S.G., Mostaid, M.S., Liu, C., Pantelis, C., Everall, I.P., and Bousman, C.A. (2017). Meta-analysis supports GWAS-implicated link between GRM3 and schizophrenia risk. Transl Psychiatry 7, e1196–e1196. 10.1038/tp.2017.172.

32. Dogra, S., and Conn, P.J. (2021). Targeting metabotropic glutamate receptors for the treatment of depression and other stress-related disorders. Neuropharmacology 196, 108687. 10.1016/j.neuropharm.2021.108687.

33. Roth, B.L. (2019). Molecular pharmacology of metabotropic receptors targeted by neuropsychiatric drugs. Nat Struct Mol Biol 26, 535–544. 10.1038/s41594-019-0252-8.

34. Adams, D.H., Kinon, B.J., Baygani, S., Millen, B.A., Velona, I., Kollack-Walker, S., and Walling, D.P. (2013). A long-term, phase 2, multicenter, randomized, open-label, comparative safety study of pomaglumetad methionil (LY2140023 monohydrate) versus atypical antipsychotic standard of care in patients with schizophrenia. BMC Psychiatry 13, 143. 10.1186/1471-244X-13-143.

35. Adams, D.H., Zhang, L., Millen, B.A., Kinon, B.J., and Gomez, J.-C. (2014). Pomaglumetad Methionil (LY2140023 Monohydrate) and Aripiprazole in Patients with Schizophrenia: A Phase 3, Multicenter, Double-Blind Comparison. Schizophrenia Research and Treatment 2014, 758212. 10.1155/2014/758212.

36. Oosterlaken, M., Rogliardo, A., Lipina, T., Lafon, P.-A., Tsitokana, M.E., Keck, M., Cahuzac, H., Prieu-Sérandon, P., Diem, S., Derieux, C., et al. (2025). Nanobody therapy rescues behavioural deficits of NMDA receptor hypofunction. Nature, 1–9. 10.1038/s41586-025-09265-8.

37. Downing, A.M., Kinon, B.J., Millen, B.A., Zhang, L., Liu, L., Morozova, M.A., Brenner, R., Rayle, T.J., Nisenbaum, L., Zhao, F., et al. (2014). A double-blind, placebo-controlled comparator study of LY2140023 monohydrate in patients with schizophrenia. BMC Psychiatry 14, 351. 10.1186/s12888-014-0351-3.

38. Litman, R.E., Smith, M.A., Doherty, J.J., Cross, A., Raines, S., Gertsik, L., and Zukin, S.R. (2016). AZD8529, a positive allosteric modulator at the mGluR2 receptor, does not improve symptoms in schizophrenia: A proof of principle study. Schizophrenia Research 172, 152–157. 10.1016/j.schres.2016.02.001.

39. Grabb, M.C., and Potter, W.Z. (2022). Central Nervous System Trial Failures: Using the Fragile X Syndrome–mGluR5 Drug Target to Highlight the Complexities of Translating Preclinical Discoveries Into Human Trials. Journal of Clinical Psychopharmacology 42, 234. 10.1097/JCP.0000000000001553.

40. Conn, P.J., Lindsley, C.W., Meiler, J., and Niswender, C.M. (2014). Opportunities and challenges in the discovery of allosteric modulators of GPCRs for treating CNS disorders. Nat Rev Drug Discov 13, 692–708. 10.1038/nrd4308.

41. Jeffrey Conn, P., Christopoulos, A., and Lindsley, C.W. (2009). Allosteric modulators of GPCRs: a novel approach for the treatment of CNS disorders. Nat Rev Drug Discov 8, 41–54. 10.1038/nrd2760.

42. McCullock, T.W., and Kammermeier, P.J. (2021). The evidence for and consequences of metabotropic glutamate receptor heterodimerization. Neuropharmacology 199, 108801. 10.1016/j.neuropharm.2021.108801.

43. Moreno Delgado, D., Møller, T.C., Ster, J., Giraldo, J., Maurel, D., Rovira, X., Scholler, P., Zwier, J.M., Perroy, J., Durroux, T., et al. (2017). Pharmacological evidence for a metabotropic glutamate receptor heterodimer in neuronal cells. Elife 6, e25233. 10.7554/eLife.25233.

44. Yin, S., Noetzel, M.J., Johnson, K.A., Zamorano, R., Jalan-Sakrikar, N., Gregory, K.J., Conn, P.J., and Niswender, C.M. (2014). Selective Actions of Novel Allosteric Modulators Reveal Functional Heteromers of Metabotropic Glutamate Receptors in the CNS. J. Neurosci. 34, 79–94. 10.1523/JNEUROSCI.1129-13.2014.

45. Meng, J., Xu, C., Lafon, P.-A., Roux, S., Mathieu, M., Zhou, R., Scholler, P., Blanc, E., Becker, J.A.J., Le Merrer, J., et al. (2022). Nanobody-based sensors reveal a high proportion of mGlu heterodimers in the brain. Nat Chem Biol 18, 894–903. 10.1038/s41589-022-01050-2.

46. Lee, J., Munguba, H., Gutzeit, V.A., Singh, D.R., Kristt, M., Dittman, J.S., and Levitz, J. (2020). Defining the Homo- and Heterodimerization Propensities of Metabotropic Glutamate Receptors. Cell Reports 31, 107605. 10.1016/j.celrep.2020.107605.

47. Habrian, C., Latorraca, N., Fu, Z., and Isacoff, E.Y. (2023). Homo- and hetero-dimeric subunit interactions set affinity and efficacy in metabotropic glutamate receptors. Nat Commun 14, 8288. 10.1038/s41467-023-44013-4.

48. Lin, X., Provasi, D., Niswender, C.M., Asher, W.B., and Javitch, J.A. (2024). Elucidating the molecular logic of a metabotropic glutamate receptor heterodimer. Nat Commun 15, 8552. 10.1038/s41467-024-52822-4.

49. Philibert, C.E., Disdier, C., Lafon, P.-A., Bouyssou, A., Oosterlaken, M., Galant, S., Pizzoccaro, A., Tuduri, P., Ster, J., Liu, J., et al. (2024). TrkB receptor interacts with mGlu _2_ receptor and mediates antipsychotic-like effects of mGlu _2_ receptor activation in the mouse. Sci. Adv. 10, eadg1679. 10.1126/sciadv.adg1679.

50. Werthmann, R.C., Tzouros, M., Lamerz, J., Augustin, A., Fritzius, T., Trovò, L., Stawarski, M., Raveh, A., Diener, C., Fischer, C., et al. (2021). Symmetric signal transduction and negative allosteric modulation of heterodimeric mGlu1/5 receptors. Neuropharmacology 190, 108426. 10.1016/j.neuropharm.2020.108426.

51. Scholler, P., Nevoltris, D., de Bundel, D., Bossi, S., Moreno-Delgado, D., Rovira, X., Møller, T.C., El Moustaine, D., Mathieu, M., Blanc, E., et al. (2017). Allosteric nanobodies uncover a role of hippocampal mGlu2 receptor homodimers in contextual fear consolidation. Nat Commun 8, 1967. 10.1038/s41467-017-01489-1.

52. Zhao, Y., Chen, S., Swensen, A.C., Qian, W.-J., and Gouaux, E. (2019). Architecture and subunit arrangement of native AMPA receptors elucidated by cryo-EM. Science 364, 355–362. 10.1126/science.aaw8250.

53. Yu, J., Rao, P., Clark, S., Mitra, J., Ha, T., and Gouaux, E. (2021). Hippocampal AMPA receptor assemblies and mechanism of allosteric inhibition. Nature 594, 448–453. 10.1038/s41586-021-03540-0.

54. Sun, C., Zhu, H., Clark, S., and Gouaux, E. (2023). Cryo-EM structures reveal native GABAA receptor assemblies and pharmacology. Nature 622, 195–201. 10.1038/s41586-023-06556-w.

55. Zhou, J., Noviello, C.M., Teng, J., Moore, H., Lega, B., and Hibbs, R.E. (2025). Resolving native GABAA receptor structures from the human brain. Nature 638, 562–568. 10.1038/s41586-024-08454-1.

56. Zhang, M., Feng, J., Xie, C., Song, N., Jin, C., Wang, J., Zhao, Q., Zhang, L., Wang, B., Sun, Y., et al. (2025). Assembly and architecture of endogenous NMDA receptors in adult cerebral cortex and hippocampus. Cell 188, 1198–1207.e13. 10.1016/j.cell.2025.01.004.

57. Xu, R., Jiang, Q., Xu, H., Zhang, L., Hu, X., Lu, Z., Deng, H., Xiong, H., Zhang, S., Chen, Z., et al. (2026). Conformational diversity and fully opening mechanism of native NMDA receptor. Nature 652, 1405–1414. 10.1038/s41586-026-10139-w.

58. Park, J., and Gouaux, E. (2026). Efficient and rapid isolation of native AMPA receptor complexes for cryo-EM. Protein Science 35, e70483. 10.1002/pro.70483.

59. Patil, S.T., Zhang, L., Martenyi, F., Lowe, S.L., Jackson, K.A., Andreev, B.V., Avedisova, A.S., Bardenstein, L.M., Gurovich, I.Y., Morozova, M.A., et al. (2007). Activation of mGlu2/3 receptors as a new approach to treat schizophrenia: a randomized Phase 2 clinical trial. Nat Med 13, 1102–1107. 10.1038/nm1632.

60. Levitz, J., Habrian, C., Bharill, S., Fu, Z., Vafabakhsh, R., and Isacoff, E.Y. (2016). Mechanism of Assembly and Cooperativity of Homomeric and Heteromeric Metabotropic Glutamate Receptors. Neuron 92, 143–159. 10.1016/j.neuron.2016.08.036.

61. Chiu, Y.-T., Deutch, A.Y., Wang, W., Schmitz, G.P., Huang, K.L., Kocak, D.D., Llorach, P., Bowyer, K., Liu, B., Sciaky, N., et al. (2023). A suite of engineered mice for interrogating psychedelic drug actions. Preprint at bioRxiv, 10.1101/2023.09.25.559347 https://doi.org/10.1101/2023.09.25.559347.

62. Wright, R.A., Johnson, B.G., Zhang, C., Salhoff, C., Kingston, A.E., Calligaro, D.O., Monn, J.A., Schoepp, D.D., and Marek, G.J. (2013). CNS distribution of metabotropic glutamate 2 and 3 receptors: Transgenic mice and [3H]LY459477 autoradiography. Neuropharmacology 66, 89–98. 10.1016/j.neuropharm.2012.01.019.

63. Fordyce, B.A., Chiu, Y.-T., Wright, N.J., Sakamoto, K., Lyons, S.P., Webb, T.S., Tilton, H.E., Walsh, J.J., Marek, G., Setola, V., et al. (2026). No evidence for direct physical interaction of 5-HT2A-mGluR2 receptors in vitro or in vivo. Preprint at bioRxiv, 10.64898/2026.06.28.734515 https://doi.org/10.64898/2026.06.28.734515.

64. Schoepp, D.D., Johnson, B.G., Wright, R.A., Salhoff, C.R., Mayne, N.G., Wu, S., Cockerman, S.L., Burnett, J.P., Belegaje, R., Bleakman, D., et al. (1997). LY354740 is a potent and highly selective group II metabotropic glutamate receptor agonist in cells expressing human glutamate receptors. Neuropharmacology 36, 1–11. 10.1016/s0028-3908(96)00160-8.

65. Stevens, T.A., Tomaleri, G.P., Hazu, M., Wei, S., Nguyen, V.N., DeKalb, C., Voorhees, R.M., and Pleiner, T. (2024). A nanobody-based strategy for rapid and scalable purification of human protein complexes. Nat Protoc 19, 127–158. 10.1038/s41596-023-00904-w.

66. Fridy, P.C., Li, Y., Keegan, S., Thompson, M.K., Nudelman, I., Scheid, J.F., Oeffinger, M., Nussenzweig, M.C., Fenyö, D., Chait, B.T., et al. (2014). A robust pipeline for rapid production of versatile nanobody repertoires. Nat Methods 11, 1253–1260. 10.1038/nmeth.3170.

67. Chen, X., Liu, H., Shim, A.H.R., Focia, P.J., and He, X. (2008). Structural basis for synaptic adhesion mediated by neuroligin-neurexin interactions. Nat Struct Mol Biol 15, 50–56. 10.1038/nsmb1350.

68. Gjørlund, M.D., Carlsen, E.M.M., Kønig, A.B., Dmytrieva, O., Petersen, A.V., Jacobsen, J., Berezin, V., Perrier, J.-F., and Owczarek, S. (2017). Soluble Ectodomain of Neuroligin 1 Decreases Synaptic Activity by Activating Metabotropic Glutamate Receptor 2. Front. Mol. Neurosci. 10. 10.3389/fnmol.2017.00116.

69. Doornbos, M.L.J., Pérez-Benito, L., Tresadern, G., Mulder-Krieger, T., Biesmans, I., Trabanco, A.A., Cid, J.M., Lavreysen, H., IJzerman, A.P., and Heitman, L.H. (2016). Molecular mechanism of positive allosteric modulation of the metabotropic glutamate receptor 2 by JNJ-46281222. British Journal of Pharmacology 173, 588–600. 10.1111/bph.13390.

70. El-Baba, T.J., Lutomski, C.A., Bennett, J.L., Lawrence, S.A.S., Burnap, S.A., Butroid, F.I., Ramsay, O.B., Radzevičius, T., Wu, D., Song, H., et al. (2026). Molecular dissection of protein complexes isolated from sections of human brain. Preprint at bioRxiv, 10.64898/2026.04.11.717886 https://doi.org/10.64898/2026.04.11.717886.

71. Kim, Y., Gumpper, R.H., Liu, Y., Kocak, D.D., Xiong, Y., Cao, C., Deng, Z., Krumm, B.E., Jain, M.K., Zhang, S., et al. (2024). Bitter taste receptor activation by cholesterol and an intracellular tastant. Nature 628, 664–671. 10.1038/s41586-024-07253-y.

72. Krumm, B.E., DiBerto, J.F., Olsen, R.H.J., Kang, H.J., Slocum, S.T., Zhang, S., Strachan, R.T., Huang, X.-P., Slosky, L.M., Pinkerton, A.B., et al. (2023). Neurotensin Receptor Allosterism Revealed in Complex with a Biased Allosteric Modulator. Biochemistry 62, 1233–1248. 10.1021/acs.biochem.3c00029.

73. Choi, J., Chen, J., Schreiber, S.L., and Clardy, J. (1996). Structure of the FKBP12-Rapamycin Complex Interacting with Binding Domain of Human FRAP. Science 273, 239–242. 10.1126/science.273.5272.239.

74. Kunishima, N., Shimada, Y., Tsuji, Y., Sato, T., Yamamoto, M., Kumasaka, T., Nakanishi, S., Jingami, H., and Morikawa, K. (2000). Structural basis of glutamate recognition by a dimeric metabotropic glutamate receptor. Nature 407, 971–977. 10.1038/35039564.

75. Tora, A.S., Rovira, X., Cao, A.-M., Cabayé, A., Olofsson, L., Malhaire, F., Scholler, P., Baik, H., Van Eeckhaut, A., Smolders, I., et al. (2018). Chloride ions stabilize the glutamate-induced active state of the metabotropic glutamate receptor 3. Neuropharmacology 140, 275–286. 10.1016/j.neuropharm.2018.08.011.

76. DiRaddo, J.O., Miller, E.J., Bowman-Dalley, C., Wroblewska, B., Javidnia, M., Grajkowska, E., Wolfe, B.B., Liotta, D.C., and Wroblewski, J.T. (2015). Chloride is an Agonist of Group II and III Metabotropic Glutamate Receptors. Molecular Pharmacology 88, 450–459. 10.1124/mol.114.096420.

77. Monn, J.A., Prieto, L., Taboada, L., Pedregal, C., Hao, J., Reinhard, M.R., Henry, S.S., Goldsmith, P.J., Beadle, C.D., Walton, L., et al. (2015). Synthesis and Pharmacological Characterization of C4-Disubstituted Analogs of 1S,2S,5R,6S-2-Aminobicyclo[3.1.0]hexane-2,6-dicarboxylate: Identification of a Potent, Selective Metabotropic Glutamate Receptor Agonist and Determination of Agonist-Bound Human mGlu2 and mGlu3 Amino Terminal Domain Structures. J. Med. Chem. 58, 1776–1794. 10.1021/jm501612y.

78. Monn, J.A., Prieto, L., Taboada, L., Hao, J., Reinhard, M.R., Henry, S.S., Beadle, C.D., Walton, L., Man, T., Rudyk, H., et al. (2015). Synthesis and Pharmacological Characterization of C4-(Thiotriazolyl)-substituted-2-aminobicyclo[3.1.0]hexane-2,6-dicarboxylates. Identification of (1R,2S,4R,5R,6R)-2-Amino-4-(1H-1,2,4-triazol-3-ylsulfanyl)bicyclo[3.1.0]hexane-2,6-dicarboxylic Acid (LY2812223), a Highly Potent, Functionally Selective mGlu2 Receptor Agonist. J. Med. Chem. 58, 7526–7548. 10.1021/acs.jmedchem.5b01124.

79. Monn, J.A., Henry, S.S., Massey, S.M., Clawson, D.K., Chen, Q., Diseroad, B.A., Bhardwaj, R.M., Atwell, S., Lu, F., Wang, J., et al. (2018). Synthesis and Pharmacological Characterization of C4β-Amide-Substituted 2-Aminobicyclo[3.1.0]hexane-2,6-dicarboxylates. Identification of (1S,2S,4S,5R,6S)-2-Amino-4-[(3-methoxybenzoyl)amino]bicyclo[3.1.0]hexane-2,6-dicarboxylic Acid (LY2794193), a Highly Potent and Selective mGlu3 Receptor Agonist. J. Med. Chem. 61, 2303–2328. 10.1021/acs.jmedchem.7b01481.

80. Olsen, R.H.J., DiBerto, J.F., English, J.G., Glaudin, A.M., Krumm, B.E., Slocum, S.T., Che, T., Gavin, A.C., McCorvy, J.D., Roth, B.L., et al. (2020). TRUPATH, an open-source biosensor platform for interrogating the GPCR transducerome. Nat Chem Biol 16, 841–849. 10.1038/s41589-020-0535-8.

81. Black, J.W., and Leff, P. (1997). Operational models of pharmacological agonism. Proceedings of the Royal Society of London. Series B. Biological Sciences 220, 141–162. 10.1098/rspb.1983.0093.

82. Taussig, R., Iñiguez-Lluhi, J.A., and Gilman, A.G. (1993). Inhibition of Adenylyl Cyclase by Giα. Science 261, 218–221. 10.1126/science.8327893.

83. Chen, J., and Iyengar, R. (1993). Inhibition of cloned adenylyl cyclases by mutant-activated Gi-alpha and specific suppression of type 2 adenylyl cyclase inhibition by phorbol ester treatment. Journal of Biological Chemistry 268, 12253–12256. 10.1016/S0021-9258(18)31381-4.

84. Taussig, R., Tang, W.J., Hepler, J.R., and Gilman, A.G. (1994). Distinct patterns of bidirectional regulation of mammalian adenylyl cyclases. Journal of Biological Chemistry 269, 6093–6100. 10.1016/S0021-9258(17)37574-9.

85. Masuho, I., Skamangas, N.K., Muntean, B.S., and Martemyanov, K.A. (2021). Diversity of the Gβγ complexes defines spatial and temporal bias of GPCR signaling. Cell Systems 12, 324–337.e5. 10.1016/j.cels.2021.02.001.

86. Kankanamge, D., Tennakoon, M., Karunarathne, A., and Gautam, N. (2022). G protein gamma subunit, a hidden master regulator of GPCR signaling. Journal of Biological Chemistry 298. 10.1016/j.jbc.2022.102618.

87. Normoyle, K.P., Lillis, K.P., Egawa, K., McNally, M.A., Paulchakrabarti, M., Coudhury, B.P., Lau, L., Shiu, F.H., and Staley, K.J. (2025). Displacement of extracellular chloride by immobile anionic constituents of the brain’s extracellular matrix. The Journal of Physiology 603, 353–378. 10.1113/JP285463.

88. Ripke, S., Neale, B.M., Corvin, A., Walters, J.T.R., Farh, K.-H., Holmans, P.A., Lee, P., Bulik-Sullivan, B., Collier, D.A., Huang, H., et al. (2014). Biological insights from 108 schizophrenia-associated genetic loci. Nature 511, 421–427. 10.1038/nature13595.

89. Differential Expression of Metabotropic Glutamate Receptors 2 and 3 in Schizophrenia: A Mechanism for Antipsychotic Drug Action? American Journal of Psychiatry.

90. Wang, M., Zuris, J.A., Meng, F., Rees, H., Sun, S., Deng, P., Han, Y., Gao, X., Pouli, D., Wu, Q., et al. (2016). Efficient delivery of genome-editing proteins using bioreducible lipid nanoparticles. Proceedings of the National Academy of Sciences 113, 2868–2873. 10.1073/pnas.1520244113.

91. Yamada, K., McCarty, D.M., Madden, V.J., and Walsh, C.E. (2003). Lentivirus Vector Purification Using Anion Exchange HPLC Leads to Improved Gene Transfer. BioTechniques 34, 1074–1080. 10.2144/03345dd04.

92. Ai, H., Henderson, J.N., Remington, S.J., and Campbell, R.E. (2006). Directed evolution of a monomeric, bright and photostable version of Clavularia cyan fluorescent protein: structural characterization and applications in fluorescence imaging. Biochemical Journal 400, 531–540. 10.1042/BJ20060874.

93. Götzke, H., Kilisch, M., Martínez-Carranza, M., Sograte-Idrissi, S., Rajavel, A., Schlichthaerle, T., Engels, N., Jungmann, R., Stenmark, P., Opazo, F., et al. (2019). The ALFA-tag is a highly versatile tool for nanobody-based bioscience applications. Nat Commun 10, 4403. 10.1038/s41467-019-12301-7.

94. Tyanova, S., and Cox, J. (2018). Perseus: A Bioinformatics Platform for Integrative Analysis of Proteomics Data in Cancer Research. In Cancer Systems Biology: Methods and Protocols, L. von Stechow, ed. (Springer), pp. 133–148. 10.1007/978-1-4939-7493-1_7.

95. Kong, A.T., Leprevost, F.V., Avtonomov, D.M., Mellacheruvu, D., and Nesvizhskii, A.I. (2017). MSFragger: ultrafast and comprehensive peptide identification in mass spectrometry–based proteomics. Nat Methods 14, 513–520. 10.1038/nmeth.4256.

96. Yu, F., Haynes, S.E., and Nesvizhskii, A.I. (2021). IonQuant Enables Accurate and Sensitive Label-Free Quantification With FDR-Controlled Match-Between-Runs. Molecular & Cellular Proteomics 20. 10.1016/j.mcpro.2021.100077.

97. Pino, L.K., Searle, B.C., Bollinger, J.G., Nunn, B., MacLean, B., and MacCoss, M.J. (2020). The Skyline ecosystem: Informatics for quantitative mass spectrometry proteomics. Mass Spectrometry Reviews 39, 229–244. 10.1002/mas.21540.

98. Schorb, M., Haberbosch, I., Hagen, W.J.H., Schwab, Y., and Mastronarde, D.N. (2019). Software tools for automated transmission electron microscopy. Nat Methods 16, 471–477. 10.1038/s41592-019-0396-9.

99. Peck, J.V., Fay, J.F., and Strauss, J.D. (2022). High-speed high-resolution data collection on a 200 keV cryo-TEM. IUCrJ 9, 243–252. 10.1107/S2052252522000069.

100. Zheng, S.Q., Palovcak, E., Armache, J.-P., Verba, K.A., Cheng, Y., and Agard, D.A. (2017). MotionCor2: anisotropic correction of beam-induced motion for improved cryo-electron microscopy. Nat Methods 14, 331–332. 10.1038/nmeth.4193.

101. Kimanius, D., Dong, L., Sharov, G., Nakane, T., and Scheres, S.H.W. (2021). New tools for automated cryo-EM single-particle analysis in RELION-4.0. Biochemical Journal 478, 4169–4185. 10.1042/BCJ20210708.

102. Punjani, A., Rubinstein, J.L., Fleet, D.J., and Brubaker, M.A. (2017). cryoSPARC: algorithms for rapid unsupervised cryo-EM structure determination. Nat Methods 14, 290–296. 10.1038/nmeth.4169.

103. Rohou, A., and Grigorieff, N. (2015). CTFFIND4: Fast and accurate defocus estimation from electron micrographs. Journal of Structural Biology 192, 216–221. 10.1016/j.jsb.2015.08.008.

104. Bepler, T., Morin, A., Rapp, M., Brasch, J., Shapiro, L., Noble, A.J., and Berger, B. (2019). Positive-unlabeled convolutional neural networks for particle picking in cryo-electron micrographs. Nat Methods 16, 1153–1160. 10.1038/s41592-019-0575-8.

105. Kimanius, D., Jamali, K., Wilkinson, M.E., Lövestam, S., Velazhahan, V., Nakane, T., and Scheres, S.H.W. (2024). Data-driven regularization lowers the size barrier of cryo-EM structure determination. Nat Methods 21, 1216–1221. 10.1038/s41592-024-02304-8.

106. Asarnow, D., Palovcak, E., and Cheng, Y. (2019). UCSF pyem v0.5. Version v0.5 (Zenodo). 10.5281/zenodo.3576630 https://doi.org/10.5281/zenodo.3576630.

107. Wright, N.J., Zhang, F., Suo, Y., Kong, L., Yin, Y., Fedor, J.G., Sharma, K., Borgnia, M.J., Im, W., and Lee, S.-Y. (2024). Antiviral drug recognition and elevator-type transport motions of CNT3. Nat Chem Biol. 10.1038/s41589-024-01559-8.

108. Waterhouse, A., Bertoni, M., Bienert, S., Studer, G., Tauriello, G., Gumienny, R., Heer, F.T., de Beer, T.A.P., Rempfer, C., Bordoli, L., et al. (2018). SWISS-MODEL: homology modelling of protein structures and complexes. Nucleic Acids Res 46, W296–W303. 10.1093/nar/gky427.

109. Emsley, P., Lohkamp, B., Scott, W.G., and Cowtan, K. (2010). Features and development of Coot. Acta Cryst D 66, 486–501. 10.1107/S0907444910007493.

110. Williams, C.J., Headd, J.J., Moriarty, N.W., Prisant, M.G., Videau, L.L., Deis, L.N., Verma, V., Keedy, D.A., Hintze, B.J., Chen, V.B., et al. (2018). MolProbity: More and better reference data for improved all-atom structure validation. Protein Science 27, 293–315. 10.1002/pro.3330.

111. Adams, P.D., Afonine, P.V., Bunkóczi, G., Chen, V.B., Davis, I.W., Echols, N., Headd, J.J., Hung, L.-W., Kapral, G.J., Grosse-Kunstleve, R.W., et al. (2010). PHENIX: a comprehensive Python-based system for macromolecular structure solution. Acta Cryst D 66, 213–221. 10.1107/S0907444909052925.

112. Meng, E.C., Goddard, T.D., Pettersen, E.F., Couch, G.S., Pearson, Z.J., Morris, J.H., and Ferrin, T.E. (2023). UCSF ChimeraX: Tools for structure building and analysis. Protein Science 32, e4792. 10.1002/pro.4792.

113. DiRaddo, J.O., Miller, E.J., Hathaway, H.A., Grajkowska, E., Wroblewska, B., Wolfe, B.B., Liotta, D.C., and Wroblewski, J.T. (2014). A Real-Time Method for Measuring cAMP Production Modulated by Gαi/o-Coupled Metabotropic Glutamate Receptors. The Journal of Pharmacology and Experimental Therapeutics 349, 373–382. 10.1124/jpet.113.211532.

114. Huang, X.-P., Karpiak, J., Kroeze, W.K., Zhu, H., Chen, X., Moy, S.S., Saddoris, K.A., Nikolova, V.D., Farrell, M.S., Wang, S., et al. (2015). Allosteric ligands for the pharmacologically dark receptors GPR68 and GPR65. Nature 527, 477–483. 10.1038/nature15699.

115. Huang, X.-P., Kenakin, T.P., Gu, S., Shoichet, B.K., and Roth, B.L. (2020). Differential Roles of Extracellular Histidine Residues of GPR68 for Proton-Sensing and Allosteric Modulation by Divalent Metal Ions. Biochemistry 59, 3594–3614. 10.1021/acs.biochem.0c00576.

