## Supplemental for "Cryo-EM structures of brain-derived G protein-coupled receptors"

**Supplemental Figures**

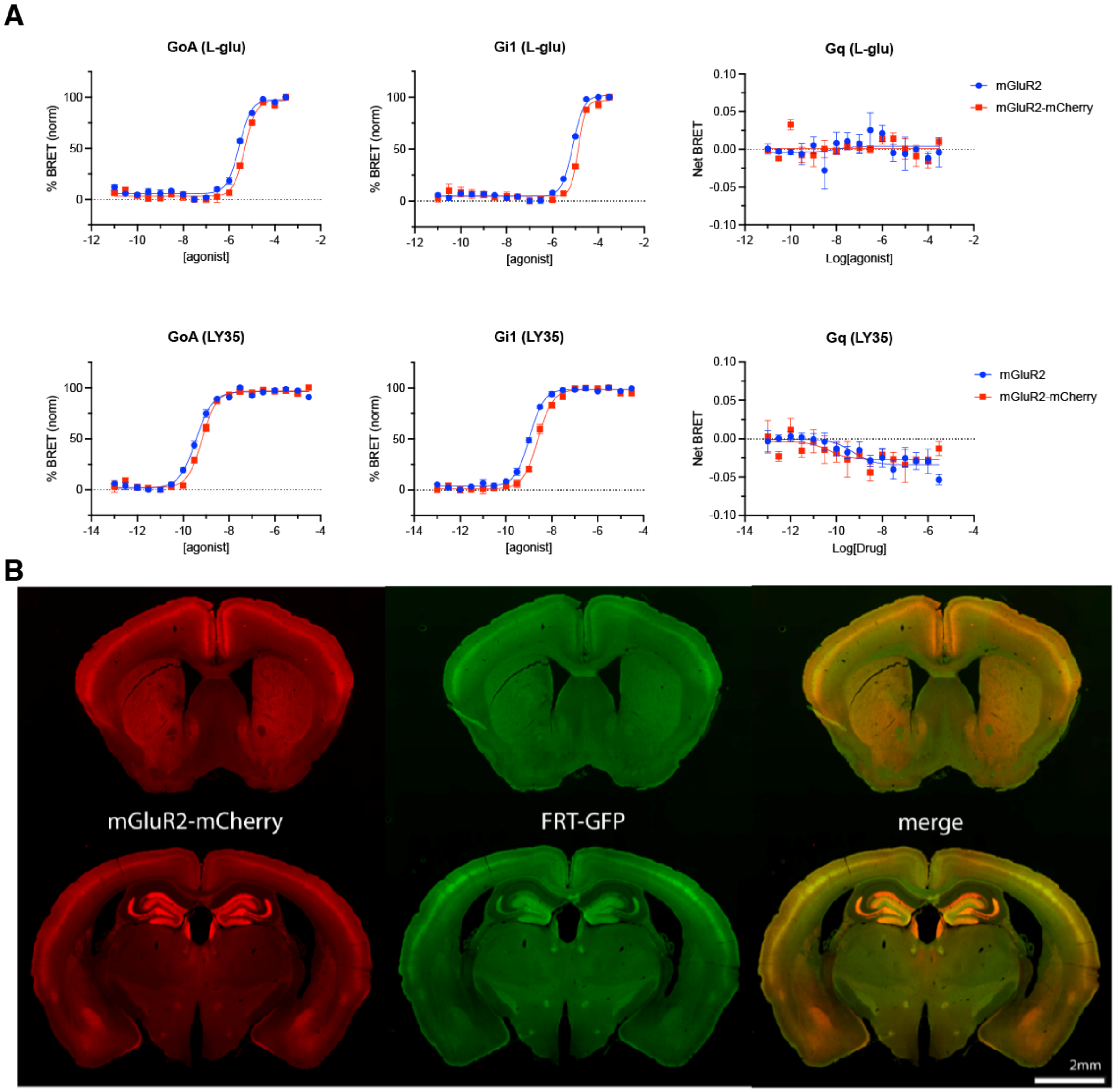


**Figure S1 | Validation of mGluR2-mCherry and *Grm2*$^{mCherry\text{-}FlpO}$ transgenic line**

**A,** Dose-response curves from TRUPATH for the WT unmodified and mCherry tagged mGluR2 (n=3 independent experiments; data shown as mean and s.e.m.) **B,** *Grm2* $^{+/mCherry\text{-}FlpO}$ x RCE (R26R CAG-boosted EGFP):FRT mice were used to validate FlpO functionality. Immunofluorescence was performed on brain sections an anti-RFP antibody (1:1000) to enhance the mGluR2–mCherry signal. GFP expression, resulting from FlpO–FRT–mediated recombination was detected and imaged using an Olympus slide scanner with a 10× objective. The experiments in this Figure were conducted with 3 mice with similar results.

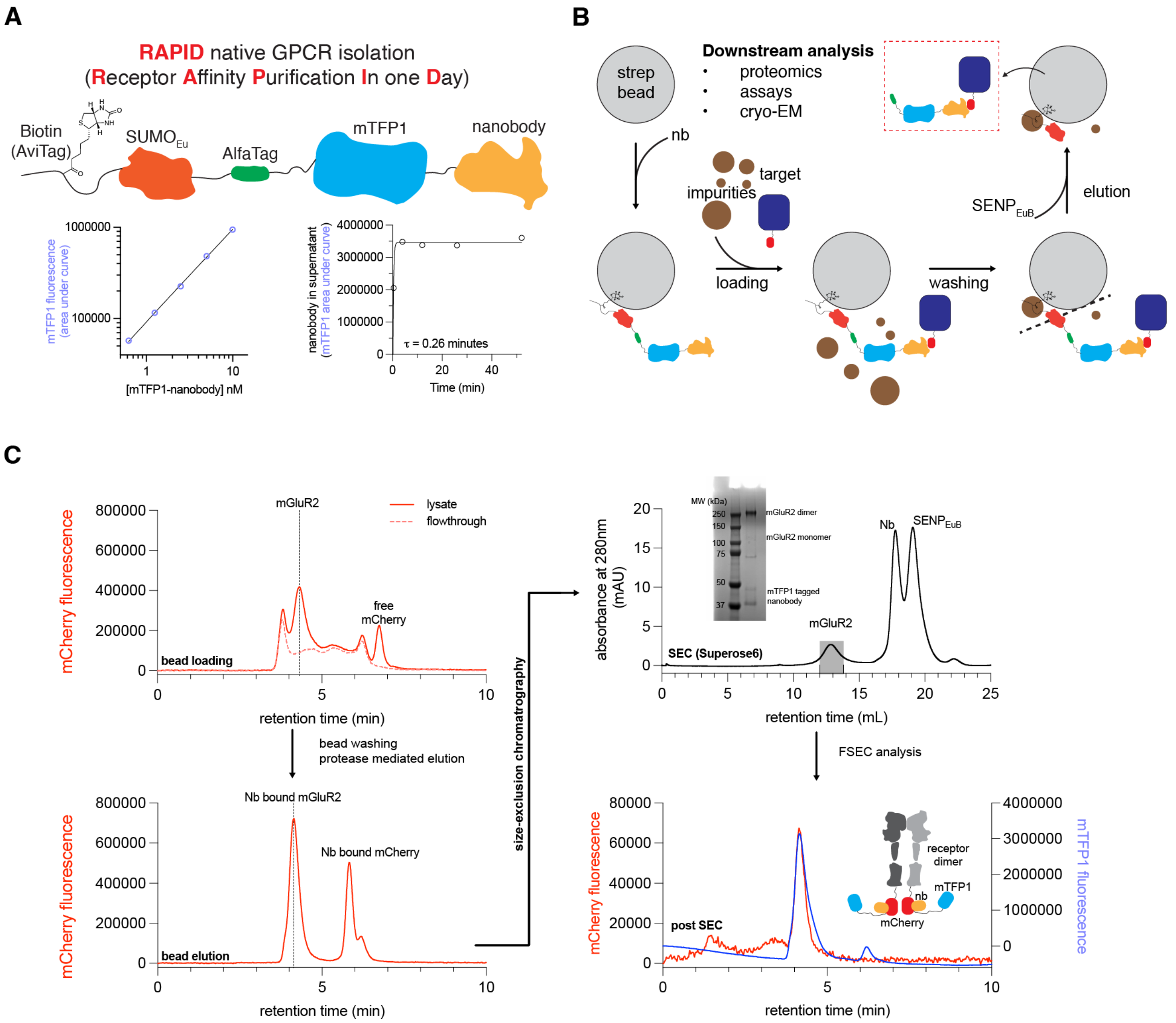


**Figure S2 | Nanobody-based receptor isolation optimization and validation**

**A,** Overview of the RAPID construct (top). Typical detection limits and dynamic range with fluorescence HPLC shown at bottom left. Timescale of SENP$_{EuB}$ protease mediated liberation of nanobody from streptavidin magnetic beads at bottom right. **B,** Cartoon schematic of RAPID mediated target isolation from cell lysates. **C,** Fluorescence HPLC analysis pipeline of RAPID isolation of recombinantly expressed (lentivirus) mGluR2-mCherry from HEK293T cells. Bead loading is evident by target peak decrease in the flowthrough (mCherry signal; top left); target monodispersity in bead elution (mCherry signal; bottom left) and SEC profile (A280 signal; top right) indicates biochemically stable receptor. SDS-PAGE analysis confirms target purity and identity (top right). Target identity was also confirmed by fluorescence HPLC of the SEC fractions, utilizing both mCherry and mTFP1 signal (bottom right).

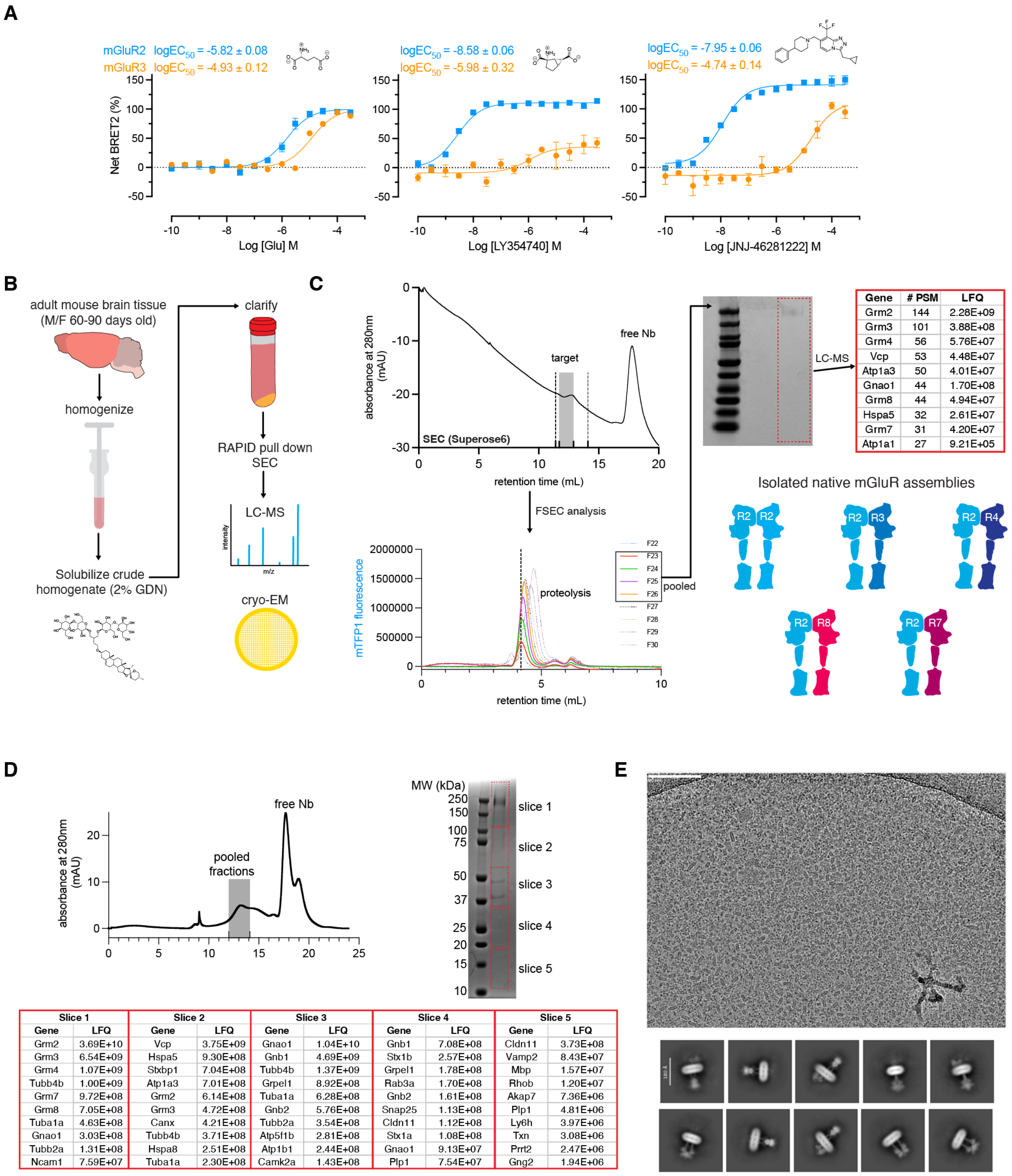


| Gene | # PSM | LFQ |
|---|---|---|
| Grm2 | 144 | 2.28E+09 |
| Grm3 | 101 | 3.88E+08 |
| Grm4 | 56 | 5.76E+07 |
| Vcp | 53 | 4.48E+07 |
| Atp1a3 | 50 | 4.01E+07 |
| Gnao1 | 44 | 1.70E+08 |
| Grm8 | 44 | 4.94E+07 |
| Hspa5 | 32 | 2.61E+07 |
| Grm7 | 31 | 4.20E+07 |
| Atp1a1 | 27 | 9.21E+05 |

| Slice 1 | | Slice 2 | | Slice 3 | | Slice 4 | | Slice 5 | |
|---|---|---|---|---|---|---|---|---|---|
| Gene | LFQ | Gene | LFQ | Gene | LFQ | Gene | LFQ | Gene | LFQ |
| Grm2 | 3.69E+10 | Vcp | 3.75E+09 | Gnao1 | 1.04E+10 | Gnb1 | 7.08E+08 | Cldn11 | 3.73E+08 |
| Grm3 | 6.54E+09 | Hspa5 | 9.30E+08 | Gnb1 | 4.69E+09 | Stx1b | 2.57E+08 | Vamp2 | 8.43E+07 |
| Grm4 | 1.07E+09 | Stxbp1 | 7.04E+08 | Tubb4b | 1.37E+09 | Grpel1 | 1.78E+08 | Mbp | 1.57E+07 |
| Tubb4b | 1.00E+09 | Atp1a3 | 7.01E+08 | Grpel1 | 8.92E+08 | Rab3a | 1.70E+08 | Rhob | 1.20E+07 |
| Grm7 | 9.72E+08 | Grm2 | 6.14E+08 | Tuba1a | 6.28E+08 | Gnb2 | 1.61E+08 | Akap7 | 7.36E+06 |
| Grm8 | 7.05E+08 | Grm3 | 4.72E+08 | Gnb2 | 5.76E+08 | Snap25 | 1.13E+08 | Plp1 | 4.81E+06 |
| Tuba1a | 4.63E+08 | Canx | 4.21E+08 | Tubb2a | 3.54E+08 | Cldn11 | 1.12E+08 | Ly6h | 3.97E+06 |
| Gnao1 | 3.03E+08 | Tubb4b | 3.71E+08 | Atp5f1b | 2.81E+08 | Stx1a | 1.08E+08 | Txn | 3.08E+06 |
| Tubb2a | 1.31E+08 | Hspa8 | 2.51E+08 | Atp1b1 | 2.44E+08 | Gnao1 | 9.13E+07 | Prrt2 | 2.47E+06 |
| Ncam1 | 7.59E+07 | Tuba1a | 2.30E+08 | Camk2a | 1.43E+08 | Plp1 | 7.54E+07 | Gng2 | 1.94E+06 |

**Figure S3 | RAPID isolation of mGluR2 containing endogenous complexes from brain tissue**

**A,** Concentration-response of L-glutamate, LY35, or JNJ at the murine receptors as assessed by $G_{oA}$ BRET2. Assays were performed in chloride-containing buffer for mGluR2 and in chloride-free buffer for mGluR3 (see Methods; data shown as mean and s.e.m. from *n*=3 experiments). **B,** Tissue work-up pipeline chart for isolation of mGluR2 complexes from mouse whole brains. **C,** Medium-scale mGluR2 complex RAPID isolation and subsequent size-exclusion chromatography (SEC) from whole brain tissue (10 brains). The dashed lines in the top left panel denote range of SEC fractions taken to FSEC analysis (F22-30), while the dark shaded box denotes SEC pooled fractions for SDS-PAGE and subsequent proteomics analysis (F23-26). Top 10 hits as ranked by peptide spectrum matches (PSMs) are shown in the top right panel, with possible homo- and heterodimeric assemblies shown in the bottom right panel. **D**, Size-exclusion chromatography profile of GDN-purified endogenous mGluR2 assemblies (top left), SDS-PAGE analysis of pooled peak fractions used for cryo-EM vitrification (top right), and proteomics analysis of selected band slices from the gel (bottom). **E**, Representative micrograph from the endogenous mGluR2 cryo-EM dataset (top) and representative 2D classes of the receptor particle projections (bottom).

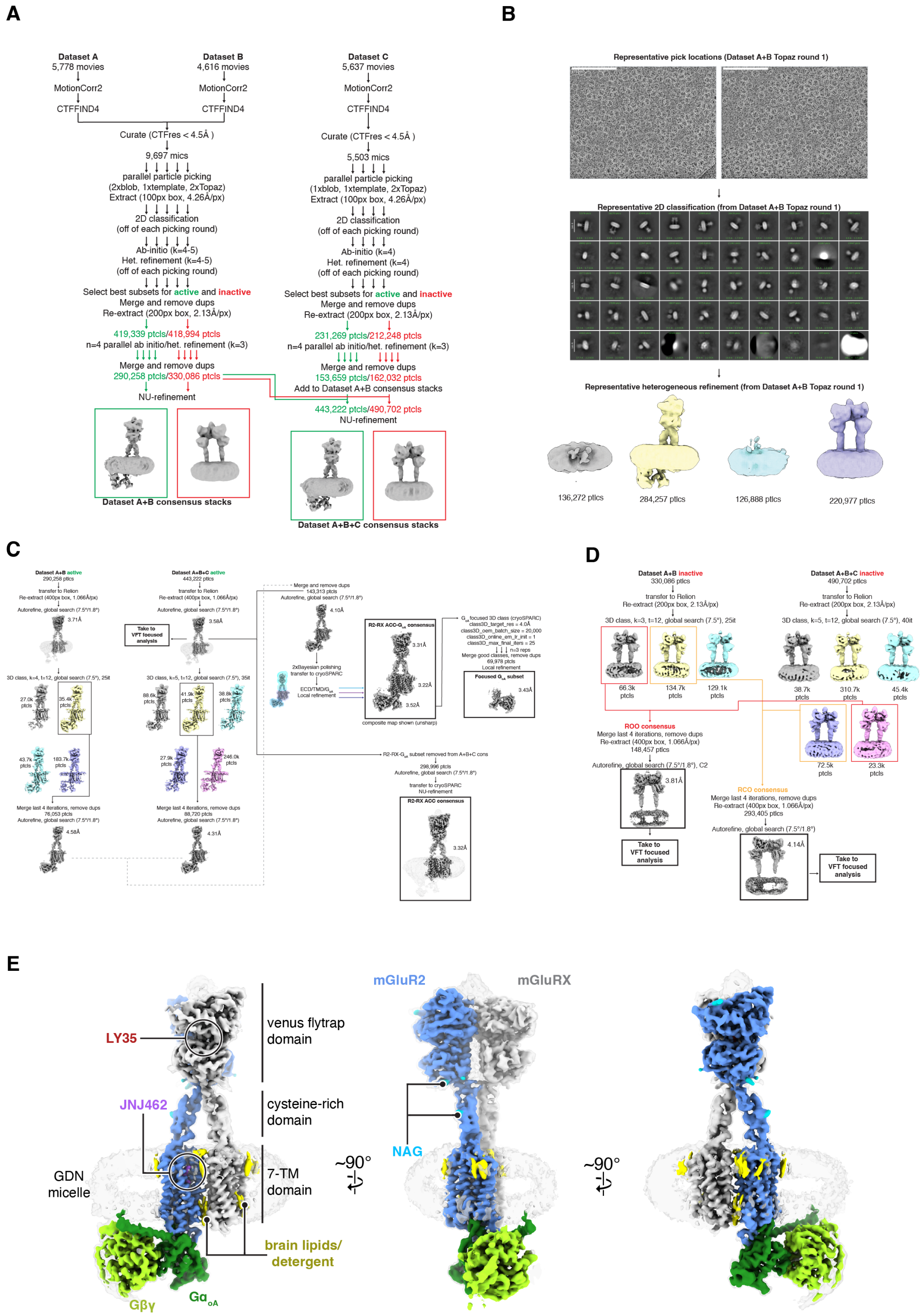


### Figure S4 | Initial image processing and 3D classification based on conformational state

**A,** Motion correction, CTF correction, particle picking, 2D classification and initial 3D classifications. **B,** Representative pick locations, 2D classes, and 3D classes from one of the replicate picking/classification runs. **C,** Processing pipeline for the active state conformational states, based on G-protein signal strength.

**D,** Processing pipeline for the inactive states, based on overall receptor conformation. **E,** Composite cryo-EM map (unsharpened, threshold=0.25) for the higher resolution consensus reconstruction of the ternary complex. The R2 assigned subunit is colored in blue, ambiguous subunit colored in grey, agonist colored red, ago-PAM colored purple, glycosylation (N-acetylglucosamine; NAG) in cyan, G-proteins colored green and detergents/brain lipids colored in yellow. A lower threshold (0.15) map rendering is overlayed in transparent grey to show the GDN micelle in relation to the protein elements of the complex.

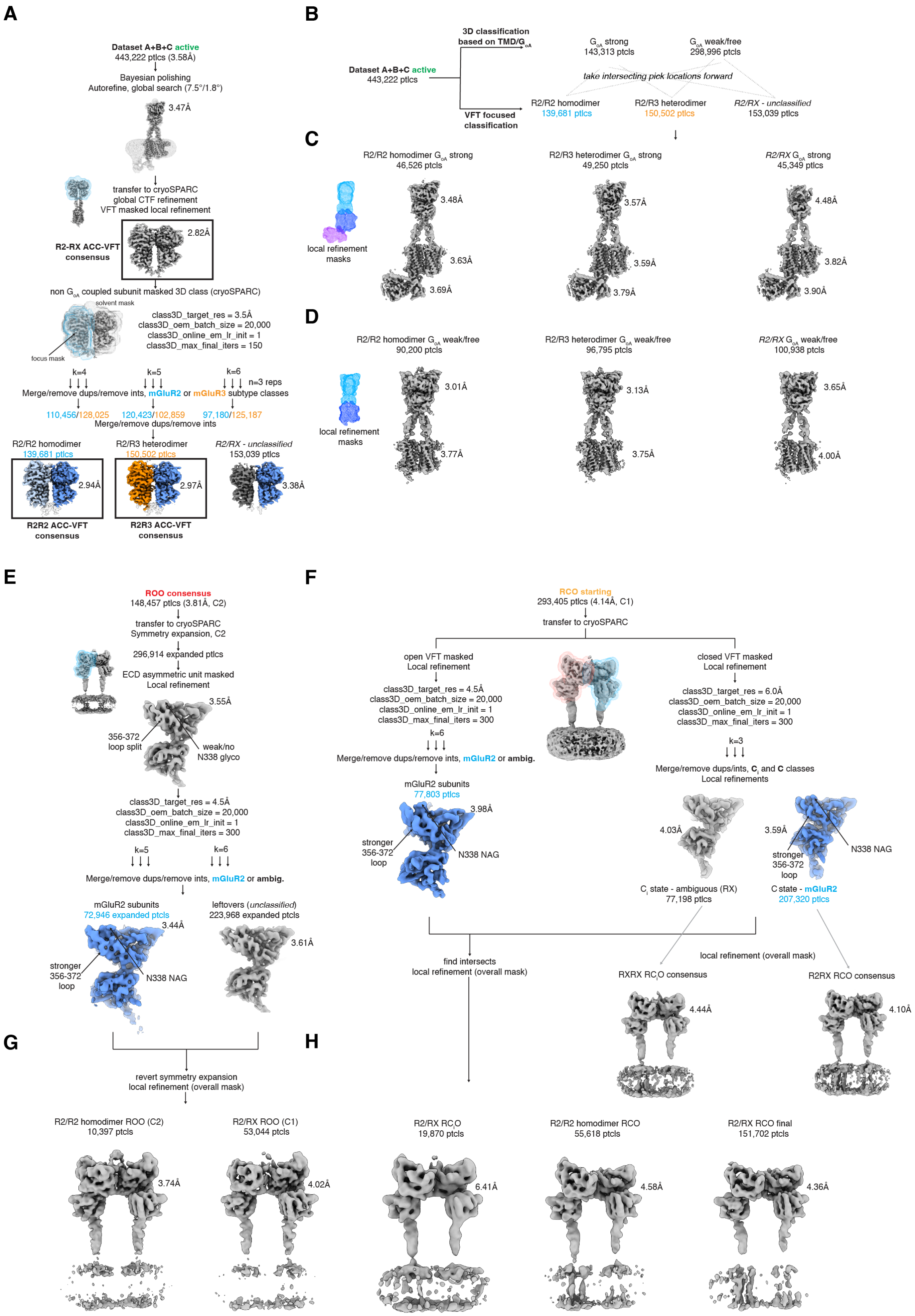
A
Dataset A+B+C active
443,222 ptlcs (3.58Å)
Bayesian polishing
Autorefine, global search (7.5°/1.8°)
3.47Å
transfer to cryoSPARC
global CTF refinement
VFT masked local refinement
R2-RX ACC-VFT consensus
2.82Å
non G$_{oA}$ coupled subunit masked 3D class (cryoSPARC)
solvent mask
focus mask
class3D_target_res = 3.5Å
class3D_oem_batch_size = 20,000
class3D_online_em_lr_init = 1
class3D_max_final_iters = 150
k=4
k=5
k=6
n=3 reps
Merge/remove dups/remove ints, mGluR2 or mGluR3 subtype classes
110,456/128,025
120,423/102,859
97,180/125,187
Merge/remove dups/remove ints
R2/R2 homodimer
139,681 ptlcs
R2/R3 heterodimer
150,502 ptlcs
R2/RX - unclassified
153,039 ptlcs
2.94Å
2.97Å
3.38Å
R2R2 ACC-VFT consensus
R2R3 ACC-VFT consensus
B
3D classification based on TMD/G$_{oA}$
Dataset A+B+C active
443,222 ptlcs
G$_{oA}$ strong
143,313 ptlcs
G$_{oA}$ weak/free
298,996 ptlcs
take intersecting pick locations forward
VFT focused classification
R2/R2 homodimer
139,681 ptlcs
R2/R3 heterodimer
150,502 ptlcs
R2/RX - unclassified
153,039 ptlcs
C
R2/R2 homodimer G$_{oA}$ strong
46,526 ptcls
R2/R3 heterodimer G$_{oA}$ strong
49,250 ptlcs
R2/RX G$_{oA}$ strong
45,349 ptlcs
local refinement masks
3.48Å
3.63Å
3.69Å
3.57Å
3.59Å
3.79Å
4.48Å
3.82Å
3.90Å
D
R2/R2 homodimer G$_{oA}$ weak/free
90,200 ptcls
R2/R3 heterodimer G$_{oA}$ weak/free
96,795 ptcls
R2/RX G$_{oA}$ weak/free
100,938 ptcls
local refinement masks
3.01Å
3.77Å
3.13Å
3.75Å
3.65Å
4.00Å
E
ROO consensus
148,457 ptcls (3.81Å, C2)
transfer to cryoSPARC
Symmetry expansion, C2
296,914 expanded ptlcs
ECD asymmetric unit masked
Local refinement
3.55Å
356-372 loop split
weak/no N338 glyco
class3D_target_res = 4.5Å
class3D_oem_batch_size = 20,000
class3D_online_em_lr_init = 1
class3D_max_final_iters = 300
k=5
k=6
Merge/remove dups/remove ints, mGluR2 or ambig.
mGluR2 subunits
72,946 expanded ptcls
leftovers (unclassified)
223,968 expanded ptcls
3.44Å
3.61Å
stronger 356-372 loop
N338 NAG
F
RCO starting
293,405 ptlcs (4.14Å, C1)
transfer to cryoSPARC
open VFT masked
Local refinement
class3D_target_res = 4.5Å
class3D_oem_batch_size = 20,000
class3D_online_em_lr_init = 1
class3D_max_final_iters = 300
k=6
Merge/remove dups/remove ints, mGluR2 or ambig.
mGluR2 subunits
77,803 ptlcs
3.98Å
stronger 356-372 loop
N338 NAG
closed VFT masked
Local refinement
class3D_target_res = 6.0Å
class3D_oem_batch_size = 20,000
class3D_online_em_lr_init = 1
class3D_max_final_iters = 300
k=3
Merge/remove dups/ints, C$_i$ and C classes
Local refinements
4.03Å
3.59Å
stronger 356-372 loop
N338 NAG
C$_i$ state - ambiguous (RX)
77,198 ptlcs
C state - mGluR2
207,320 ptlcs
find intersects
local refinement (overall mask)
local refinement (overall mask)
RXRX RC$_i$O consensus
R2RX RCO consensus
4.44Å
4.10Å
G
revert symmetry expansion
local refinement (overall mask)
R2/R2 homodimer ROO (C2)
10,397 ptlcs
R2/RX ROO (C1)
53,044 ptlcs
3.74Å
4.02Å
H
R2/RX RC$_i$O
19,870 ptlcs
R2/R2 homodimer RCO
55,618 ptlcs
R2/RX RCO final
151,702 ptlcs
6.41Å
4.58Å
4.36Å

**Figure S5 | VFT focused classifications based on subtypes**

**A,** Focused refinements and classifications of the VFT dimer within the active state consensus stack, to separate receptor subtypes. **B**, Schematic for finding particle intersects for distinct assemblies in distinct receptor states. **C**, Final reconstructions for distinct assemblies in the ACC-G state (unsharpened composite maps from ECD, 7TM and G-protein local refinements). GS-FSC resolutions from the local maps are shown next to the domain. **D**, Final reconstructions for distinct assemblies in the ACC state (unsharpened composite maps from ECD and 7TM local refinements). GS-FSC resolutions from the local maps are shown next to the domain. **E,** VFT focused classification (symmetry expansion) within the ROO state to identify R2 subtype subunit particles. **F**, VFT focused classification of the open or closed protomers within RCO to identify R2 subtype subunit particles. A partially closed state ($C_i$) was identified from further classification of the closed protomer. **G**, Reversion of symmetry expansion to identify particle subsets and obtain reconstructions for R2R2 ROO homodimer and R2RX ROO dimer. **H**, Particle intersection to identify particle subsets and obtain reconstructions for R2RX R$C_i$O dimer, R2R2 RCO homodimer, and R2RX RCO dimer.

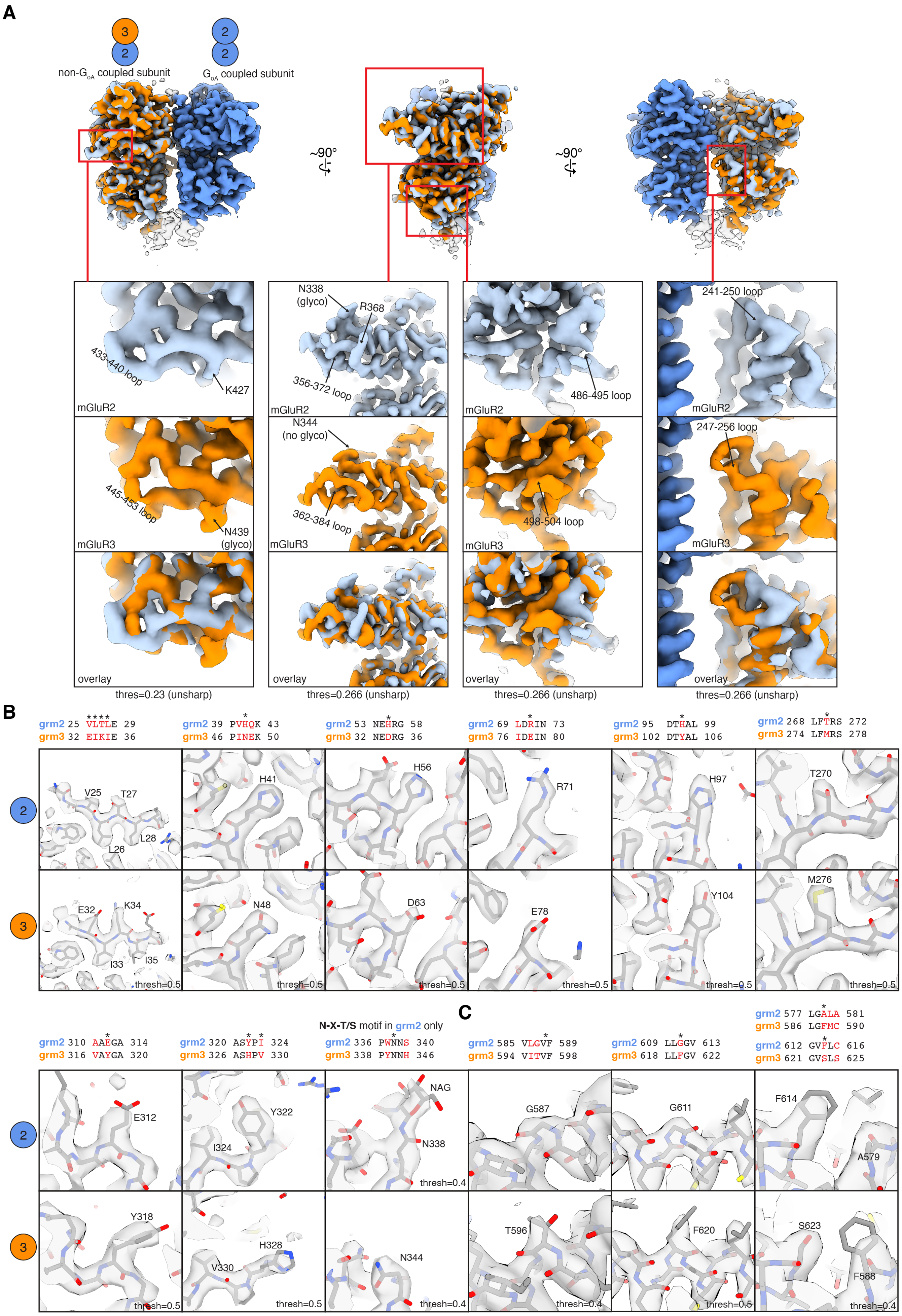

A
non-G$_{oA}$ coupled subunit
G$_{oA}$ coupled subunit
~90°
~90°
N338 (glyco)
R368
433-440 loop
K427
356-372 loop
486-495 loop
241-250 loop
mGluR2
N344 (no glyco)
247-256 loop
445-453 loop
N439 (glyco)
362-384 loop
498-504 loop
mGluR3
overlay
thres=0.23 (unsharp)
thres=0.266 (unsharp)
B
grm2 25 VLTLE 29
grm3 32 EIKIE 36
grm2 39 PVHQK 43
grm3 46 PINEK 50
grm2 53 NEHRG 58
grm3 32 NEDRG 36
grm2 69 LDRIN 73
grm3 76 IDEIN 80
grm2 95 DTHAL 99
grm3 102 DTYAL 106
grm2 268 LFTRS 272
grm3 274 LFMRS 278
V25
T27
L28
L26
H41
H56
R71
H97
T270
E32
K34
I33
I35
N48
D63
E78
Y104
M276
thresh=0.5
N-X-T/S motif in grm2 only
C
grm2 310 AAEGA 314
grm3 316 VAYGA 320
grm2 320 ASYPI 324
grm3 326 ASHPV 330
grm2 336 PWNNS 340
grm3 338 PYNNH 346
grm2 585 VLGVF 589
grm3 594 VITVF 598
grm2 609 LLGGV 613
grm3 618 LLFGV 622
grm2 577 LGALA 581
grm3 586 LGFMC 590
grm2 612 GVFLC 616
grm3 621 GVSLS 625
E312
Y322
I324
NAG
N338
G587
G611
F614
A579
Y318
H328
V330
N344
T596
F620
S623
F588
thresh=0.4

**Figure S6 | mGluR2 and mGluR3 subtype distinguishing features in the ACC states**

**A,** Overlay of the higher resolution R2R2 ACC VFT consensus and R2R3 ACC VFT consensus cryo-EM reconstructions, highlighting differences in divergent loop regions and subtype-specific glycosties (unsharpened maps shown). **B,** Subtype distinguishing positions within the VFT at chain B (non-$G_{oA}$ coupled subunit), highlighting differences in side-chain densities and subtype-specific glycosties (auto sharpened maps shown or the R2R2 ACC VFT and R2R3 ACC VFT consensus reconstructions). Multiple sequence alignment of subtypes R2 and R3 shown above each highlighted position. **C,** Subtype distinguishing positions within the 7TM domain of the final ACC-G reconstructions, highlighting differences in side-chain densities (chain B of the final R2R2 ACC-G and R2R3 ACC-G reconstructions; sharpened composite maps used). Signal from this domain was not utilized in 3D classifications to sort homodimers and heterodimers.

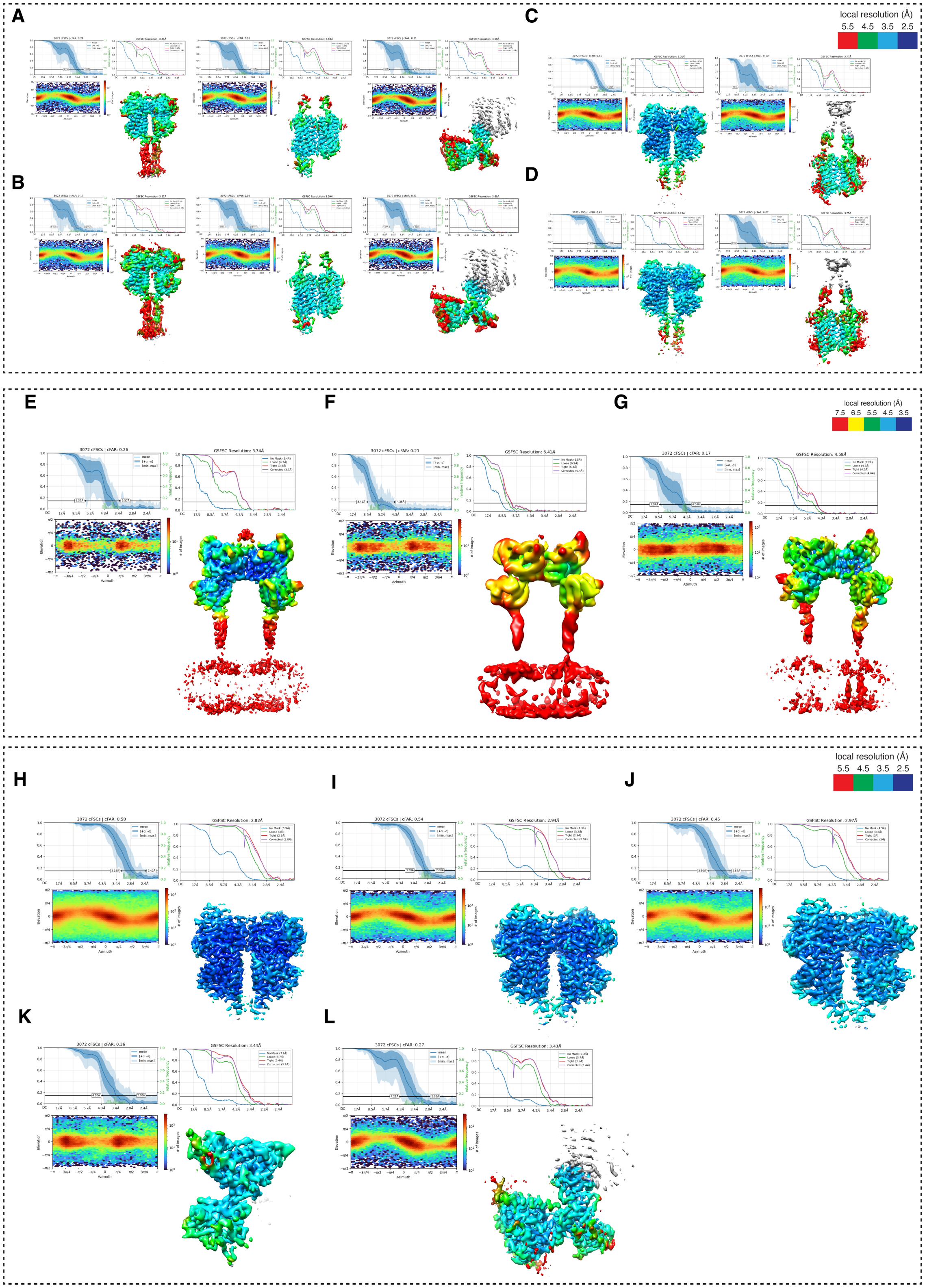


**Figure S7 | GS-FSC curves, particle assigned angular distributions, and local resolution for the endogenous receptor active state reconstructions**

Conical GS-FSC, masked GS-FSC, particle assigned orientation distributions and local resolution estimates for the ECD focused (left), 7TM focused (center), and G-protein focused (right) reconstructions for the distinct ACC-G states: **A,** R2R2-ACC-G; **B,** R2R3-ACC-G. Conical GS-FSC, masked GS-FSC, particle assigned orientation distributions and local resolution estimates for the ECD focused (left), 7TM focused (right), and G-protein focused (right) reconstructions for the distinct ACC states: **C,** R2R2-ACC; **D,** R2R3-ACC. Conical GS-FSC, masked GS-FSC, particle assigned orientation distributions and local resolution estimates for inactive states: **E**, R2R2-ROO; **F**, R2RX-RC$_i$O; **G**, R2R2-RCO. Conical GS-FSC, masked GS-FSC, particle assigned orientation distributions and local resolution estimates for the local reconstructions: **H,** R2RX ACC VFT consensus; **I,** R2R2 ACC VFT consensus; **J,** R2R3 ACC VFT consensus; **K,** R2 ROO VFT C2-expanded; **L,** $G_{oA}$ higher resolution subset. Volume regions outside of the refinement masks that have no local resolution value assigned are shown in grey.

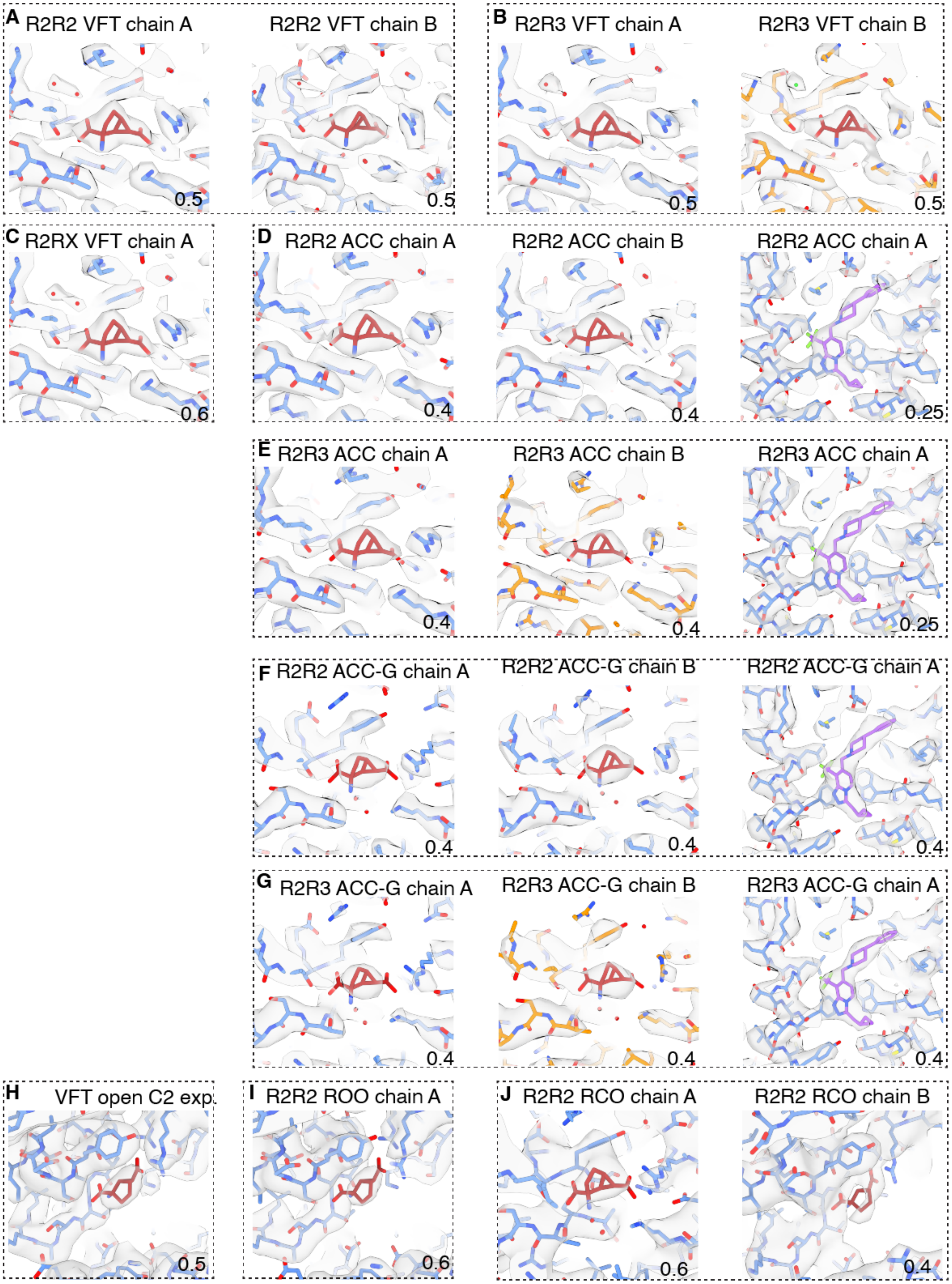
A R2R2 VFT chain A
R2R2 VFT chain B
0.5
0.5
B R2R3 VFT chain A
R2R3 VFT chain B
0.5
0.5
C R2RX VFT chain A
0.6
D R2R2 ACC chain A
R2R2 ACC chain B
R2R2 ACC chain A
0.4
0.4
0.25
E R2R3 ACC chain A
R2R3 ACC chain B
R2R3 ACC chain A
0.4
0.4
0.25
F R2R2 ACC-G chain A
R2R2 ACC-G chain B
R2R2 ACC-G chain A
0.4
0.4
0.4
G R2R3 ACC-G chain A
R2R3 ACC-G chain B
R2R3 ACC-G chain A
0.4
0.4
0.4
H VFT open C2 exp.
0.5
I R2R2 ROO chain A
0.6
J R2R2 RCO chain A
R2R2 RCO chain B
0.6
0.4

**Figure S8 | Ligand densities for the endogenous receptor reconstructions**

Cryo-EM map signal ligands in the endogenous receptor reconstructions: **A**, R2R2 ACC VFT consensus; **B**, R2R3 ACC VFT consensus; **C**, R2RX ACC VFT consensus; **D**, R2R2 ACC; **E**, R2R3 ACC; **F**, R2R2 ACC-G; **G**, R2R3 ACC-G; **H**, R2 ROO VFT (C2-expanded); **I**, R2R2 ROO; **J**, R2R2 RCO. LY35 colored in dark red, JNJ in purple, R2 protein in blue and R3 protein in orange. The title of the reconstruction and chain ID is shown per panel, and the map threshold used is shown at the bottom right of each panel.

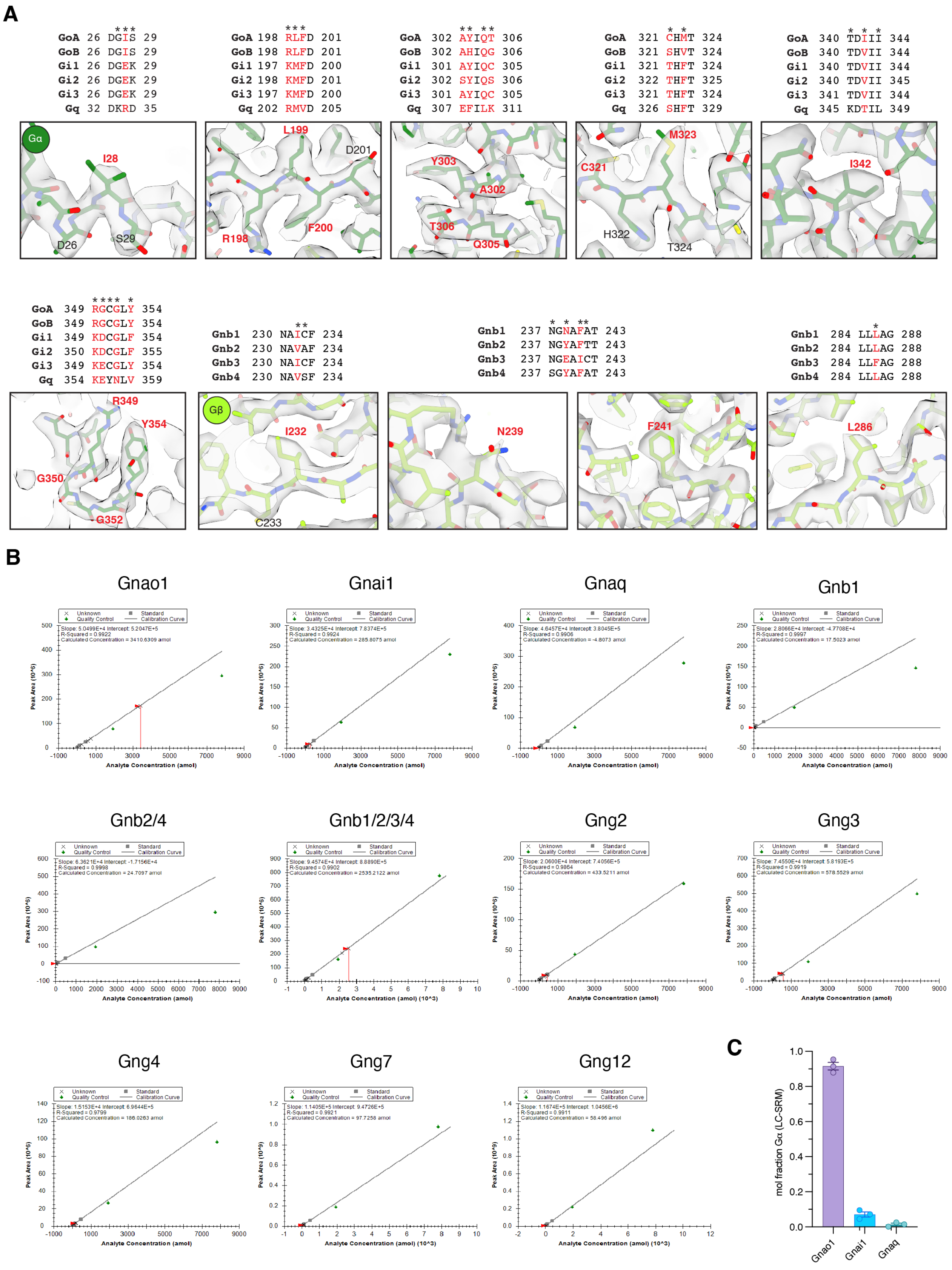
A
GoA 26 DGIS 29
GoB 26 DGIS 29
Gi1 26 DGEK 29
Gi2 26 DGEK 29
Gi3 26 DGEK 29
Gq 32 DKRD 35
GoA 198 RLFD 201
GoB 198 RLFD 201
Gi1 197 KMFD 200
Gi2 198 KMFD 201
Gi3 197 KMFD 200
Gq 202 RMVD 205
GoA 302 AYIQT 306
GoB 302 AHIQG 306
Gi1 301 AYIQC 305
Gi2 302 SYIQS 306
Gi3 301 AYIQC 305
Gq 307 EFILK 311
GoA 321 CHMT 324
GoB 321 SHVT 324
Gi1 321 THFT 324
Gi2 322 THFT 325
Gi3 321 THFT 324
Gq 326 SHFT 329
GoA 340 TDIII 344
GoB 340 TDVII 344
Gi1 340 TDVII 344
Gi2 340 TDVII 345
Gi3 341 TDVII 344
Gq 345 KDTIL 349
Gα
I28
D26
S29
L199
D201
R198
F200
Y303
A302
T306
Q305
C321
M323
H322
T324
I342
GoA 349 RGCGLY 354
GoB 349 RGCGLY 354
Gi1 349 KDCGLF 354
Gi2 350 KDCGLF 355
Gi3 349 KECGLY 354
Gq 354 KEYNLV 359
Gnb1 230 NAICF 234
Gnb2 230 NAVAF 234
Gnb3 230 NAICF 234
Gnb4 230 NAVSF 234
Gnb1 237 NGNAFAT 243
Gnb2 237 NGYAFTT 243
Gnb3 237 NGEAICT 243
Gnb4 237 SGYAFAT 243
Gnb1 284 LLLAG 288
Gnb2 284 LLLAG 288
Gnb3 284 LLFAG 288
Gnb4 284 LLLAG 288
R349
Y354
G350
G352
Gβ
I232
C233
N239
F241
L286
B
Gnao1
Gnai1
Gnaq
Gnb1
Gnb2/4
Gnb1/2/3/4
Gng2
Gng3
Gng4
Gng7
Gng12
Analyte Concentration (amol)
Analyte Concentration (amol) (10^3)
Peak Area (10^5)
Peak Area (10^9)
C
mol fraction Gα (LC-SRM)
Gnao1
Gnai1
Gnaq

**Figure S9 | Map-based subtype assignment for Gα and Gβ subunits**

**A**, Cryo-EM map and model for the high-resolution G-protein local subset, with multiple sequence alignment of relevant subtypes shown above each structural Region. Gα model depicted in dark green, Gβ model depicted in light green. Divergent positions of interest highlighted in red. **B**, Representative standard curves for LC-SRM for each proteolytic peptide standard, with unknown sample peak integrations plotted from a representative mGluR2 purification (drugs added). **C**, Summary of mol fractions of Gnao1, Gnai1 and Gnaq co-purified with mGluR2 from brain homogenates (drugs added) determined from LC-SRM (n=3 independent purifications).

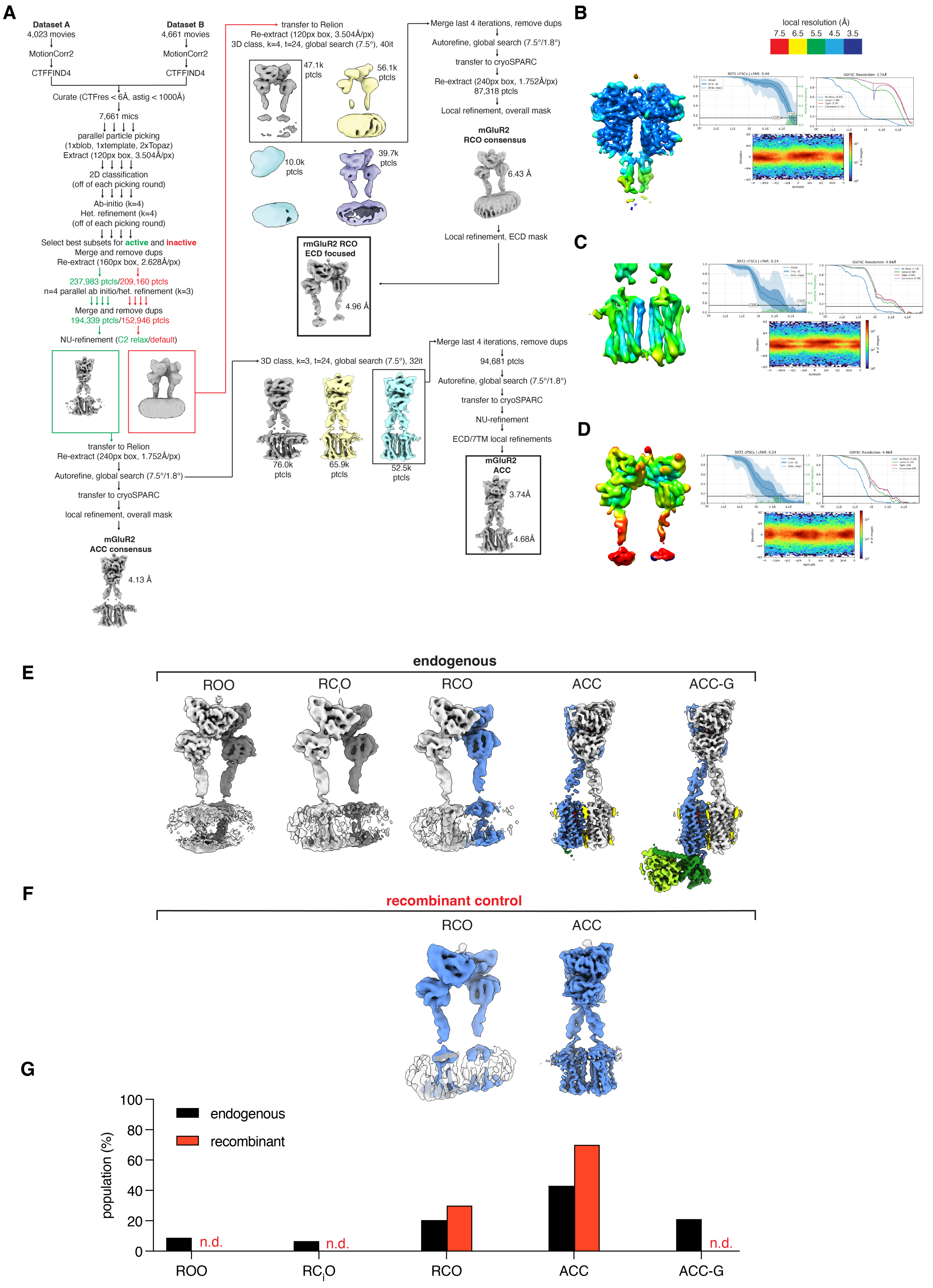
A
Dataset A
4,023 movies
MotionCorr2
CTFFIND4
Dataset B
4,661 movies
MotionCorr2
CTFFIND4
Curate (CTFres < 6Å, astig < 1000Å)
7,661 mics
parallel particle picking
(1xblob, 1xtemplate, 2xTopaz)
Extract (120px box, 3.504Å/px)
2D classification
(off of each picking round)
Ab-initio (k=4)
Het. refinement (k=4)
(off of each picking round)
Select best subsets for active and inactive
Merge and remove dups
Re-extract (160px box, 2.628Å/px)
237,983 ptcls/209,160 ptcls
n=4 parallel ab initio/het. refinement (k=3)
Merge and remove dups
194,339 ptcls/152,946 ptcls
NU-refinement (C2 relax/default)
transfer to Relion
Re-extract (240px box, 1.752Å/px)
Autorefine, global search (7.5°/1.8°)
transfer to cryoSPARC
local refinement, overall mask
mGluR2
ACC consensus
4.13 Å
transfer to Relion
Re-extract (120px box, 3.504Å/px)
3D class, k=4, t=24, global search (7.5°), 40it
47.1k
ptcls
56.1k
ptcls
10.0k
ptcls
39.7k
ptcls
rmGluR2 RCO
ECD focused
4.96 Å
Merge last 4 iterations, remove dups
Autorefine, global search (7.5°/1.8°)
transfer to cryoSPARC
Re-extract (240px box, 1.752Å/px)
87,318 ptcls
Local refinement, overall mask
mGluR2
RCO consensus
6.43 Å
Local refinement, ECD mask
3D class, k=3, t=24, global search (7.5°), 32it
76.0k
ptcls
65.9k
ptcls
52.5k
ptcls
Merge last 4 iterations, remove dups
94,681 ptcls
Autorefine, global search (7.5°/1.8°)
transfer to cryoSPARC
NU-refinement
ECD/7TM local refinements
mGluR2
ACC
3.74Å
4.68Å
B
local resolution (Å)
7.5 6.5 5.5 4.5 3.5
GSFSC Resolution: 3.74Å
C
GSFSC Resolution: 4.68Å
D
GSFSC Resolution: 4.96Å
E
endogenous
ROO
RC$_i$O
RCO
ACC
ACC-G
F
recombinant control
RCO
ACC
G
100
80
60
40
20
0
population (%)
endogenous
recombinant
n.d.
n.d.
n.d.
ROO
RC$_i$O
RCO
ACC
ACC-G

**Figure S10 | Image processing and validation of the recombinant control mGluR2 dataset**

**A,** Processing pipeline for the recombinant mGluR2 dataset. Conical GS-FSC, masked GS-FSC, particle assigned orientation distributions and local resolution estimates for: **B**, the ACC-ECD focused reconstruction; **C,** the ACC-7TM focused reconstruction; **D,** RCO-ECD focused reconstruction. **E,** Consensus reconstructions from the endogenous dataset compared with the consensus reconstructions from the recombinant mGluR2 control dataset (**F**). **G,** Particle population distributions across the conformational equilibria for either dataset. Only distinct particles were used for the distribution calculation (see Methods).

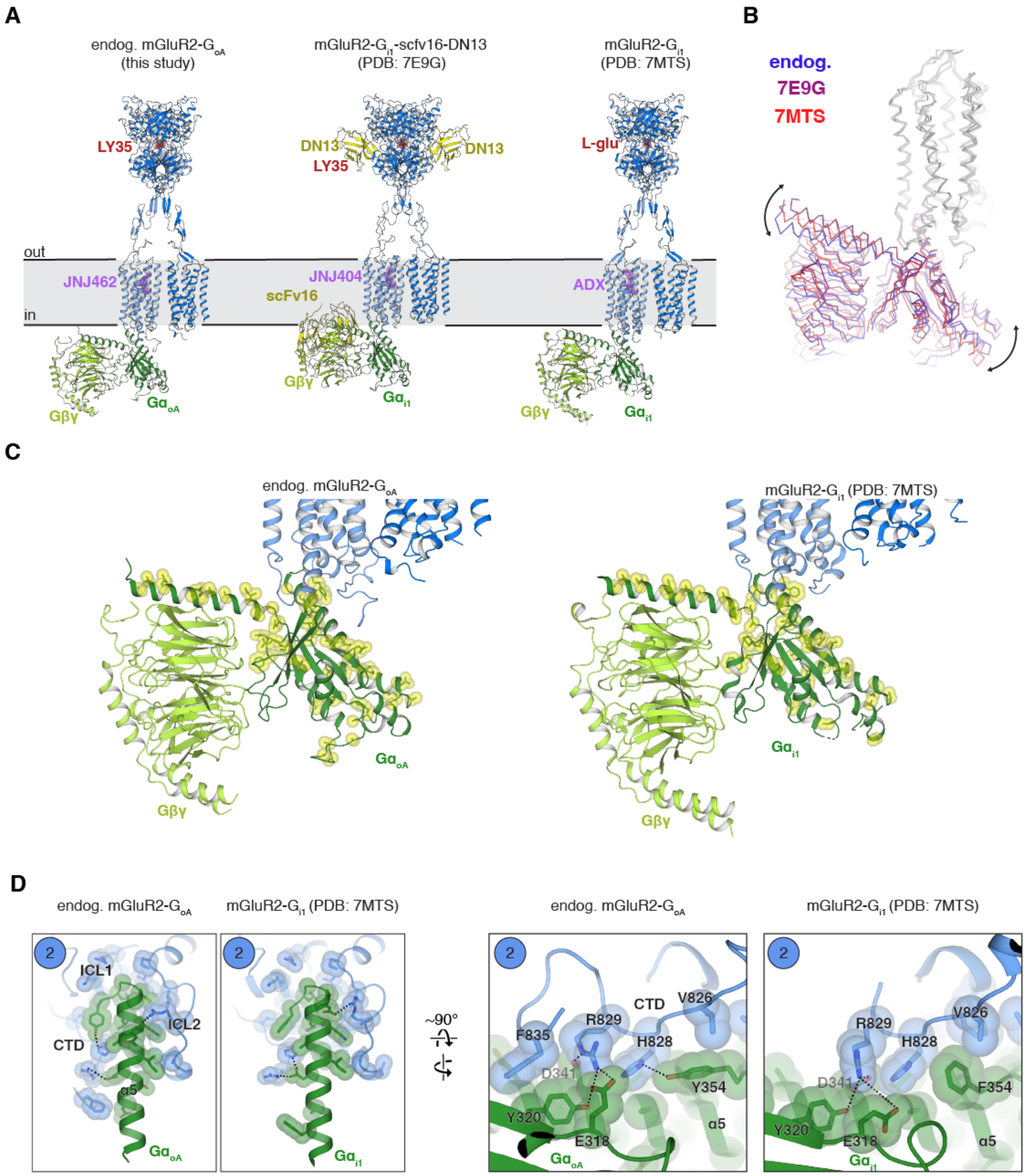


**Figure S11 | Endogenous mGluR2-G$_{oA}$ protein interactions compared with recombinant mGluR2-G$_{i1}$**

**A**, Comparison of the endogenous R2R2-ACC-G structure with representative published recombinant mGluR2-G$_i$ complex structures (PDB IDs 7E9G and 7MTS). **B,** Structural superposition of the endogenous R2R2-ACC-G structure with 7E9G and 7MTS. Alignment is targeted on the G-protein coupled 7TM**.** **C,** Divergent residues between Gnao1 and Gnai1 shown in yellow on the endogenous R2R2-ACC-G complex and on a representative recombinant mGluR2-G$_{i1}$ structure (7MTS). **D,** Comparison of the G-protein interaction interface in the endogenous or recombinant (7MTS) complex structures. The interaction interface is mainly comprised of receptor elements ICL1, ICL2 and C-terminus, in addition to the alpha-5 helix of the Gα subunit.

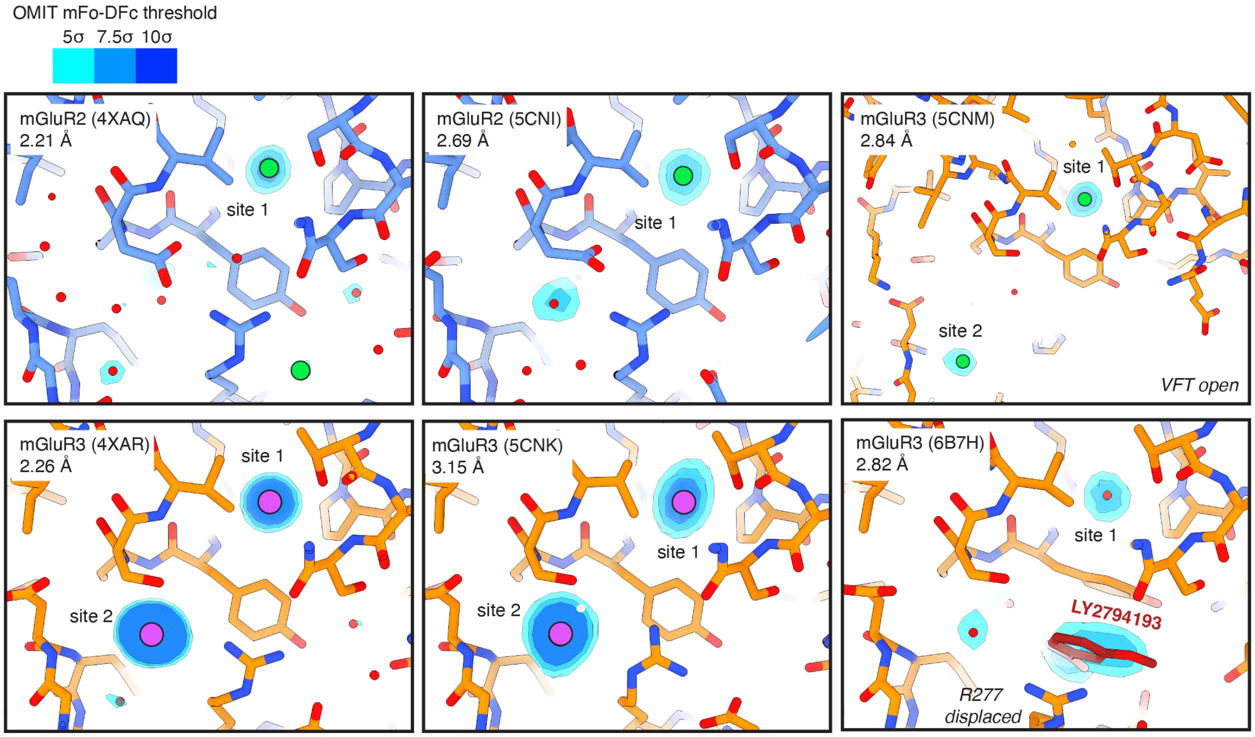


**Figure S12 | VFT ion modulatory sites within previous published X-ray structures**

OMIT mFo-DFc difference maps at the proposed chloride sites in previously published crystal structures of recombinantly expressed mGluR2 or mGluR3 VFTs (see Methods). Reported resolution and PDB ID shown in top left of each panel.

#### Supplemental Tables

##### **Table S1 |** Cryo-EM data collection statistics

| | Brain isolated mGluR2 complexes<br>+ 10 μM LY35<br>+10 μM JNJ462 | HEK293T rmGluR2 control<br>+ 1 μM LY35<br>+1 μM JNJ462 |
|---|---|---|
| **Data collection and processing** | | |
| Microscope | PNCC Krios #3<br>Krios | UNC-CH Cryo-EM core<br>Artica |
| Electron Gun | FEG | FEG |
| Detector | Gatan K3 | Gatan K3 |
| Magnification | 81,000 | 45,000 |
| Voltage (kV) | 300 | 200 |
| Electron exposure (e–/$Å^2$) | 50 | 54 |
| Energy filter slit width (eV) | 10 | n/a |
| Defocus range (μm) | -2.2 to -0.7 | -1.5 to -0.5 |
| Physical pixel size (Å) | 1.0655 (0.533 super-res) | 0.876 |
| Initial symmetry imposed | C1 | C1 |
| Number of collected movies | 16,031 | 8,684 |
| Initial clean receptor particle images (no.) | 875,254 | 281,657 |

**Table S2 |** Refinement and validation statistics for brain-isolated mGluR2 assemblies (global and composite reconstructions)

| | R2R2 ROO | R2RX $RC_iO$ | R2R2 RCO | R2R2 ACC | R2R3 ACC | R2R2 ACC-G | R2R3 ACC-G |
|---|---|---|---|---|---|---|---|
| **EMDB IDs*** | EMD-71943 | EMD-71944 | EMD-71945 | EMD-71941 (EMD-71931, EMD-71932) | EMD-71942 (EMD-71933, EMD-71934) | EMD-71940 (EMD-71925, EMD-71926, EMD-71927) | EMD-71939 (EMD-71928, EMD-71929, EMD-71930) |
| **PDB ID** | 9PWV | 9PWW | 9PWX | 9PWT | 9PWU | 9PWS | 9PWR |
| Symmetry imposed | C2 | C1 | C1 | C1 | C1 | C1 | C1 |
| Final particle images (no.) | 10,397 | 19,870 | 55,618 | 90,200 | 96,795 | 46,723 | 49,588 |
| Map resolution (Å)** | 3.74 | 6.41 | 4.58 | 3.01/3.77 | 3.13/3.75 | 3.48/3.63/3.69 | 3.57/3.59/3.79 |
| FSC threshold=0.143 | | | | | | | |
| **Refinement** | | | | | | | |
| Map sharpening *B* factor (Å$^2$)*** | -74.2 | 0 | -166.2 | -50/-60 | -50/-60 | -40/-60/-55 | -50/-60-/-60 |
| Model composition | | | | | | | |
| Non-hydrogen atoms | 7,120 | 5,022 | 6,858 | 11,403 | 11,350 | 15,672 | 15,712 |
| Protein residues | 1,018 | 1,018 | 1,015 | 1,516 | 1,520 | 2,154 | 2,143 |
| Ligands | 40F: 2, NAG: 8 | n/a | 40F: 2, NAG: 6 | 40F: 2, JNJ:1, NAG: 9, 9Z9: 1 | 40F: 2, JNJ:1, NAG: 9, 9Z9: 3 | 40F: 2, JNJ:1, NAG: 10, 9Z9: 9, LBN: 1 | 40F: 2, JNJ:1, NAG: 8, 9Z9: 6, LBN: 2 |
| *B* factors (Å$^2$) | | | | | | | |
| Protein | 221.70 | 1178.12 | 173.61 | 85.42 | 88.80 | 84.21 | 80.17 |
| Ligand | 255.70 | n/a | 191.61 | 112.57 | 113.15 | 80.65 | 77.02 |
| R.m.s. deviations | | | | | | | |
| Bond lengths (Å) | 0.005 | 0.005 | 0.006 | 0.007 | 0.006 | 0.005 | 0.006 |
| Bond angles (°) | 0.973 | 1.096 | 1.027 | 0.989 | 0.991 | 0.998 | 1.023 |
| Validation | | | | | | | |
| MolProbity score | 1.11 | 1.45 | 1.58 | 1.33 | 1.28 | 1.39 | 1.44 |
| Clashscore | 3.24 | 5.80 | 9.10 | 5.06 | 5.17 | 4.40 | 4.75 |
| Poor rotamers (%) | 0.00 | 0.00 | 0.40 | 0.37 | 0.38 | 0.95 | 0.21 |
| Ramachandran plot | | | | | | | |
| Favored (%) | 98.42 | 97.33 | 97.52 | 97.74 | 98.27 | 97.05 | 96.84 |
| Allowed (%) | 1.58 | 2.57 | 2.48 | 2.26 | 1.73 | 2.95 | 3.16 |
| Disallowed (%) | 0.00 | 0.10 | 0.00 | 0.00 | 0.00 | 0.00 | 0.00 |

*Focus map depositions in paratheses

**Map resolution for local refinements are provided (in the order: ECD/7TM/$G_{oA}$)

***Post-processing *B*-factors are provided for each local refinement used to generate the composite map (in the order: ECD/7TM/$G_{oA}$)

**Table S3 |** Refinement and validation statistics for brain-isolated mGluR2 assemblies (focused reconstructions)

| | R2RX ACC VFT consensus | R2R2 ACC VFT consensus | R2R3 ACC VFT consensus | $G_{oA}$ higher resolution subset | R2 ROO VFT (C2-expanded) |
|---|---|---|---|---|---|
| **EMDB ID** | EMD-71947 | EMD-71948 | EMD-71949 | EMD-71946 | EMD-71950 |
| **PDB ID** | 9PWZ | 9PX0 | 9PX1 | 9PWY | 9PX2 |
| Symmetry imposed | C1 | C1 | C1 | C1 | C2-expanded ASU |
| Final particle images (no.) | 443,222 | 139,681 | 150,502 | 69,978 | 72,946 (sym exp.) |
| Map resolution (Å) FSC threshold=0.143 | 2.82 | 2.94 | 2.97 | 3.43 | 3.44 |
| **Refinement** | | | | | |
| Map sharpening *B* factor (Å$^2$) | -94.8 | -79.8 | -83.5 | -98.0 | -60 |
| Model composition | | | | | |
| Non-hydrogen atoms | 3,556 | 7,027 | 7,043 | 4,264 | 3,411 |
| Protein residues | 453 | 906 | 909 | 602 | 453 |
| Ligands | 40F: 1, NAG: 5, CL: 1 | 40F: 2, NAG: 10, CL: 2 | 40F: 2, NAG: 9, CL: 3 | n/a | 40F: 1, NAG: 4 |
| *B* factors (Å$^2$) | | | | | |
| Protein | 72.43 | 65.71 | 76.00 | 85.29 | 124.48 |
| Ligand | 109.65 | 107.30 | 103.96 | n/a | 159.80 |
| R.m.s. deviations | | | | | |
| Bond lengths (Å) | 0.007 | 0.004 | 0.004 | 0.005 | 0.006 |
| Bond angles (°) | 1.021 | 0.941 | 0.911 | 0.917 | 1.024 |
| Validation | | | | | |
| MolProbity score | 1.46 | 1.29 | 1.26 | 1.39 | 1.18 |
| Clashscore | 6.64 | 3.22 | 4.91 | 5.78 | 3.96 |
| Poor rotamers (%) | 0.86 | 1.19 | 0.59 | 0.74 | 0.94 |
| Ramachandran plot | | | | | |
| Favored (%) | 97.55 | 97.77 | 98.22 | 97.64 | 98.00 |
| Allowed (%) | 2.45 | 2.23 | 1.78 | 2.36 | 2.00 |
| Disallowed (%) | 0.00 | 0.00 | 0.00 | 0.00 | 0.00 |

**Table S4 |** Refinement and validation statistics for recombinant mGluR2

| | recom. mGluR2 ACC (subset, composite) | recom. mGluR2 RCO (ECD local) |
|---|---|---|
| **EMDB ID*** | EMD-71951 (EMD-71935, EMD-71936) | EMD-71952 |
| **PDB ID** | 9PX3 | 9PX4 |
| Symmetry imposed | C1 | C1 |
| Final particle images (no.) | 94,681 | 87,318 |
| Map resolution (Å)** | 3.74/4.68 | 4.96 |
| FSC threshold=0.143 | | |
| **Refinement** | | |
| Map sharpening *B* factor (Å$^2$)*** | -80/-100 | -100 |
| Model composition | | |
| Non-hydrogen atoms | 9,931 | 5,106 |
| Protein residues | 1,526 | 1,018 |
| Ligands | 40F: 2, NAG: 8 | NAG: 6 |
| *B* factors (Å$^2$) | | |
| Protein | 113.00 | 269.11 |
| Ligand | 115.41 | 340.77 |
| R.m.s. deviations | | |
| Bond lengths (Å) | 0.008 | 0.005 |
| Bond angles (°) | 1.078 | 1.081 |
| Validation | | |
| MolProbity score | 1.27 | 0.93 |
| Clashscore | 4.05 | 1.19 |
| Poor rotamers (%)**** | 0.61 | 0.00 |
| Ramachandran plot | | |
| Favored (%) | 97.62 | 97.52 |
| Allowed (%) | 2.38 | 2.48 |
| Disallowed (%) | 0.00 | 0.00 |

*Focus map depositions in paratheses

**Map resolution for local refinements are provided (in the order: ECD/7TM)

***Post-processing *B*-factors are provided for each local refinement used to generate the composite map (in the order: ECD/7TM)

****side-chains truncated for the RCO structure owing to modest resolution of the reconstruction

#### **Table S5 |** Final distinct particle counts used for population analyses

**Recombinant mGluR2 control dataset**

| | ROO | RCiO | RCO | ACC | ACC-G | total | percent redundant |
|---|---|---|---|---|---|---|---|
| **particles** | n.d. | n.d. | 87318 | 194339 | n.d. | 281657 | ***5.6*** |
| **distinct particles** | n.d. | n.d. | 79793 | 186145 | n.d. | 265938 | |
| **percent (of distinct)** | n.d. | n.d. | 30.0 | 70.0 | n.d. | | |

**endogenous receptor dataset**

***consensus stacks***

| | ROO | RCiO | RCO | ACC | ACC-G | total | percent redundant |
|---|---|---|---|---|---|---|---|
| **particles** | 148457 | 77198 | 207320 | 298966 | 143313 | 875254 | ***26.4*** |
| **distinct particles** | 56604 | 42461 | 131713 | 277818 | 135956 | 644552 | |
| **percent (of distinct)** | 8.8 | 6.6 | 20.4 | 43.1 | 21.1 | | |

***distinct assemblies***

| | R2R2 - ROO | R2RX - ROO | R2RX - RCiO | R2R2 - RCO | R2RX -RCO | R2R2 - ACC | R2R3 - ACC | R2RX - ACC | R2R2 - ACC-G | R2R3 - ACC-G | R2RX - ACC-G | total | percent redundant |
|---|---|---|---|---|---|---|---|---|---|---|---|---|---|
| **particles** | 10397 | 53044 | 19870 | 55618 | 151702 | 90200 | 96795 | 100938 | 46723 | 49588 | 45350 | 720225 | ***13.1*** |
| **distinct particles** | 6679 | 29370 | 14621 | 44929 | 122679 | 85550 | 91827 | 94806 | 44847 | 47564 | 43117 | 625989 | |
| **percent (of distinct)** | 1.1 | 4.7 | 2.3 | 7.2 | 19.6 | 13.7 | 14.7 | 15.1 | 7.2 | 7.6 | 6.9 | | |

**Table S6 |** Proteolytic peptide standards for LC-SRM

| Peptide | Protein target | m/z | z | t start (min) | t stop (min) | Polarity |
|---|---|---|---|---|---|---|
| EDPLLTPVPASENPFR (light) | Gng2 | 891.4571 | 2 | 40 | 46 | Positive |
| IEASLC[+57.021464]R (light) | Gng3 | 424.7184 | 2 | 17 | 25 | Positive |
| EDPLIIPVPASENPFR (light) | Gng4 | 897.4753 | 2 | 44 | 52 | Positive |
| IEAGIER (light) | Gng7 | 394.2191 | 2 | 15 | 23 | Positive |
| LEASIER (light) | Gng12 | 409.2243 | 2 | 16 | 24 | Positive |
| YYLDSLDR (light) | Gnao1 | 522.7535 | 2 | 28 | 34 | Positive |
| IAQPNYIPTQQDVLR (light) | Gnai1 | 878.473 | 2 | 32 | 40 | Positive |
| EYQLSDSTK (light) | Gnaq | 535.7537 | 2 | 16 | 22 | Positive |
| QEAEQLK (light) | Gnb1 | 423.2218 | 2 | 5 | 16 | Positive |
| LLVSASQDGK (light) | Gnb1,2,3,4 | 509.2824 | 2 | 18 | 25 | Positive |
| QEAEQLR (light) | Gnb2,4 | 437.2249 | 2 | 9 | 17 | Positive |

#### **Table S7 |** Fitting parameters for chloride PAM assays

| | mGluR2 - L-Glu | mGluR2 - LY27 | mGluR2 - LY35 | mGluR2 - JNJ462 | mGluR3 - L-Glu | mGluR3 - LY27 | mGluR3 - LY35 | mGluR3 - JNJ462 |
|---|---|---|---|---|---|---|---|---|
| **LogKA** | = -5.250 | = -7.780 | = -7.903 | = -8.214 | = -5.300 | = -8.700 | = -6.364 | = -4.657 |
| **LogKB** | -0.5716 | = -0.9898 | -0.8986 | -1.396 | = -1.470 | = -1.470 | = -1.470 | = -1.470 |
| **Basal** | = 0.000 | = 0.000 | = 0.000 | = 0.000 | = 0.000 | = 0.000 | = 0.000 | = 0.000 |
| **Emax** | = 150.0 | = 150.0 | = 150.0 | = 150.0 | = 200.0 | = 200.0 | = 200.0 | = 200.0 |
| **Tao_A** | 1.44 | 1.382 | 1.556 | 1.841 | 0.9711 | 1.092 | 0.6975 | 0.9197 |
| **Alpha** | 2.108 | 1.926 | 2.565 | 0.3107 | 3.712 | 3.176 | 1.452 | 0.8107 |
| **Bta** | = 1.000 | = 1.000 | = 1.000 | = 1.000 | 1.124 | 1.112 | 1.888 | 2.064 |
| **B** | = 0.000 | = 0.000 | = 0.000 | = 0.000 | = 0.000 | = 0.000 | = 0.000 | = 0.000 |
| **Tao_B** | 1.849 | 1.197 | 1.165 | 0.7419 | 0.993 | 1.185 | 1.197 | 1.002 |
| **n** | 2.405 | 2.249 | 2.32 | 2.33 | 1.605 | 2.192 | 1.861 | 1.796 |
| **log alpha** | 0.3238 | 0.2847 | 0.4091 | -0.5076 | 0.5696 | 0.5019 | 0.1619 | -0.09116 |
| **log bta** | = 0.000 | = 0.000 | = 0.000 | = 0.000 | 0.05082 | 0.04607 | 0.276 | 0.3148 |
| **alpha * beta** | 2.108 | 1.926 | 2.565 | 0.3107 | 4.172 | 3.532 | 2.741 | 1.673 |
